# Selection-free CRISPR-Cas9 editing protocol for distant *Dictyostelid* species

**DOI:** 10.1101/2025.03.23.644600

**Authors:** Mireia Garriga-Canut, Nikki Cannon, Matt Benton, Andrea Zanon, Samuel T. Horsfield, Jacob Scheurich, Kim Remans, John Lees, Alexandre Paix, Jordi van Gestel

## Abstract

*Dictyostelids* are a species-rich clade of cellular slime molds that are widely found in soils and have been studied for over a century. Most research focusses on *Dictyostelium discoideum*, which – due to its ease of culturing and genetic tractability – has been adopted as a model species in the fields of developmental biology, cell biology and microbiology. Over decades, genome editing methods in *D. discoideum* have steadily improved but remain relatively time-consuming and limited in scope, effective in a few species only. Here, we introduce a CRISPR-Cas9 editing protocol that is cloning-free, selection-free, highly-efficient, and effective across *Dictyostelid* species. After optimizing our protocol in *D. discoideum*, we obtained knock-out efficiencies of ∼80% and knock-in efficiencies of ∼30% without antibiotic selection. Efficiencies depend on template concentrations, insertion sizes, homology arms and target sites. Since our protocol is selection-free, we can isolate mutants as soon as one day post-transfection, vastly expediting the generation of knock-outs, fusion proteins and expression reporters. Our protocol also makes it possible to generate several knock-in mutations simultaneously in the same cells. Boosted by cell-sorting and fluorescent microscopy, we could readily apply our CRISPR-Cas9 editing protocol to phylogenetically distant *Dictyostelid* species, which diverged hundreds of millions of years ago and have never been genome edited before. Our protocol therefore opens the door to performing broad-scale genetic interrogations across *Dicyostelids*.

## Introduction

*Dictyostelids*, commonly known as social amoebae or cellular slime molds, are a ubiquitous group of soil amoebae that are known for their biphasic life cycle where cells alternate between a solitary feeding phase – phagocytizing soil bacteria – and a collective dispersal phase – aggregating into spore-bearing fruiting bodies (Raper 1984; Bonner 2009). Although there are over a hundred described *Dictyostelid* species (Schaap et al. 2006; Baldauf et al. 2018; Sheikh et al. 2018), only one of them, *Dictyostelium discoideum* (Raper 1935), has broadly been adopted as a ‘go-to’ model species in several research fields: in cell biology, *D. discoideum* is, for example, used to study chemotaxis (Van Haastert and Devreotes 2004), pinocytosis (Vines and King 2019; Kay et al. 2024) and phagocytosis (Cosson and Soldati 2008; Jauslin et al. 2021); in developmental biology, it is used to study pattern formation (Tomchik and Devreotes 1981; Gregor et al. 2010), slug migration (Francis 1964), and morphogenesis (Loomis 2015); and, in microbiology, it is used to explore microbial cooperation (Medina et al. 2019; Ostrowski 2019), predation (Tsuchiya et al. 1972; Stewart et al. 2022; Steele et al. 2023), pathogenicity (Steinert and Heuner 2005; Cardenal-Muñoz et al. 2018) and other ecological interactions (Kuserk 1980; Kessin et al. 1996; Bonner and Lamont 2005; Landolt et al. 2006). *D. discoideum* is easy to culture and stock, grows rapidly and is genetically tractable (Eichinger et al. 2005; Fey et al. 2007, 2013). In contrast to wild isolates, lab derivatives of *D. discoideum* can furthermore be grown axenically (Sussman and Sussman 1967), which simplifies experimental practices.

Since the generation of the first knock-out mutant in 1987 (De Lozanne and Spudich 1987; Loomis 1987; Witke et al. 1987), genome editing methods in *D. discoideum* have strongly improved (Paschke et al. 2018, 2019) with, among others, the development of selectable markers (Sutoh 1993), restriction enzyme-mediated DNA integration (REMI) (Kuspa and Loomis 1992; Gruenheit et al. 2021), PCR-mediated gene disruptions (Kuwayama et al. 2002), and a synthetic biology toolbox (Kundert et al. 2020). More recently, Muromoto and colleagues pioneered the first CRISPR-Cas9 editing protocol in *D. discoideum* (Sekine et al. 2018; Iriki et al. 2019; Muramoto et al. 2019; Yamashita et al. 2021), marking another major leap in genome editing. Their protocol is based on the expression of an all-in-one CRISPR plasmid, encoding *Streptococcus pyogenes* Cas9 (SpyCas9) and a single guide RNA (sgRNA), making genome editing cost-effective, easy to implement and accessible for library construction (Sekine et al. 2018). Not surprisingly, CRISPR-Cas9 editing is now widely adopted by the field (e.g., Jauslin et al. 2021) and led to the generation of the first pooled genome-wide CRISPR-based knock-out library in *D. discoideum* (Ogasawara et al. 2022), complementing previous efforts using REMI-seq (Gruenheit et al. 2021; Stewart et al. 2022).

Despite the enormous progress in genome editing methods, several limitations remain. First, it remains relatively time-consuming to generate targeted knock-out or knock-in mutants in *D. discoideum*: it can easily take a few weeks from producing cloning vectors to isolating mutant cells. This can limit the scope of mutants that can be compared in any one study, especially when comparing mutants individually (i.e., arrayed format). Second, most genome editing protocols are specifically optimized for *D. discoideum* and, therefore, cannot be readily applied to other *Dictyostelid* species. In fact, most genome editing protocols exclusively focus on axenic strains, including the existing CRISPR-Cas9 editing protocols. This constrains genetic analyses across *Dictyostelid* species, which contrasts the long-standing tradition of performing broad-scale phenotypic comparisons among species, focusing on for example the diversity of multicellular phenotypes (Raper 1984; Schaap et al. 2006; Schilde et al. 2014; Kuzdzal-Fick et al. 2023). To be effective across species, a genome editing protocol should ideally be (1) selection-free, to preclude the need for antibiotic markers, since *Dictyostelids* show strong differences in their resistance to antibiotics both across growth conditions and between species (Paschke et al. 2018; Narita et al. 2020; Zhu et al. 2023); (2) effective under non-axenic growth conditions, since most *Dictyostelid* species cannot be grown axenically; and (3) require little to no optimization, such as the optimization of cloning vectors, to easily target a broad range of *Dictyostelid* species.

In this study, we present a CRISPR-Cas9 editing protocol that is both time efficient and effective across species: our method is cloning-free, selection-free, effective in non-axenic growth conditions and readily applicable to distant *Dictyostelid* species. Building on CRISPR methods developed in other eukaryotes (Doudna and Charpentier 2014; Paix et al. 2017*a*, 2017*b*), we directly deliver the ribonucleoprotein (RNP) complex to *Dictyostelid* cells together with a homology-directed repair (HDR) template using electroporation. This approach significantly improves knock-out and knock-in efficiencies, being on par with the plasmid-based CRISPR-Cas9 protocols, but without the need for cloning or antibiotic selection. Boosted by single-cell sorting, we can furthermore isolate knock-out or knock-in mutants as soon as one day post-transfection, which strongly accelerates the rate at which mutants can be generated. We first optimized our protocol for *D. discoideum* and then showcased its impact by generating fluorescent knock-in mutants in six widely diverse *Dictyostelid* species, three of which have never been genetically modified before. Our CRISPR-Cas9 editing protocol therefore significantly expands the genetic toolkit for phylogenetically distant *Dictyostelids*, while simplifying and expediting genome editing in *D. discoideum*.

## Results

### RNP complex mediates high knock-out efficiencies in *D. discoideum*

Genome editing with CRISPR-Cas9 is mediated by the SpyCas9 endonuclease that, guided by RNA, causes a double-strand break that triggers a cell’s DNA repair machinery. Repair either leads to error-prone non-homologous end-joining (NHEJ) or HDR. HDR requires a repair template and therefore allows for specific gene edits. SpyCas9 forms a complex with CRISPR RNA (crRNA) and trans-activating crRNA (tracrRNA), where specificity results from the complementarity between the 20nt crRNA spacer sequence and DNA target (Jinek et al. 2012). In many studies, crRNA and tracrRNA are replaced by a single chimeric guide RNA (sgRNA). The DNA target must be flanked downstream by a short protospacer adjacent motif (PAM) sequence that is recognized by SpyCas9 (‘NGG’). Previous work has shown that CRISPR-Cas9 editing is more efficient when cells are transfected with the RNP complex, as opposed to a CRISPR plasmid, allowing for selection-free genome editing (Kim et al. 2014). With the aim of efficiently editing *D. discoideum*, and the final goal of editing distant *Dictyostelid* species, we started by testing genome editing efficiencies with the RNP complex in *D. discoideum* AX2 grown under standard axenic conditions.

To quantify knock-out efficiencies, we first generated *D. discoideum* AX2 *act5*::*mCherry* (following Paschke et al., 2018), which constitutively expresses *mCherry* under the control of the endogenous *act5* promoter. We transfected this strain with an RNP complex targeting the 5’-end of the *mCherry* CDS (see Figure 1A, Methods and Text S2) and quantified the fraction of cells with an *mCherry* knock-out (non-fluorescent cells) two days post-transfection using flow cytometry (Figure S1). As *D. discoideum* is haploid, the fraction of non-fluorescent cells is a direct readout for the fraction of knock-out mutants. Sanger sequencing of isolated clones confirmed that cells without fluorescent signal had obtained an indel mutation at the *mCherry* target site (Figure S2A-B). To trigger HDR, we also co-transfected cells with single-stranded donor oligos that serve as a template for homologous recombination and cause a knock-out by introducing a frameshift mutation and stop codon. We varied the concentration of the donor oligo (0-4.8µM), length of homology arms (14, 28, 56nt) and insertion sizes (1, 4, 37nt) (Figure 1B-D). In the absence of donor oligo, we obtained knock-out efficiencies of ∼5%, but with co-transfection of donor oligo, these efficiencies substantially increased. Knock-out efficiencies improved with higher oligo concentrations, longer homology arms, and larger insertion sizes. The highest knock-out efficiencies of ∼80% were obtained for 37bp insertions using homology arms of either 28 or 56bp (Figure 1C). The 37bp insertion includes an HA-tag sequence, frameshift mutation and stop codon (Table S6). Two-fold lower efficiencies were obtained with 1bp (44±1.9%) and 4bp insertions (41±4.6%), both of which were designed to include a frameshift mutation and stop codon (Figure 1D). Using both Sanger sequencing (Figure S2C-E) and Nanopore sequencing (Figure S3 and Table S3), we confirmed that HDR-mediated CRISPR-Cas9 editing resulted in a scarless integration of the donor oligo at the expected target site, without any off-target integrations. Control experiments where cells were transfected with SpyCas9 and tracrRNA only (i.e., without crRNA) showed no mutations in *mCherry* (Figure S2F and Table S3). Knock-out efficiencies also differed depending on the DNA target site: two of the three tested CRISPR-Cas9 targets in *mCherry* gave high knock-out efficiencies, whereas one gave a four-fold lower efficiency (Figure S4A). Finally, we also compared different SpyCas9 proteins, including wild-type (WT) and high-fidelity variants, and different SpyCas9 purification methods (see Methods), which showed that the best knock-out efficiencies were obtained with ultrapure WT SpyCas9 (see Figure S4B and Methods). The best-performing commercially available SpyCas9 proteins generated about two-fold lower knock-out efficiencies, presumably due to their lower purity. Given that knockout efficiencies peaked for donor oligos with a 37bp insertion and 28bp flanking homology arms, we used these oligos at a concentration of 2.4µM along with ultrapure WT SpyCas9 for all knock-out experiments described below, unless specified otherwise.

**Figure 1.**
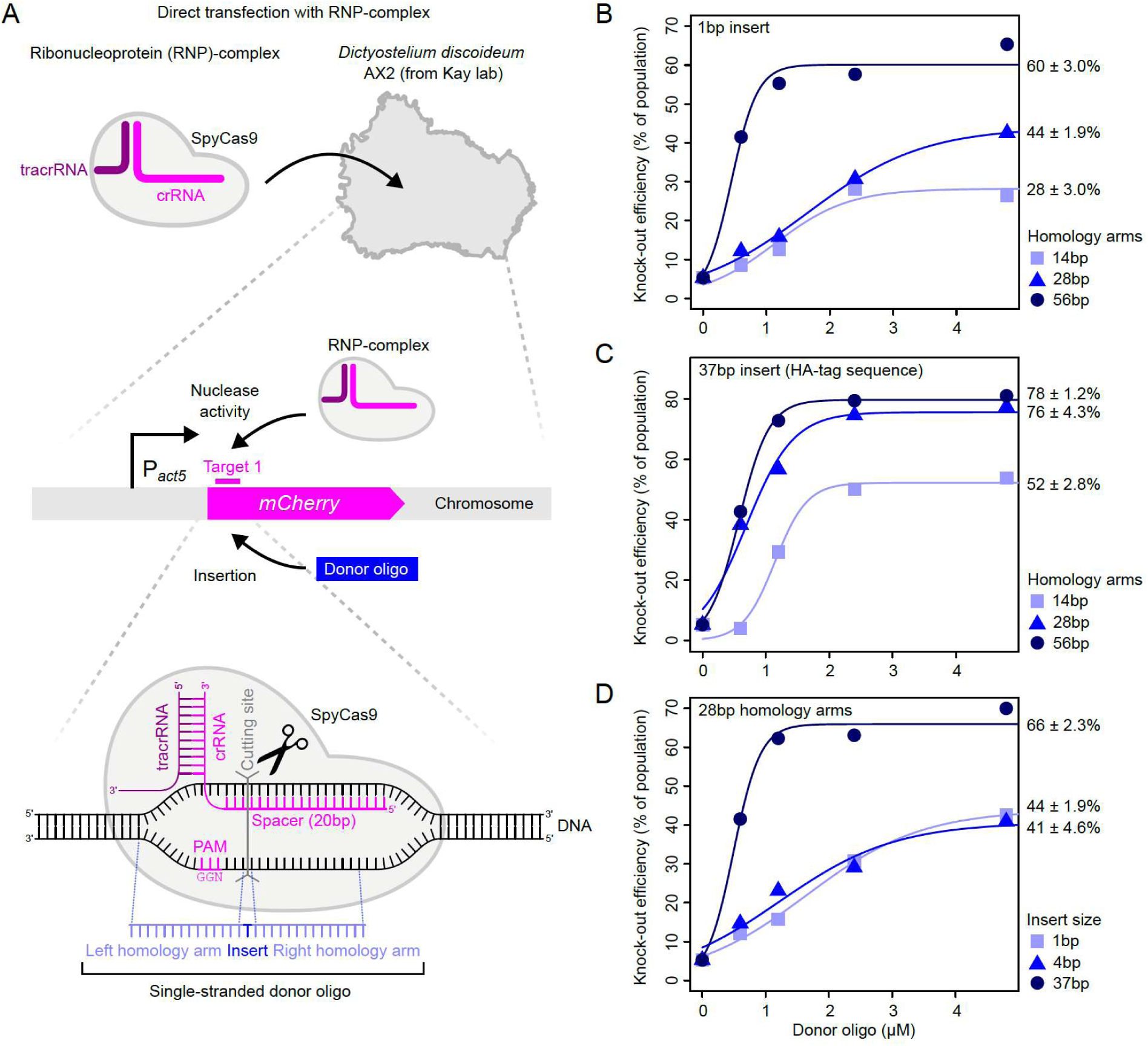
High knock-out efficiency through donor oligo mediated CRISPR-Cas9 editing. (A) Schematic depiction of CRISPR-Cas9 editing with RNP complex consisting of *Streptococcus pyogenes* Cas9 (SpyCas9), CRISPR RNA (crRNA) and trans-activating crRNA (tracrRNA). Single-stranded donor oligo encodes for insertion sequence as well as flanking homology arms mediating homology-directed repair (HDR). Knock-outs are produced by targeting amino acid 9 of *mCherry* CDS of *D. discoideum* AX2 *act5*::*mCherry*. The same target is used for figures hereafter, unless noted otherwise. (B) Knock-out efficiencies for 1bp insertion (frameshift mutation) and 14, 28 and 56bp homology arms. (C) Knock-out efficiencies for 37bp insertion (HA-tag sequence, frameshift mutation and stop codon) with 14, 28, and 56bp homology arms. (D) Knock-out efficiencies for 1, 4 (stop codon and frameshift mutation) or 37bp insertion with 28bp homology arms. Percentages in (B), (C) and (D) show estimated maximum knock-out efficiencies and standard errors based on sigmoidal fits. As there can be considerable variation between cell batches (Figure S4A), the comparisons of knock-out efficiencies within each panel are based on a single batch of cells. The same applies to figure panels with sigmoidal fits hereafter. See Data S1 for data.

### RNP complex mediates high knock-in efficiencies in *D. discoideum*

Next, we examined whether the RNP complex could also boost knock-in efficiencies. To this end, we co-transfected cells with a donor PCR product encoding *mNeonGreen-P2A* and target-specific homology arms (34/37bp) for generating an in-frame knock-in mutation targeting the 5’end of *mCherry* (Methods and Figure 2A). P2A induces ribosomal skipping during translation and can thus be used to express multiple proteins from a single promoter (Zhu et al. 2023). With this construct, we quantified both knock-in and collateral knock-out efficiencies (Figure 2): when a double-strand break in *mCherry* results in HDR there is a gain in green fluorescent signal and when it results in NHEJ there is a loss of red fluorescent signal. In the absence of PCR template, we get ∼5% knock-out efficiencies, as observed above (Figure 2B-C and S5). With increasing concentrations of PCR template, we get an HDR-mediated knock-in efficiency of up to 27±1.5% and a collateral NHEJ-mediated knock-out efficiency of 29±1.9%. In other words, approximately one third of the cells are unaffected, one third have a knock-out mutation, and one third have the desired knock-in mutation. The striking increase in collateral knock-out mutants with increasing template concentrations is likely the consequence of a carrier effect, where the donor PCR product either facilitates the transfection and/or stabilization of the RNP complex or stimulates cell repair mechanisms. In support, when using non-specific oligos or electroporation enhancer, in combination with donor oligos, we also observed higher knock-out efficiencies (Figure S6). Similarly, electroporation enhancer in combination with donor PCR product also slightly increases collateral knock-out efficiencies (Figure S7A-B). Other additives, such as homologous-directed recombination enhancer from Integrated DNA Technologies (IDT), did not have an effect on either knock-out or knock-in efficiencies (Figure S7C-D). As high knock-in efficiencies are obtained with short homology arms (∼30bp, see also Paix et al. 2017*a*), HDR templates can easily be generated by PCR using primers with overhangs that match the flanking sequence of the CRISPR-Cas9 cutting site, making our protocol effectively cloning-free. Our CRISPR-Cas9 editing protocol thus strongly boosts genome editing efficiencies in *D. discoideum*, resulting in the highest-ever recorded knock-out and knock-in efficiencies without selective marker.

**Figure 2.**
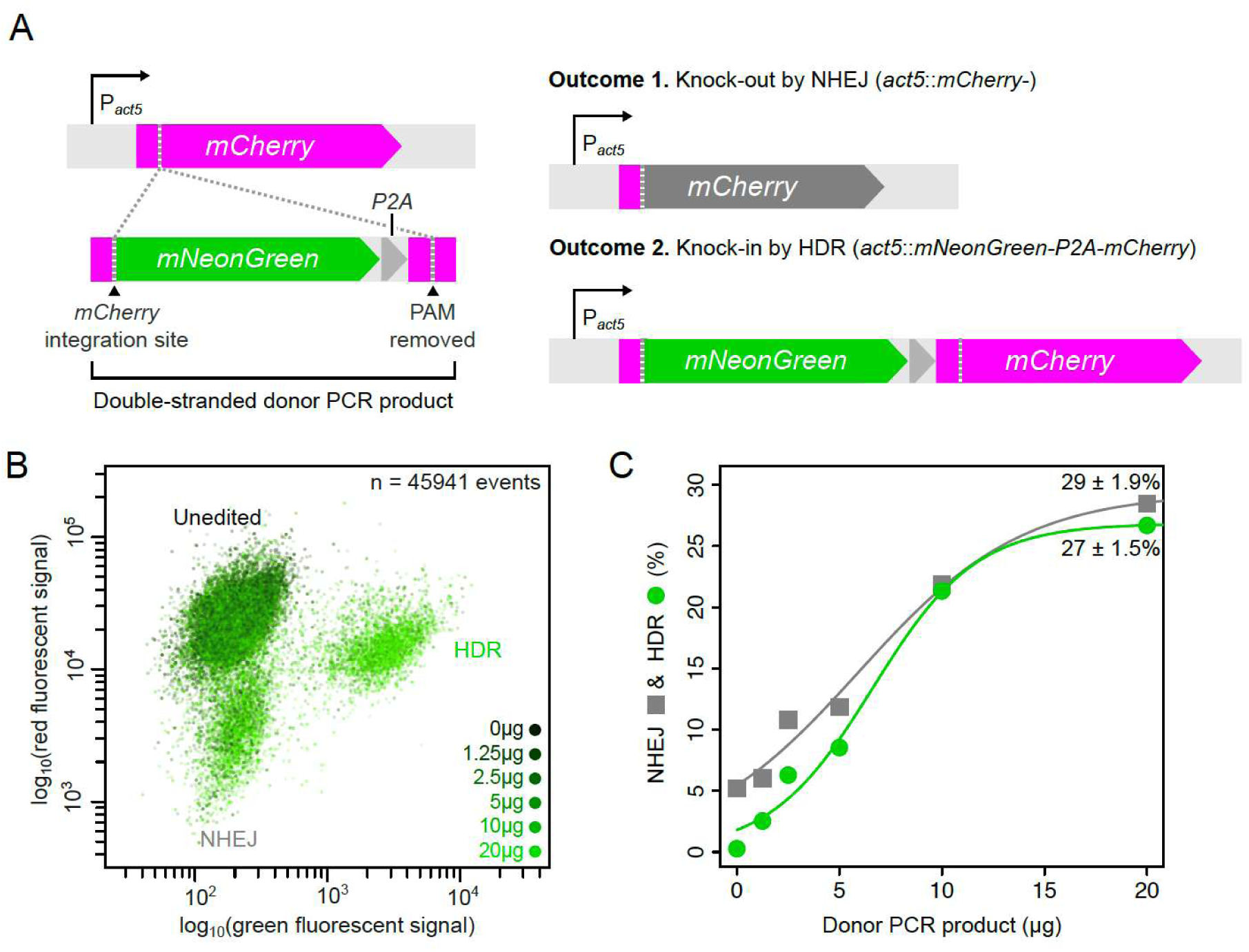
HDR and NHEJ probabilities for CRISPR-Cas9 mediated knock-in of donor PCR product. (A) Schematic of CRISPR-Cas9 editing protocol where *mNeonGreen-P2A* is knocked into the *mCherry* locus, allowing for both green and red fluorescence expression from a single *act5* promoter. Two possible outcomes of CRISPR-Cas9 editing are either (1) non-homologous end-joining (NHEJ) causing an indel in *mCherry*, leading to a knock-out by frameshift mutation, or (2) HDR leading to co-expression of *mNeonGreen* and *mCherry*. The same target and donor PCR product is used for all knock-ins experiments hereafter, unless noted otherwise. (B) Combined flow cytometry results of fluorescent signals of knock-ins produced with 0 to 20µg (0 to 36.12pmol) of donor PCR product. (C) HDR-mediated knock-in and NHEJ-mediated knock-out probabilities with 0-20µg of donor PCR product. Percentages with standard error show estimated maximum knock-in and knock-out efficiencies based on sigmoidal fit to data. See Data S1 for data.

### Genome editing effective in both axenic and non-axenic growth conditions

To target other *Dictyostelid* species, which require bacterial food for growth, our genome editing protocol should also work under non-axenic growth conditions. Traditional genome editing methods often show strong differences between axenic and non-axenic growth conditions, because bacteria can interfere with the efficacy of antibiotic selection (Paschke et al. 2018). We therefore next examined how non-axenic growth conditions affect our selection-free CRISPR-Cas9 protocol. For this we used both *D. discoideum* AX2 and NC4. The axenic strain (AX2) is derived from the non-axenic one (NC4) (Sussman and Sussman 1967; Bloomfield et al. 2015), which means their genomes are largely similar (Bloomfield et al. 2008). We directly compared the efficiencies of generating knock-out mutants using donor oligos and knock-in mutants using donor PCR product under both axenic and non-axenic growth conditions. For non-axenic growth conditions, we cultured *D. discoideum* at different densities of *Escherichia coli* B/r. Although there are somewhat lower knock-out efficiencies in the presence of bacteria, they remained relatively high across the board: 61±5.5% for *D. discoideum* AX2 and 73±1.5% for *D. discoideum* NC4 (Figure 3A-B). We did not observe consistent differences between transfections done at low and high bacterial densities across strains, suggesting that the range of bacterial densities examined here has a minimal impact. Knock-in efficiencies remained exceptionally high under non-axenic growth conditions as well for both *D. discoideum* AX2 and NC4 (Figure 3C), where we again observed a carrier effect (comparison between NHEJ-mediated knock-out efficiencies between Figure 3C and 3D).

**Figure 3.**
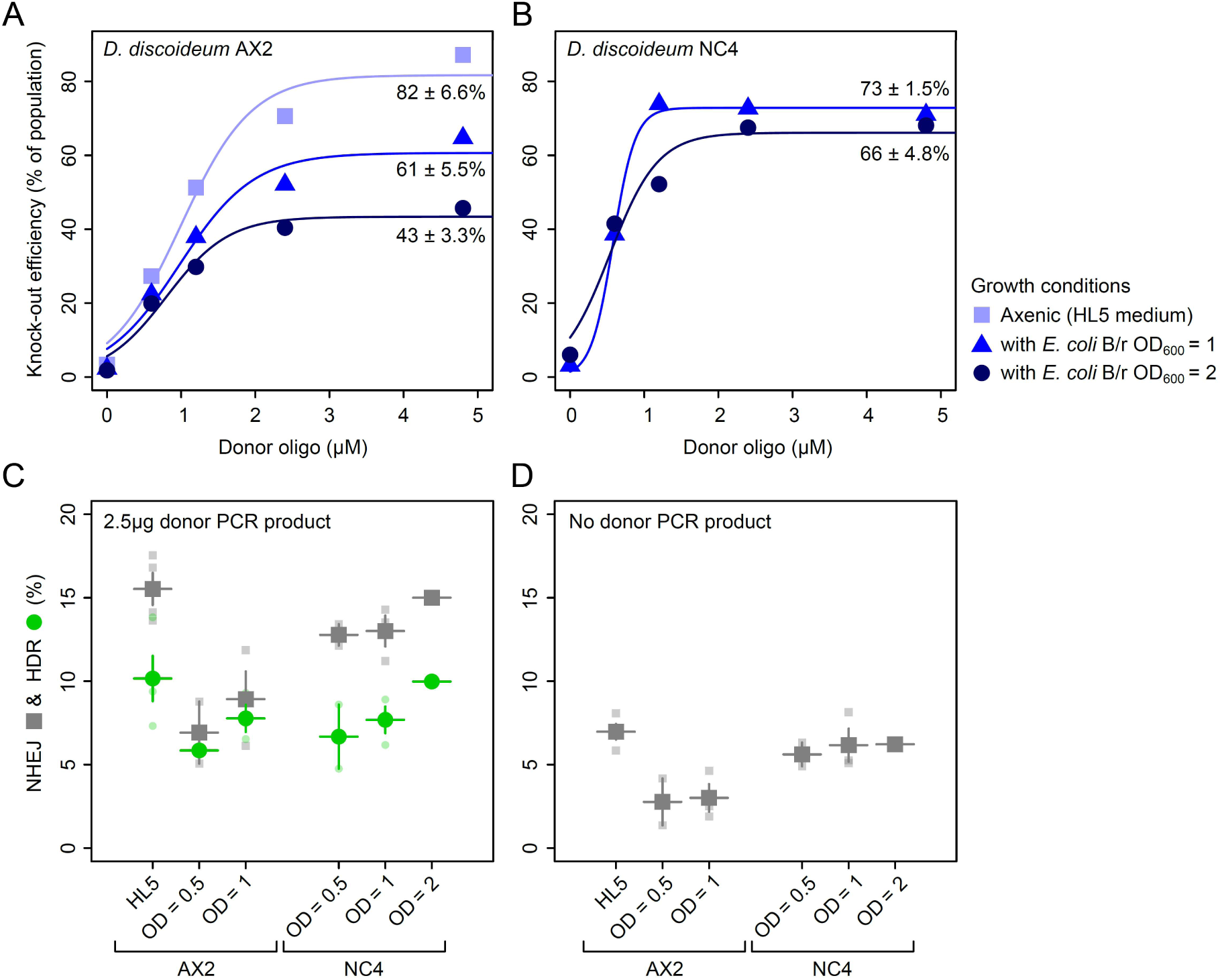
Knock-out and knock-in efficiencies under axenic and non-axenic growth conditions. Knock-out efficiencies targeting *mCherry*, using a donor oligo with 37bp insertion and 28bp homology arms, in (A) *D. discoideum* AX2 *act5*::*mCherry* and (B) *D. discoideum* NC4 *act5*::*mCherry* under axenic growth and non-axenic growth condition (with *E.coli* B/r at different cell densities as measured by OD_600_). (C) HDR and NHEJ probabilities for *D. discoideum* AX2 and NC4 with 2.5µg (4.52pmol) of donor PCR product under axenic and non-axenic growth conditions. Symbols and vertical lines show mean and standard errors respectively (n=2-3). (D) Knock-out efficiencies in absence of donor PCR product. Like observed for axenic growth conditions, there is a carrier effect: collateral NHEJ-mediated knock-out rates in the presence of donor PCR product (C) are higher than in the absence of donor PCR product (D). See Data S1 for data.

### Single-cell sorting expedites isolation of genome-edited clones

Besides genome editing, also the process of isolation requires optimization when aiming to edit distinct *Dictyostelid* species. Under non-axenic growth, mutants are typically isolated using plaque assays: transfected *Dictyostelid* cells are spread across a lawn of bacteria, where they form small plaques, originating from single cells, from which clones can be isolated within days. These clones are subsequently propagated for both genotyping and phenotyping. Although plaque assays are effective, they are often difficult to standardize across *Dictyostelid* species and pose problems when growth-deficient mutants are easily outgrown by WT cells. We therefore explored the possibility of single-cell sorting, where transfected cells are sorted in multi-well plates, as an alternative to plaque assays (Fey et al. 1995; Chen et al. 2007). By sorting, cells immediately go through a single-cell bottleneck without the need to isolate and propagate them in plaque assays, which saves considerable time. Sorting also shields growth defective mutants from WT cells.

To assess sorting accuracy, we first mixed a population of green-and red-fluorescent *D. discoideum* AX2 cells and sorted them according to fluorescent signal into two 24-well plates (Figure 4A-B). Instead of sorting cells into liquid medium (Fey et al. 1995; Chen et al. 2007), we sorted them onto a charcoal agar medium with a lawn of *E. coli* B/r, which we added to each individual well (charcoal reduces background fluorescence). This sorting approach supports both fast and high recovery rates of single cells and has the additional benefit that we can immediate phenotype clones, minimizing downstream genotyping efforts. Already three days after sorting, 35 out of the 48 sorted wells showed robust growth (Figure 4C-D). In only a single well we observed a mixture of green and red cells, while all other wells contained only green or red cells, as confirmed through both microscopy and flow cytometry (Figure 4C-D and S8). This high sorting accuracy (34/35) shows that cell sorting is an effective method for swiftly isolating individual clones, which could readily be applied to other *Dictyostelid* species. Sorting would also make it possible to selectively enrich for fluorescent mutant cells from a pool of transfected cells, which is beneficial when for example constructing fluorescent expression reporters or fusion proteins.

**Figure 4.**
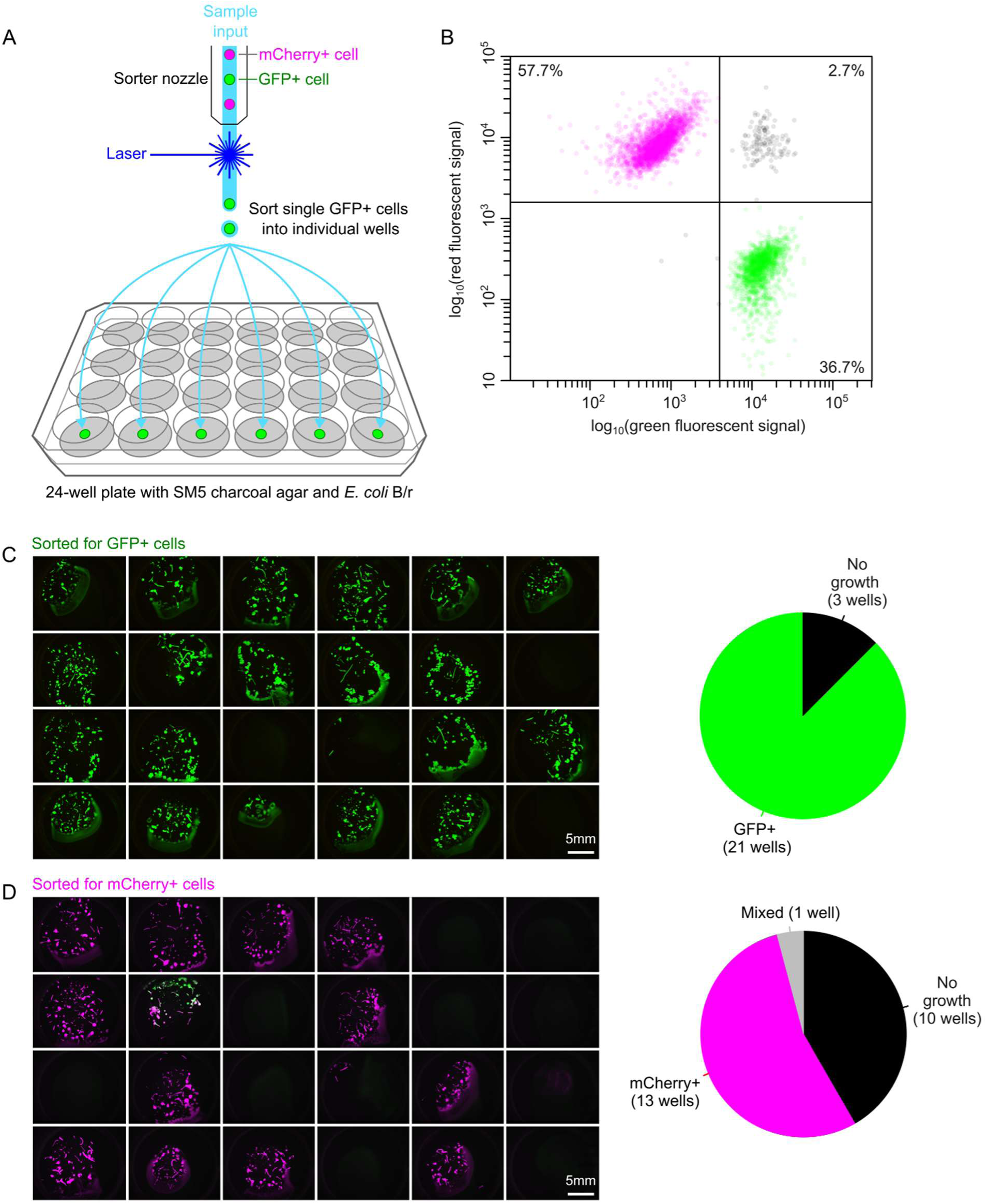
Sorting accuracy of red and green fluorescent *D. discoideum* cells across 24-well plates with charcoal agar and *E. coli* B/r. (A) Schematic depiction of sorting procedure, with the sorter nozzle maintaining a constant flow to sort single cells into wells of a 24-well plate based on their fluorescent intensity. Each well contains 1mL of SM5 charcoal agar with a lawn of *E. coli* B/r. (B) We sorted a mixed population of ∼60% mCherry+ cells and ∼40% GFP*+* cells. From the recorded events, 2.7% showed mixed signal, indicating doublets of green and red cells. (C)-(D) 24-well plates after 4 days of growth for (C) green and (D) red fluorescent (magenta) sorted cells respectively. Pie diagrams show sorting accuracy: wells show either no growth (black), GFP+ (green), mCherry+ (magenta) or mixed (grey) cells. For (C) we obtained zero mixed wells and for (D) one mixed well after sorting. See Figure S8 for detailed comparison to flow cytometer data.

### Targeting endogenous genes for constructing expression reporters and fusion proteins

Leveraging our isolation method, we next targeted endogenous genes for creating expression reporters and fusion proteins. We started by constructing fluorescent expression reporters for *ecmA* and *act5* by introducing in-frame *mNeonGreen-P2A* knock-ins in their native loci, akin to our knock-ins in Figure 2. EcmA is an extracellular matrix protein expressed by prestalk cells during the slug stage (McRobbie et al. 1988; Jermyn and Williams 1991), while Act5 is a major actin protein that is constitutively expressed during growth (Joseph et al. 2008). We applied our sorting protocol to isolate genome edited clones. Since *ecmA* is not expressed during growth, we sorted transfected cells blindly (i.e., single cells were sorted irrespective of their fluorescence signal), while for *act5* we sorted for green-fluorescent cells. All cells were sorted two days post-transfection and imaged a few days later. Within the well, cells form a feeding front, slugs and fruiting bodies, allowing us to directly examine through fluorescent microscopy whether genome editing was successful. For *ecmA*, 15% of wells with growth showed expression of our reporter, while for *act5* all wells with growth had green cells (Figure 5A-B). All genomic edits were confirmed by PCR and there were no indications of gene silencing upon perpetual propagation of individual clones. When profiling slugs, we confirmed that, as previously reported, *ecmA* is expressed in the ∼20% anterior part of slugs (Loomis 1987; Jermyn and Williams 1991), irrespective of slug size, while *act5* was uniformly expressed throughout the slug (Figure 5C-D).

**Figure 5.**
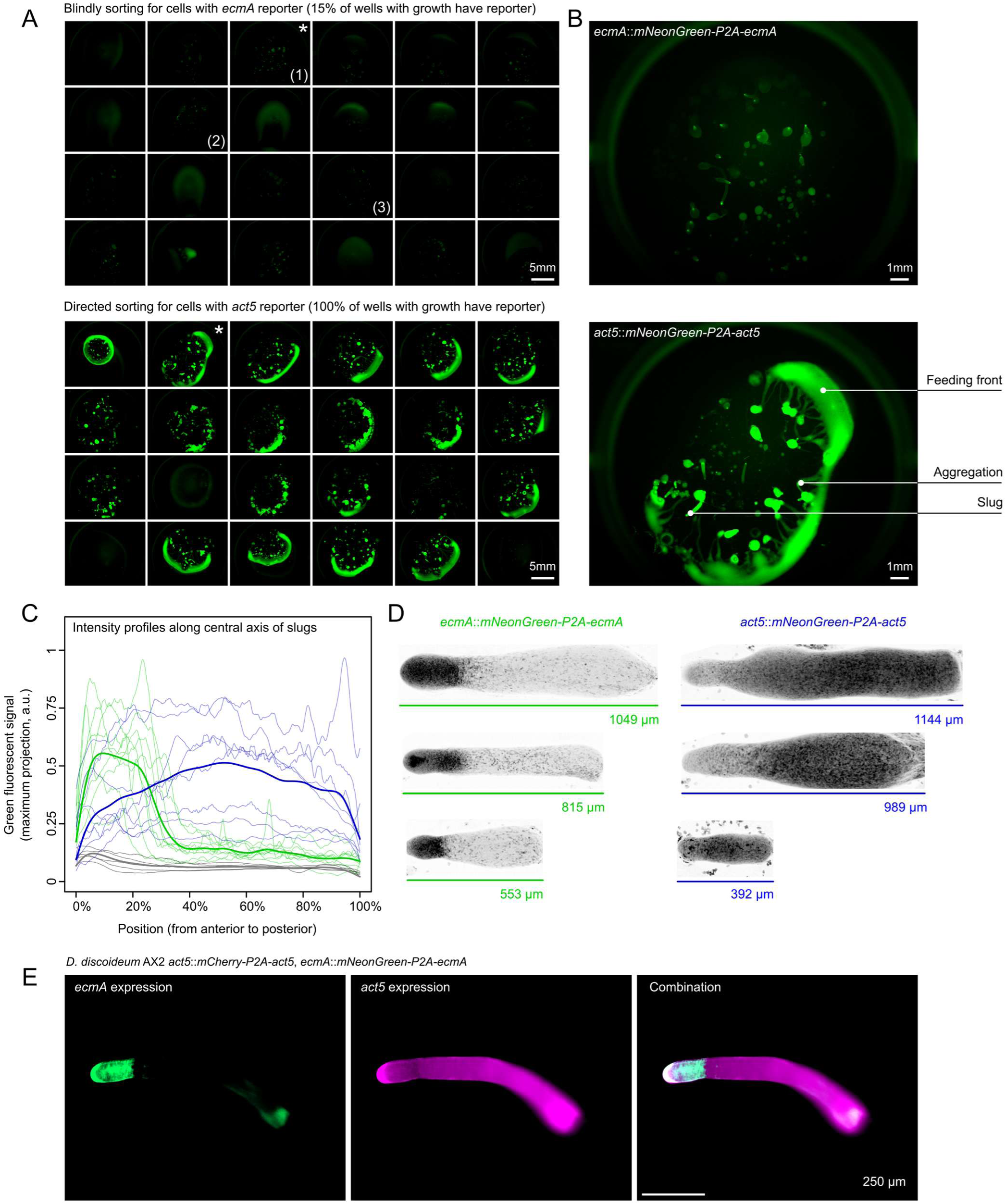
CRISPR-Cas9 generated *ecmA* and *act5* expression reporters. *D. discoideum* AX2 cells were transfected with RNP complex targeting the 27^th^ amino acid of *ecmA* or the first amino acid for *act5* and a donor PCR product encoding for *mNeonGreen-P2A* in frame with *ecmA* or *act5*, respectively. (A) 24-well plates of sorted transfected cells after 4 days of growth for both the *ecmA*-or *act5*-expression reporters (top and bottom panel, respectively). For the *ecmA* reporter, cells were sorted blindly, while for the *act5* reporter, cells were sorted based on green fluorescence signal. Wells with no growth give a diffuse green fluorescence signal due to autofluorescence of the bacteria lawn. For *ecmA*, three of the wells with growth were positive (as indicated by the numbers). For *act5*, all of the wells with growth were positive. (B) Examples of positive wells for *ecmA* and *act5* expression reporters, highlighted by asterisk in (A). Wells show feeding front, aggregation and slug formation. Cells from positive wells were propagated further to SM5 charcoal agar plates with *E. coli* B/r as food source for profiling slugs (charcoal was added to minimize background fluorescence). (C) Expression profiles of slugs, from anterior to posterior, for WT (grey), *ecmA* (green) and *act5* (blue) expression reporters. Transparent lines show maximal projections along central axes of slugs from anterior to posterior, based on a 20µm rolling window. Solid lines show smooth splines across all maximal projections (n=5-9). (D) Example images (with inverse LUT) of slugs for *ecmA* and *act5* expression reporters, confirming that slugs express *ecmA* in the anterior 20% of the slug irrespective of their size (see File S2 and File S3 for image data). (E) Example of double knock-in mutant (*D. discoideum* AX2 *act5*::*mCherry-P2A-act5, ecmA*::*mNeonGreen-P2A-ecmA*), where *act5* and *ecmA* knock-ins were generated simultaneously using, respectively, *mCherry-P2A* and *mNeonGreen-P2A* as templates: (left) *ecmA* expression (green), (middle) *act5* expression (magenta) and (right) composite image. See Figure S9 for associated flow cytometry results.

We also applied our genome editing protocol towards creating endogenous fusion proteins. For this, we targeted *Act5* and *H2Bv3* and created either N-terminus or C-terminus fusion proteins with *mNeonGreen*. H2Bv3 is one of the main histone proteins involved in chromatin structure and thus localizes to the nucleus (Stevense et al. 2011), while Act5 polymerizes near the leading-edge of moving cells (Ishikawa-Ankerhold and Müller-Taubenberger 2019). As for the above experiments, we obtained high knock-in efficiencies, especially when targeting the N-terminus (Figure 6A). Following our expectation, mNeonGreen-H2Bv3 fusion protein localized to the nucleus, while mNeonGreen-Act5 was cytoplasmic (Figure 6B). When quantifying mNeonGreen-Act5 expression along the radial axis of cells (Methods), from their periphery to the center, we observed that – like expected – most Act5 localized near the periphery (Figure 6C-D).

**Figure 6.**
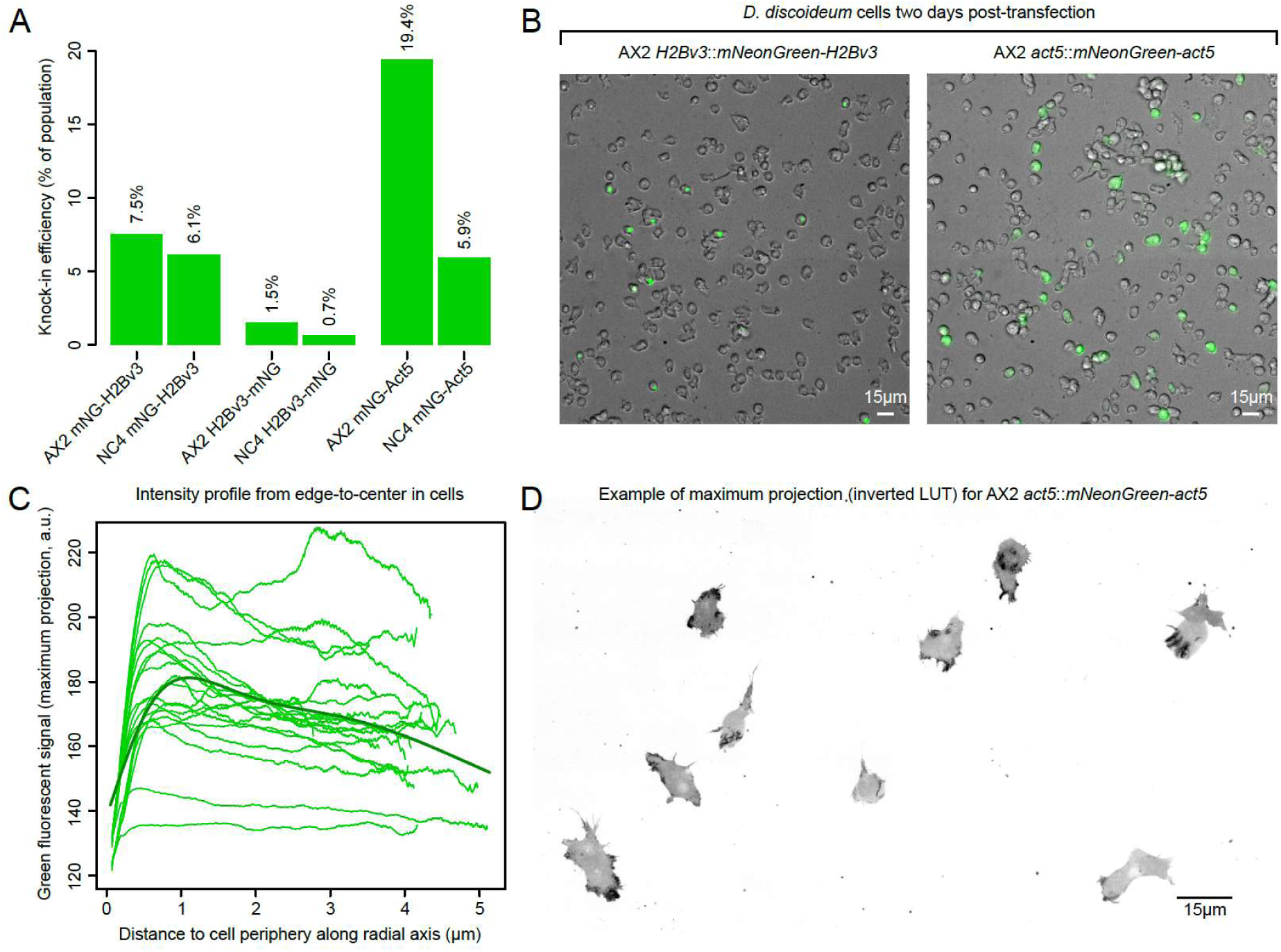
CRISPR-Cas9 generated H2Bv3 and Act5 protein fusions. (A) Knock-in efficiencies for *mNeonGreen*, using 10µg (18.06pmol) donor PCR product, targeting *H2Bv3* and *act5* loci for producing mNeonGreen-H2Bv3, H2Bv3-mNeonGreen and mNeonGreen-Act5 protein fusions in *D. discoideum* AX2 and NC4. (B) Cells two days post-transfection, showing nucleus-localized H2Bv3 protein and cytoplasmic-localized Act5 protein. (C) Expression profiles of mNG-Act5 along radial axis of cells from periphery to center, highlighting enriched localization of mNG-Act5 near the periphery. (D) Example of mNG-Act5 expression in *D. discoideum* cells (see File S4 for image data). mNG=mNeonGreen.

In summary, our CRISPR-Cas9 editing protocol in combination with single-cell sorting makes it possible to target endogenous genes and create expression reporters and fusion proteins within days: cells can be sorted 1-2 days post-transfection and imaged 3-4 days later. Since our protocol is cloning-free, new knock-ins can be generated quickly by simply designing and ordering new crRNAs and donor PCR oligos with homology arm overhangs (i.e., there is no need to re-design a cloning vector), thereby minimizing preparation time. This strongly contrasts with traditional genome editing methods, where it can often take several weeks to construct and examine knock-in mutants (requiring cloning, antibiotic selection and plaquing).

### Genome editing of multiple genes simultaneously

Since CRISPR-Cas9 editing is targeted and does not rely on selective markers, we hypothesized that it should also be possible to create multiple knock-ins simultaneously. To test this, we mixed the RNP complexes targeting both *act5* and *ecmA*, and performed a single transfection with *mCherry-P2A* and *mNeonGreen-P2A* as repair templates to create a double expression reporter for *act5* and *ecmA*, respectively (Figure S9A-C). From the 36 wells we imaged after sorting for red-fluorescent cells, we detected double knock-in mutants in six wells (Figure 5E). Given this high knock-in efficiency (17%), the maximum number of fluorescent reporters that can be generated at once is likely to be even higher, especially when these reporters are already expressed during sorting and allow for combinatorial gating.

We also examined whether we could generate both a knock-in and knock-out mutation simultaneously in a combinatorial transfection. Using *D. discoideum* AX2 *act5*::*mCherry*, we specifically knocked in *mNeonGreen* at the N-terminus of *H2Bv3* to create an endogenous fusion protein and simultaneously knocked out *mCherry* (without donor oligo to only allow for NHEJ-mediated knock-out). Like for the double knock-in experiment (Figure 5E and S9C), we observed many cells with both a knock-in and knock-out mutation (Figure S9D-F). Double mutants were even enriched: cells with a knock-in had a significantly higher probability of also having a knock-out mutation (22% instead of 10%) and, conversely, cells with a knock-out had a significantly higher probability of also having a knock-in mutation (18% instead of 8%) (Fisher’s Exact Test; *p* < 10^-10^, odds ratio=2.3 with [1.8-2.8] 95%-confidence intervals, n=7223 cells; Figure S9D-F). Combinatorial transfections can therefore be used to enrich for knock-out mutants without immediate phenotype by sorting for parallel knock-ins that do have a phenotype (e.g., protein fusion or expression reporter) (see also Arribere et al. 2014).

As combinatorial transfections require no additional preparation, our CRISPR-Cas9 editing protocol strongly simplifies the way multiple genome edits can be generated compared to previous methods, where knock-ins are often produced sequentially by reusing selection markers through Cre recombinase-mediated removal (Faix et al. 2004; Linkner et al. 2012) or where a combinatorial CRISPR-Cas9 plasmid needs to be generated before (Sekine et al. 2018).

### Genome editing effective in many *Dictyostelid* species

With our optimized editing and sorting protocol in hand, we next targeted other *Dictyostelid* species. To identify CRISPR-Cas9 targets and minimize potential off-target hits, we ideally need genomes with high-quality assemblies. To date, there are only 15 *Dictyostelid* reference genomes, of which two have chromosome-level assemblies (*D. discoideum* and *Dictyostelium firmibasis*). From all available genomes, including those with contig-level assemblies, we selected 8 representative species with *act5* homologs that could be targeted (Figure S10): *Dictyostelium firmibasis*, *Dictyostelium purpureum*, *Polysphondylium violaceum*, *Tieghemostelium lacteum*, *Speleostelium caveatum*, *Cavendaria fasciculatum*, *Heterostelium pallidum* and *Acytostelium subglobosum.* For each species, we directly used our CRISPR-Cas9 editing protocol, without any further optimization, to knock-in *mNeonGreen-P2A* in the N-terminus of the *act5* homolog. Two days post-transfection we sorted transfected cells by selecting for green-fluorescent cells. From the 8 species, we detected green cells in 6 species, from which we could isolate 5 species (Figure 7A-B). To our knowledge, for three of those species, *T. lacteum, D. purpureum and D. firmibasis,* targeted knock-ins have never been generated before, while for the remaining two (*P. violaceum* and *H. pallidum*) knock-ins were previously generated by integrating antibiotic markers (Fey et al. 1995; Narita et al. 2020).

**Figure 7.**
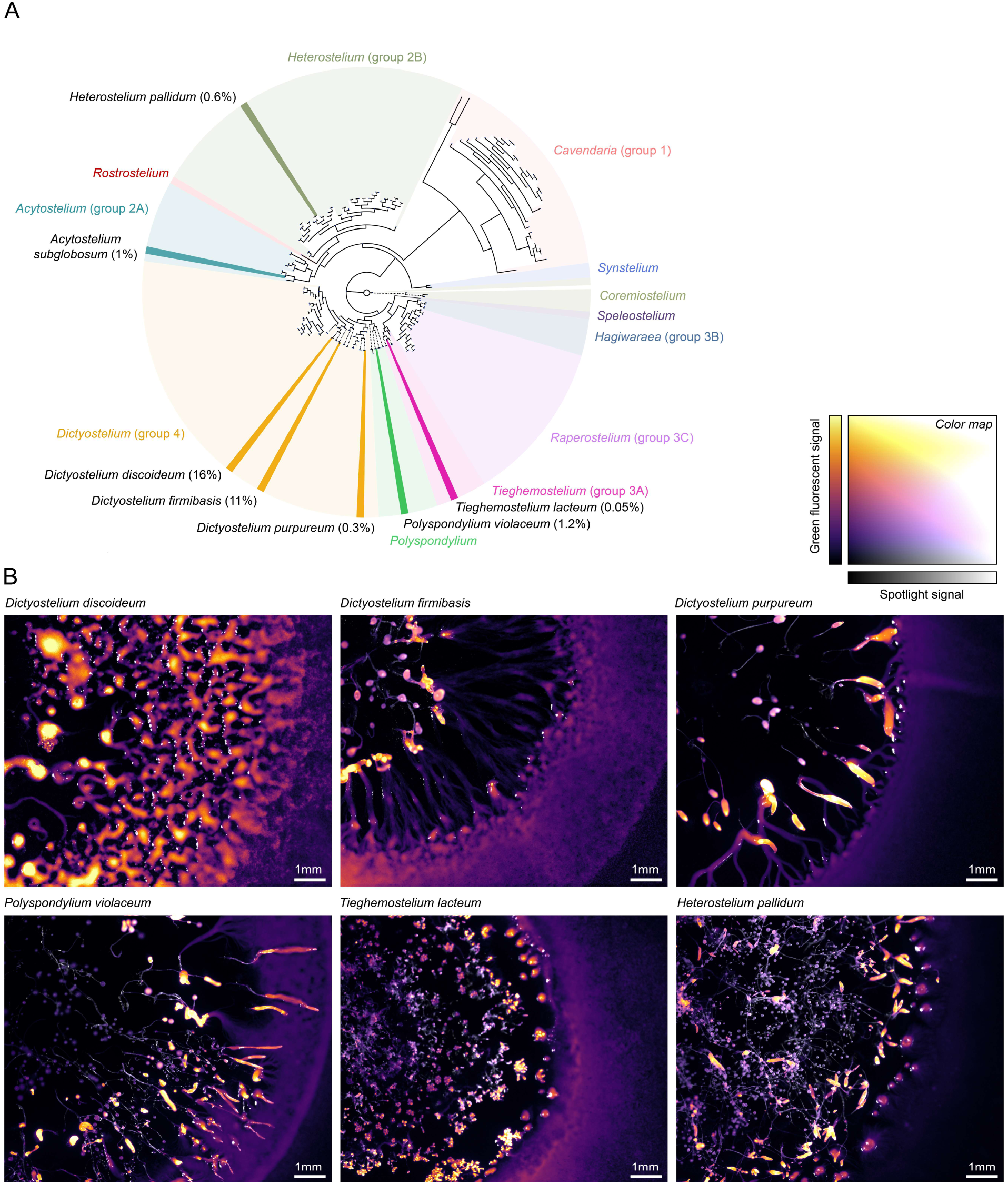
CRISPR-Cas9 editing of distant *Dictyostelid* species. (A) Phylogenetic tree of *Dictyostelids*, based on multiple sequence alignment of 18S rDNA (Sheikh et al. 2018), with highlighted *Dictyostelid* species that were successfully edited (Families are shown in different colors; see Figure S10 for detailed phylogeny). Percentages show fraction of GFP+ cells after knock-in of *mNeonGreen-P2A* in *act5* homolog by CRISPR-Cas9 editing (i.e., knock-in efficiency). (B) Feeding front for all successfully edited and isolated *Dictyostelid* species: *D. discoideum*, *D. firmibasis*, *D. purpureum*, *P. violaceum*, *T. lacteum* and *H. pallidum*. *A. subglobosum* was successfully edited, but could not be isolated, likely due to a growth defect. Inferno color scale shows green fluorescence, visualizing the feeding front (falsely colored purple) and aggregation (falsely colored yellow), and grey scale shows spotlight. For images gamma-scaling was applied (see Methods). For complete time-lapse data see File S5 and S6.

For all species, knock-in efficiencies were much lower than in *D. discoideum* (16%), ranging from 0.05% to 11%. Lower efficiencies could result from reduced transfection efficiencies under our electroporation settings, which were optimized for *D. discoideum* (see Text S2) (Narita et al. 2020). For instance, using our electroporation settings, we noticed that most *Speleostelium caveatum* cells did not survive electroporation. Lower efficiencies may also result from lower genome qualities, which can cause errors in designing crRNAs and homology arms and make it difficult to rule out potentially deleterious off-target hits. In support, the *Dictyostelid* species with the best genome assembly, *D. firmibasis*, produced the closest knock-in efficiency (∼11%) compared to *D. discoideum*. It could also be that *act5* homologs in other species do not function as a safe locus and lead to growth defects when knocking in *mNeonGreen-P2A*. For instance, in *A. subglobosum*, we observed green fluorescent cells during sorting, but never managed to isolate and propagate those cells afterwards. For the other *Dictyostelid* species we did not observe any noticeable phenotypic defects (Figure 7B): WT and fluorescent knock-in strains showed similar growth and aggregation dynamics when grown on *E. coli* B/r (see time-lapse movies in Files S5 and S6).

In conclusion, despite the lower knock-in efficiencies, our CRISPR-Cas9 editing and isolation protocol could readily be applied to distant *Dictyostelid* species, even those that have never been edited before. This strongly expands the genetic toolbox for modifying *Dictyostelid* species. With an increasing number of high-quality genomes becoming available, we expect that many more *Dictyostelid* species can be targeted soon as well, making it possible to perform broad-scale genetic interrogations across *Dictyostelid* families.

## Discussion

In this study, we establish a selection-free CRISPR-Cas9 editing and isolation protocol for *Dictyostelids* that comes with several major benefits. Compared to existing protocols, our CRISPR-Cas9 editing protocol strongly expedites the construction of individual mutants in *D. discoideum* for both axenic and non-axenic strains: mutants can be isolated as soon as one day post-transfection and stocked a few days later. For knock-out mutants generated with donor oligos, our protocol gives ∼80% efficiencies, and for knock-ins with large gene fragments (∼1kb) these are close to 30%. When constructing fluorescent expression reporters or fusion proteins, mutants can be sorted directly based on fluorescent signal, minimizing any downstream validation efforts. Sorting could also promote the isolation of larger knock-in constructs (>1kb), which are often associated with lower knock-in efficiencies. By sorting onto agar medium, we also expedite the isolation of strains with expression reporters and protein fusions that are expressed after growth, by screening sorted plates during development (e.g., during aggregation, slug formation, and fruiting) using fluorescent microscopy. Finally, since our protocol is selection-free, we can easily generate several knock-in or knock-out mutations simultaneously, targeting different loci across the genome, which for example makes it possible to construct double expression reporters and target proteins with redundant functions (e.g., when knocking out homologs; provided that target sites are sufficiently distinct).

Our protocol could promote high-content screens in *D. discoideum*, where proteins from the same pathway or protein family are targeted in arrayed knock-out libraries (Bock et al. 2022), thereby complementing pooled and/or non-targeted mutant screens (Gruenheit et al. 2021; Ogasawara et al. 2022; Stewart et al. 2022). Both donor oligos and crRNAs can be ordered in bulk, using multi-well plates, making it possible to create tens of mutants in parallel. By creating compact and targeted mutant libraries (Bock et al. 2022), one can systematically investigate molecular pathways and protein families. *D. discoideum* could for example be used for high-content screens targeting endocytosis (Vines and King 2019), phagocytosis (Cosson and Soldati 2008; Jauslin et al. 2021) and chemotaxis (Van Haastert and Devreotes 2004). *D. discoideum* also shows strong enrichment of important protein families (Eichinger et al. 2005) that can be targeted as well, including polyketide synthases (Zucko et al. 2007), ABC transporters (Miranda et al. 2013), actin-binding proteins (Joseph et al. 2008), G-protein coupled receptors (Hall et al. 2022), and tyrosine kinases (Kin et al. 2023). By creating arrayed libraries with tens of knock-out mutants within the same background strain, it becomes possible to systematically interrogate how proteins affect cellular phenotypes, growth and development.

When targeting a single integration site, our CRISPR-Cas9 editing protocol also promotes the construction of pooled knock-in libraries with hundreds, if not thousands, of mutants. This makes it possible to generate large-scale gain-of-function, protein-fusion and expression libraries within a single safe locus (e.g., *act5*). Although plasmid-based expression libraries and screening methods for gain-of-function mutants in *D. discoideum* already exist (Robinson and Spudich 2000; Li et al. 2016), the ability to generate knock-in libraries has been limited by low knock-in efficiencies. With a knock-in efficiency of ∼30%, our CRISPR-Cas9 editing protocol makes it possible to generate ∼10^6^ knock-in mutants from a single transfection only. When these mutants are also barcoded, large-scale gain-of-function screens can be performed across a wide range of culturing conditions with minimal expenses. Similar gain-of-function libraries have been generated in a broad range of bacterial (Urtecho et al. 2023; Huang et al. 2024) and eukaryotic species already (Jones et al. 2008; Fuqua et al. 2020). Knock-in libraries can also be used for deep mutational scanning studies to investigate the function of a specific protein. One could for example systematically introduce amino acid substitutions along a protein or within a specific protein domain (Papkou et al. 2019; Jones et al. 2020). Given the rich biology of *D. discoideum* and ease of high-throughput culturing, deep mutational scanning methods can be a powerful tool for interrogation key proteins underlying multicellular developmental (e.g., adhesion proteins) and cell biology (e.g., phagocytic receptors).

Finally, without modifications, we showed that our CRISPR-Cas9 editing and isolation protocol can be applied to other *Dictyostelid* species, including species that diverged hundreds of millions of years ago and were never genetically modified before. As long as a sufficiently high-quality genome is available and transfection is possible, genome editing should in principle be possible as well. Thus far, only two *Dictyostelid* species have genomes of the highest assembly level, but our protocol was also effective for species with lower quality genome assemblies. With improving long-read sequencing and the rapid expansion of available genome sequences, we expect that our CRISPR-Cas9 editing protocol can soon be applied to tens of *Dictyostelid* species. For some species, transfection protocols might need optimization, but for most species we expect that our protocol would readily work. Our protocol therefore contributes to the increasing number of protists that are accessible through genetics (Faktorová et al. 2020; Nomura et al. 2020; Akella et al. 2021; Combredet and Brunet 2024). By editing other *Dictyostelid* species, while building on decades of genetic research in *D. discoideum* (Loomis 2015) and comparative research across *Dictyostelids* (Raper 1984; Schaap et al. 2006), we can explore how cellular and developmental processes diverged over hundreds of millions of years – making the *Dictyostelids* one of the few phylogenetic groups at which genotypic and phenotypic interrogations can be performed at scale.

## Materials and Methods

### Reagents and strains

A complete list of strains, as well as Dicty Stock Center ID (if applicable), is available in Table S1, S2, S3 and S4. Selected strains will be deposited to Dicty Stock Center and ATCC. For knock-in and knock-out mutants, we also list the crRNAs, donor oligos, donor PCR template and (confirmation) primers in Table S2, S4-S6. crRNA and primer sequences are given in Table S5 and S6. The pMG005 vector (ordered from Geneart) with a codon-optimized *mNeonGreen* and *P2A* peptide sequence (Zhu et al. 2023) is provided in Text S1 and File S1. pMG005 was used as template to create donor PCR products. pMG008 was used as template for the *mCherry-P2A* knock-in for Figure 5E and S9 and is provided in Text S1 and File S1. pMG005 and pMG008 are deposited to Addgene (IDs 237215 and 237216 respectively). Lists of all reagents, buffers and other materials are provided as part of the detailed CRISPR-Cas9 editing protocol in Text S2.

### Strains and culturing conditions

For each experiment, *Dictyostelid* cells were freshly inoculated on SM5-agar plates with *E. coli* B/r from stocks and, for axenic strains, subsequently propagated in HL5+FAB without bacteria before transfection. SM5-agar plates with *E. coli* B/r were prepared in the following way: 50mL LB media was inoculated with *E. coli* B/r cells from a glycerol stock and cultured at 30°C (220rpm) for 16h. *E. coli* B/r cells were harvested by centrifugation (4000xg, 5 min), washed three times in 1mL of KK2-MC buffer (Text S2), and normalized to an OD_600_ = 2. 200µL of normalized *E. coli* B/r cell suspension was spread on SM5-agar plates and dried for 5 min to produce a food lawn. *Dictyostelid* species were directly spotted on two opposite sides of the lawn from glycerol stocks (Table S1) and grown at 22°C for 3-5 days, until large feeding fronts were visible. For axenic strains, the feeding fronts of 4 to 6 plates were harvested in 1mL of HL5, washed 3 times in HL5 and resuspended in 1mL HL5+FAB (Text S2). Cells were counted using a Countess III cell counter (ThermoFisher) and grown for 2-3 days in 50mL of HL5+FAB at a starting concentration of 10^5^ cell/mL at 22°C (180rpm) until they reached exponential phase (0.8-1.2·10^6^ cells/mL). Typically, cells reached exponential growth after 2 days.

### Expression and purification of SpyCas9

Different variants of recombinant SpyCas9 were purified by the Protein Expression and Purification Core (PeP-core) Facility at EMBL using either a quick one-day protocol for regular purity protein extract or an extensive three-day protocol for an ultrapure protein extract. In order to prevent endotoxin contaminations in both the quick and long protocol, Åkta chromatography stations were incubated with 1M NaOH for 4 hours before use, all chromatography columns were rigorously cleaned with 0.5-1M NaOH and all protein purification buffers were prepared using endotoxin-free reagents.

For the quick protocol, pHO4d-Cas9, pET-HiFi SpCas9-NLS-6xHis and pET-FLAG-spCas9-HF1 (Addgene, Catalog # 67881, #207376 and # 126770 respectively, and File S1) encoding WT SpyCas9, HiFi SpyCas9 and HF1 SpyCas9, respectively, were freshly transformed into *Escherichia coli* Rosetta2 (DE3) cells. Precultures were grown overnight at 37°C in LB medium supplemented with 100µg/mL carbenicillin and 34µg/mL chloramphenicol and used to inoculate the large-scale expression cultures. 10mL preculture was added to 1L of TB-FB supplemented with 2mM MgSO4, 0.05% glucose, 1.5% lactose, 100µg/mL carbenicillin and 34µg/mL chloramphenicol. Cultures were grown at 37°C until OD_600_ reached ∼0.6, after which the temperature was reduced to 18°C. After overnight expression at 18°C, the cultures were harvested by centrifugation (30min, 5000xg, 4°C) and pellets were flash-frozen in liquid nitrogen and stored at -80°C until the start of the protein purification step.

The cell pellet was resuspended in cold lysis buffer (20mM Tris-HCl pH 8.0, 750mM NaCl, 20mM imidazole, 10% glycerol and EDTA-free cOmplete protease inhibitors, Roche). Cells were lysed by 5 passages through a microfluidizer, followed by centrifugation (30 min, 140000xg, 4°C). The cleared lysate was loaded onto a 1mL Protino Ni-NTA column (Macherey-Nagel) pre-equilibrated with 20mM Tris-HCl pH 8.0, 250mM NaCl, 20mM imidazole and 10% glycerol. After loading, the Ni-NTA column was washed with equilibration buffer and eluted with equilibration buffer supplemented with 300mM imidazole. After SDS-PAGE analysis, elution fractions containing SpyCas9 were pooled and dialysed overnight at 4°C against 20mM HEPES pH 7.5, 500mM KCl and 10% glycerol. The next day, the SpyCas9 sample was concentrated to ∼10mg/mL, aliquoted and flash-frozen in liquid nitrogen for long-term storage. HF1 SpyCas9 contains an N-terminal MBP-fusion tag. In order to remove it, Ni-NTA elution was followed by overnight TEV cleavage, and reverse Ni-NTA before proceeding with the overnight dialysis step.

For the long protocol, in order to remove any bound bacterial nucleic acids and endotoxins after elution from the Ni-NTA column, WT SpyCas9 protein was further purified using a combination of anion exchange (IEX) and heparin chromatography. The elution fractions from the Ni-NTA were diluted 8-fold with 20mM Tris-HCl pH 8.0, 250mM NaCl, 20mM imidazole, 1mM DTT and 10% glycerol (IEX equilibration buffer) and then loaded onto a 5mL HiTrap Q HP column (Cytiva) coupled in tandem to a 1mL HiTrap Heparin HP column (Cytiva). After washing with IEX equilibration buffer, the HiTrap Q HP column (which should bind the endotoxin molecules) was removed and SpyCas9 protein was eluted from the HiTrap Heparin HP column in a gradient going from 250mM NaCl to 1M NaCl over 20 column volumes (SpyCas9 protein usually elutes ∼650mM NaCl). The elution fractions containing SpyCas9 were pooled, concentrated to ∼5mL and injected into a HiLoad 16/600 Superdex 200pg size exclusion chromatography column (SEC) pre-equilibrated with 2mM HEPES pH 7.5, 500mM KCl and 10% glycerol. SpyCas9 elution fractions were pooled and concentrated to ∼10-15mg/mL. The final SpyCas9 samples were aliquoted, flash-frozen in liquid nitrogen and stored at -80°C until usage. The yield is usually around 7mg of pure SpyCas9 protein from 1L expression culture.

The identity of the SpyCas9 protein was verified by mass spectrometry and the oligomerization state and absence of aggregates was checked by SEC-MALS and Refeyn mass photometry. The stability in the storage buffer was assessed by nano-Differential Scanning Fluorimetry (nano-DSF; T_m_ ∼45°C).

### Transfection of *Dictyostelids*

The 20nt spacer region of crRNA was custom designed for each target using Cas designer (http://www.rgenome.net/cas-designer/), by selecting SpyCas9 as PAM target (5’-NGG-3’), *D. discoideum* as target genome, a spacer length of 20nt, and providing a target sequence of interest (e.g., *act5*, *mCherry*). We selected spacer sequences without any off-target hits for 0, 1 or 2 mismatches, favoring spacers with an out-of-frame score >66 and GC content of >20%. Targets with 4 or more sequential thymidines, which can be problematic when expressing a sgRNA from a plasmid as they may cause termination by PolIII, do not need to be excluded for our protocol. crRNA was ordered as Alt-R CRISPR-Cas9 crRNA from IDT.

For axenic strains, cells were grown in HL5+FAB to exponential phase (see above), harvested by centrifuging 100mL cell suspension (2 x 50mL, 400xg, 5 min), and washed three times in 1mL ice-cold H50 buffer (Text S2). For non-axenic strains, four SM5-agar plates were grown for 3-5 days (see above) and cells were collected from the feeding front using a 10µL loop, centrifuged (400xg, 2 min) and washed three times in 1mL ice-cold H50 buffer. For both axenic and non-axenic strains, cells were counted using the Countess III cell counter and aliquoted to 1.25-5·10^6^ cells per transfection (Figure S11A). Cells were kept on ice while preparing the RNP mix.

For each transfection, a 10µL RNP complex was prepared in a 1.5mL Eppendorf tube at RT by mixing the following reagents in order. Each reagent was added slowly, while swirling the tip into the mixture, gently pipetting up and down once or twice after adding the reagent: (1) 12µM SpyCas9 (ultrapure from EMBL PeP-core facility), (2) 83mM KCl, (3) 17mM HEPES pH7.5, (4) 22µM Alt-R CRISPR-Cas9 crRNA, (4) 22µM Alt-R CRISPR-Cas9 tracrRNA (IDT) in a total volume of 10µL. As an optional step, crRNA and tracrRNA can be mixed and annealed to form the gRNA complex by incubating and 95°C for 5 min, and letting it cool down slowly, before mixing it with SpyCas9. For knock-outs, single-stranded donor oligos (IDT) were suspended at a concentration of 100µM in nuclease-free water and added to the RNP complex. Volume was adjusted to 100µL with H50 buffer. Oligo concentration in the final 100µL transfection mix ranged from 0-4.8µM. See Table S2 for crRNA and donor oligo used to produce each knock-out strain. Donor oligos under 100nt (e.g., the donor oligo we used for knock-outs with 37bp insertion and 28bp homology arms) can be synthesized as regular oligos. The efficiency of these low-cost oligos is similar to high-quality Ultramer donor oligos (Figure S11B) and can therefore be used indistinctively.

For knock-ins, donor PCR product was PCR-amplified from pMG005 (for *mNeonGreen* reporter constructs) or pMG008 (for *mCherry* reporter constructs) (Text S1 and File S1) using custom primers (Table S4 for primer set, and Table S6 for primer sequence) with 30 to 40bp homology arms to the CRISPR target site, purified using Qiagen PCR purification MiniElute column and eluted in 12µL nuclease-free water. The concentration of the donor PCR product was adjusted to 1µg/µL (1.8µM for an 899bp PCR product consisting of mNeonGreen-P2A and 34/37bp homology arms). Amounts between 0 to 20µg (0 to 36.12pmol) were added to the RNP mix as indicated. When indicated, a non-specific oligo (Table S6) and/or Alt-R Cas9 Electroporation Enhancer (IDT) was added to the RNP mix as well (Figure S6 and S7). After all the components were added, H50 buffer was slowly added to the mix at RT to have a final volume of 100µL. When indicated, Alt-R HDR Enhancer V2 (IDT) was added to the growth media.

Once the transfection mix (containing the RNP complex, donor DNA (when indicated), enhancers (when indicated) and H50 electroporation buffer) was ready, aliquots of *Dictyostelid* cells were centrifuged (400xg, 2 min) and resuspended in 100µL of transfection mix. 80µl were transferred to a pre-chilled Gene Pulser/MicroPulser Electroporation Cuvettes (Bio-Rad, Text S2) and electroporated using a Gene Pulser Xcell Microbial System with the following settings: Exponential protocol; voltage, 750V; capacitance, 25µF; resistance, infinite; cuvette, 1mm; number of pulses, 1. Time constant was between 0.7 and 1ms. Electroporated cells were kept on ice for 5 min after electroporation. For flow cytometry analysis, 20µL of electroporated cells were transferred to a 24-well plate with either 1mL of HL5+FAB medium or 1mL *E. coli* B/r suspension (OD_600_ = 1-2 in KK2-MC) per well for axenic and non-axenic growth, respectively. For sorting, 80µL of electroporated cells were transferred to a 6-well plate containing 5mL HL5+FAB or *E. coli* B/r suspension (OD_600_ = 1-2 in KK2-MC). For both cases, cells were incubated for 1 to 3 days at 22°C. Fresh media was added every 24 hours.

### Flow cytometry analysis and cell sorting

For flow cytometry, cells were harvested 2- or 3-days post-transfection. Supernatant was first carefully removed from the wells and adherent cells were subsequently washed with 1mL KK2-MC buffer and then resuspended by adding 500µL KK2-MC and gently pipetting up and down. 200µL of resuspended cells were transferred to FACS tubes for analysis. DRAQ7 (ThermoFisher) was used as a viability dye at a 1/100 dilution. Samples were analysed using the BD FACSymphony™ A3 analyzer (5 laser configuration) using BD FACSDiva software Version 9.1.2 and the following settings: FSC: 100 V, SSC: 150 V, 488-530_30: 285 V, 561-610_20: 434 V, 640-670_30: 500 V. 10.000 events were recorded at a flow rate of up to 500 events/sec. Results were analyzed using FlowJo Version 10.9.0, by gating against bacterial cells (for non-axenic cultures), dead cells (DRAQ7 negative), and doublets. For the remaining single amoebal cells, we gated based on fluorescent signal.

For sorting, cells were typically harvested 1- or 2-days post-transfection. Supernatant was carefully removed from the wells, cells adhering to the surface were washed with 5mL KK2-MC buffer containing 2x streptomycin and penicillin, and harvested in 500µL KK2-MC with 2x streptomycin and penicillin (Text S2). Resuspended cells were collected in a 5mL round-bottom tube with a 35μm mesh cell strainer and sorted using BD FACSAria Fusion Flow Cytometer using BD FACSDiva software Version 8.0.1 with the following settings: FSC, 0V; SSC, 242V; 488-530 (30), 362V; 561-610 (20), 539V; 640-670 (30), 466V; nozzle, 100µm; sorting target, 1 cell; flow rate up to 400 cells/sec. Single live (DRAQ7 negative) cells were sorted based on fluorescence in individual wells from a 24-well plate, where each well contained 1mL SM5 charcoal agar (Text S2) and a 25µL-lawn of *E. coli* B/r suspension (OD_600_ 0.66 in of KK2-MC), which was carefully spread across the well, with a glass Pasteur pipette shaped into a spreader, and dried under a laminar hood for 15 min. Sorting results were analyzed using FlowJo Version 10.9.0. Sorted cells were incubated at 22°C for 3 to 5 days, until growth was observed and then imaged using a Zeiss AxioZoom V16 (see below).

### Microscopy

For imaging slugs in Figure 5, clones were propagated from sorted wells to SM5 agar plates, containing 0.5% charcoal and a lawn of *E. coli* B/r, until a feeding front and consequently slugs could be detected (charcoal was added to minimize background fluorescence from the agar medium). As control, the same procedure was followed for WT cells. Both WT and edited slugs were imaged using the Leica Stellaris 8 inverted confocal microscope with a 10x 0.3 NA air objective. White light laser wavelength was tuned to 488nm with a smart intensity of 50.47%. A NF 488/561/730 Polarisation filter was used to eliminate fluorescent signal from the cellulose stalk. A HyD detector was used with gain of 26.6 and a collection window of 494nm-586nm. Resolution was 1024×512 pixels, scan speed was 400 Hz, line averaging of 2 was used, and pixel dwell time was 1.4125µsec. Z-stacks were taken to capture the entire slug, with a z-step size of 4.017µm. All samples were imaged on the same day with the same acquisition settings to ensure accurate quantitative analysis. Confocal images of cells in Figure 6 were taken with a Nikon Ti2-E CSU-W1 spinning-disk confocal microscope with Hamamatsu Orca FusionBT camera with a resolution of 2048×2048 pixels. Images were acquired either with a CFI Apo LWD 40x 1.15 NA water immersion (MRD77410) or CFI P-Apo Ph3 100x 1.45 NA oil (MRD31905) objective. For 40x images (Figure 6B), cells were imaged in wide-field mode with a Lumencor Spectra III LED source and 475/28 bandpass filter (200ms exposure time). For 100x images (Figure 6D), cells were imaged in spinning-disk confocal mode with 488nm laser (37.7% intensity). Z-stacks were taken to capture entire cells, with a z-step size of 0.2µm with 25 steps. Raw image data as well as expression profiles are provided in File S2, S3 and S4. See ‘Data analysis’ below for a description of our quantification methods.

Imaging of multi-well plates after cell sorting and time-lapse imaging were acquired using a Zeiss AxioZoom V16 microscope with a Plan Z 1.0x objective and an Axiocam 506 mono camera with a 1x camera adapter. For time-lapse imaging, temperature was maintained at 22°C using an Ibidi stage incubator (Silver Line, TC3) and a room temperature of 18°C. In-well temperature was checked using the supplied test probe. Time-lapse imaging experiments for Figure 7 were carried out in 6-well plates with SM5 charcoal agar and *E. coli* B/r as food source in two rounds (see also File S5 and S6), either with *D. discoideum, D. firmibasis,* and *P. violaceum*, or with *D. lacteum, P. pallidum*, and *D. purpureum*. In each round of imaging, both fluorescent genome-edited strains and WT controls for each species were imaged. Each well was imaged at a zoom of 1x and a 4×4 tile scan. Imaging was performed with fluorescence (Excelitas X-Cite Xylis LED unit, 73% power, Zeiss 38 HE GFP filter cube) with exposure time of 1 sec, two white light spotlights (Zeiss CL 9000 LED unit, 40% power) arranged from above pointing down, and transmitted light from below (Zeiss CL 9000 LED unit, 60% power). Time-lapse images in Figure 7 and File S6 were processed in the following way: Flatfield correction was performed using Fiji and the Biovoxxel toolbox (https://github.com/biovoxxel/Biovoxxel-Toolbox) with the first time point as template. Brightness, contrast, and gamma levels were adjusted to best show both cell spreading and aggregation. Adjustments varied between species, but were identical for control and fluorescent strains from each species (see File S5 and S6). Tile stitching was done in ZEN 3.10 using default settings. All supplementary files (File S1-S6) are deposited to Zenodo (https://doi.org/10.5281/zenodo.15039721).

### Quick genomic DNA (gDNA) extraction and confirmation of clones

Once the fruiting bodies were visible (3-5 days after sorting), 3-5 sorus were picked with a small tip and submerged into a PCR tube containing 20µL of Quick Genomic DNA extraction buffer (Text S2). Cells were lysed at 56°C for 45 min, and then heat inactivated at 95°C for 10 min. Primers around the editing site were designed for each construct (see primers Table S6). 2µL of gDNA were used in a 20µL clone-confirmation PCR reaction with Taq polymerase. 2µL of the PCR product were run in a 1% agarose gel for visualization of the results. Positive PCR reactions were purified with Qiagen QIAquick PCR Purification Kit, and the PCR fragments were eluted with 25µl of EB. When indicated, PCR fragments were Sanger sequenced using Eurofins TubeSeq Supreme run.

### gDNA extraction and nanopore sequencing

To extract gDNA, monoclonal cells were grown axenically in HL5+FAB until reaching exponential phase (0.8-1.2·10^6^ cells/mL). For each clone, 3·10^7^ cells were harvested by centrifugation at 400xg for 5 min and gDNA was extracted as previously described (Pilcher et al. 2007). Library prep and Nanopore sequencing was performed by EMBL’s Genomics Core (GeneCore) Facility. In brief, DNA fragment sizes were analysed using the Femto Pulse Systems (Agilent), and samples were chosen which had a suitable proportion of longer fragments (>10 kb). Shorter fragments were depleted from the samples using PacBio SMRTbell cleanup beads (PacBio 102-158-300) at a 3.7x ratio and selected fragments resuspended in PacBio Elution Buffer (PacBio 101-633-500). Libraries were prepared for long read sequencing using the Ligation sequencing gDNA – Native Barcoding Kit 24 V14 (Oxford Nanopore Technologies). 10fmol of library was loaded on an R10 GridION flowcell for sequencing.

### Nanopore sequence analysis

For genome assembly, raw Nanopore reads were base-called live during sequencing using Dorado 7.4.14 (https://github.com/nanoporetech/dorado). Passed base-called nanopore reads were concatenated per sample. The *D. discoideum* AX2 *act5*::*mCherry* nanopore reference was assembled by combining reads from all our *D. discoideum* AX2 strains with *act5*::*mCherry* to maximize coverage and thereby assembly quality (see Table S3). *De novo* genome assembly was performed using Flye 2.9.2-b1786 (Kolmogorov et al. 2019) using the following parameter settings: `flye --nano-raw concatenated_reads.fastq -t 32 -o assembly`. Our assembly showed a 96.596% completeness compared to the RefSeq genome (GCA_000004695.1) of *D. discoideum* AX4 (Eichinger et al. 2005), as determined by QUAST v5.2.0. Differences between our *D. discoideum AX2 act5*::*mCherry* (MG38; Table S1) assembly and the RefSeq genome were determined using Minimap2 2.22-r1101 (Li 2018). Except for few minor genomic rearrangements (e.g., inversions) and the *act5*::*mCherry* integration no markable differences were detected (Figure S3). Indexes for aligned fasta files were derived using samtools 1.21. Visualization was done using D-Genies (Figure S3) (Cabanettes and Klopp 2018), selecting Minimap2 v2.28 as the aligner and ‘Many repeats’ as the level of repeatedness. To examine gene edits in *mCherry* and potential off-target integration of donor oligos, the *mCherry* sequence flanking the integration site as well as 37nt donor oligo (without homology arms) were separately queried against all raw nanopore reads using nucleotide-nucleotide BLAST 2.15.0+ (Camacho et al. 2009) using the following command line: `makeblastdb -in $nanopore_reads.fasta -dbtype nucl -out $reads_with_hit.db blastn -query [query sequence] -db $ reads_with_hit.db -outfmt 6 -out $reads_to_check.txt’`. All BLAST hits were subsequently examined manually by aligning the read against our *de novo* genome assembly using SnapGene 8.0.2 (www.snapgene.com; option ‘Aligned to Reference DNA Sequence’). In addition, all BLAST hits were also exported to a fasta file and automatically aligned against our reference genome using Minimap2 2.22-r1101. The resulting BAM files were furthermore filtered (`-q 30` and `-F 0×900`) and sorted using samtools 1.21, making it possible to visualize the alignment against our reference genome using Integrative Genomics Viewer (IGV) (Table S3). No incongruencies were detected, meaning that all *mCherry* genes had the expected edit and all donor oligos were integrated in the expected *mCherry* site (i.e., no off-target integration; Table S3). The Nanopore sequence analysis was performed on EMBL’s HPC Cluster (https://doi.org/10.5281/zenodo.12785829).

All sequencing-related files (as listed in Table S3) are deposited to Zenodo (https://doi.org/10.5281/zenodo.15039721) with folder (A) containing all passed base-called Nanopore reads, (B) containing our annotated *de novo D. discoideum* AX2 *act5*::*mCherry* genome assembly (our reference genome), (C) alignment of all reads against our reference genome (i.e., original BAM files), (D) alignment of all reads containing *mCherry* against our reference genome (see Table S3), (E) alignment of all reads containing donor oligo against our reference genome (see Table S3).

## Data analysis

Knock-out and knock-in percentages were quantified from flow cytometry data using FlowJo Version 10.9.0. Maximum knock-out or knock-in efficiencies were assessed by fitting a sigmoidal curve as a function of donor DNA (oligo or PCR product) concentration in R (v4.2.2), using a non-linear least square fitting (nls function): % = *max*/(1 + *e^scaling·(mid-[donor])^*). Estimated mean and standard error for the maximal editing efficiency (*max*) and inflection point (*mid*) are provided in Data S1 for each figure (Figure 1, 2, 3, S4, S7, and S11). Sigmoidal fits are only shown when estimated parameter values are significant (*p* < 0.05). Plots in other figures were generated in R (v4.2.2) as well. Microscopy images were processed using either Fiji (v1.53t) or Matlab R2024a. For Figure 5C-D, maximum projections were acquired for each slug z-stack and then a 20µm-wide spline was manually drawn from anterior to posterior to quantify expression profiles. All raw image data and expression profiles are provided in File S3. Profiles of individual slugs in Figure 5C-D are projected from anterior to posterior using a 20µm-rolling window. Slugs in Figure 5D were artificially straightened along their anterior-posterior axis, to facilitate the comparison of *ecmA* expression between slugs of different sizes, but raw image data are provided in File S3. For Figure 6C-D, cells were segmented based on their maximum projection. Then, the distance of pixels to the cell periphery was measured using `bwdist` (function in Matlab) to project expression profiles along the radial axis of cells.

For each cell, expression intensities are projected using a 2µm-rolling window from the cell periphery to center. Maximum projections, pixel intensities and distances are provided in File S4. The phylogenetic tree in Figure 7 and S10 was constructed using RaxML, based on 18S rDNA sequence alignment of (Sheikh et al. 2018), using the following parameter settings: `raxmlHPC -f d -m GTRGAMMA -n TREE -s 1-s2.0-S1434461017300925-mmc13.fasta -# 100 -b 1 -p 1`. All supplementary files (File S1-S6) and data (Data S1) are deposited to Zenodo (https://doi.org/10.5281/zenodo.15039721).

## Acknowledgement

We thank everybody from the van Gestel group for feedback and suggestions. We particularly thank Öykü Bozalioğlu for establishing efficient electroporation conditions, Haohan Zhang for testing different isolation protocols and Vanessa Stürmer for her support in taking microscopy images. We thank EMBL’s Advanced Light Microscopy Facility (ALMF), Flow Cytometry Core Facility, and GeneCore Facility for their support. We thank Peggy Paschke for her strains and plasmids as well as her generous advice on genome editing methods. We thank Elizabeth Ostrowski and Pauline Schaap for their advice on culturing diverse *Dictyostelid* species. We thank Allyson Sgro, Emily Hager, John Durel and Elizabeth Ostrowski for their feedback on the manuscript. We thank EMBL for financial support. AP also received support from iXcore – iXlife – iXblue foundation. JvG received support from the European Union (ERC, CO-PP, 101116560). Views and opinions expressed are however those of the author(s) only and do not necessarily reflect those of the European Union or the European Research Council Executive Agency. Neither the European Union nor the granting authority can be held responsible for them.

## References

1. Akella, S., X. Ma, R. Bacova, Z. P. Harmer, M. Kolackova, X. Wen, D. A. Wright, et al. 2021. Co-targeting strategy for precise, scarless gene editing with CRISPR/Cas9 and donor ssODNs in *Chlamydomonas*. Plant Physiology 187:2637–2655.

2. Arribere, J. A., R. T. Bell, B. X. H. Fu, K. L. Artiles, P. S. Hartman, and A. Z. Fire. 2014. Efficient marker-free recovery of custom genetic modifications with CRISPR/Cas9 in *Caenorhabditis elegans*. Genetics 198:837–846.

3. Baldauf, S. L., M. Romeralo, O. Fiz-Palacios, and N. Heidari. 2018. A deep hidden diversity of *Dictyostelia*. Protist 169:64–78.

4. Bloomfield, G., Y. Tanaka, J. Skelton, A. Ivens, and R. R. Kay. 2008. Widespread duplications in the genomes of laboratory stocks of *Dictyostelium discoideum*. Genome Biology 9:R75.

5. Bloomfield, G., D. Traynor, S. P. Sander, D. M. Veltman, J. A. Pachebat, and R. R. Kay. 2015. Neurofibromin controls macropinocytosis and phagocytosis in *Dictyostelium*. eLife 4:e04940.

6. Bock, C., P. Datlinger, F. Chardon, M. A. Coelho, M. B. Dong, K. A. Lawson, T. Lu, et al. 2022. High-content CRISPR screening. Nature Reviews Methods Primers 2:1–23.

7. Bonner, J. T. 2009. The Social Amoebae: The Biology of Cellular Slime Molds. Princeton University Press, Princeton, NJ.

8. Bonner, J. T., and D. S. Lamont. 2005. Behavior of cellular slime molds in the soil. Mycologia 97:178–184.

9. Cabanettes, F., and C. Klopp. 2018. D-GENIES: dot plot large genomes in an interactive, efficient and simple way. PeerJ 6:e4958.

10. Camacho, C., G. Coulouris, V. Avagyan, N. Ma, J. Papadopoulos, K. Bealer, and T. L. Madden. 2009. BLAST+: architecture and applications. BMC Bioinformatics 10:421.

11. Cardenal-Muñoz, E., C. Barisch, L. H. Lefrançois, A. T. López-Jiménez, and T. Soldati. 2018. When Dicty met Myco, a (not so) romantic story about one amoeba and its intracellular pathogen. Frontiers in Cellular and Infection Microbiology 7:529.

12. Chen, G., O. Zhuchenko, and A. Kuspa. 2007. Immune-like phagocyte activity in the social amoeba. Science 317:678–681.

13. Combredet, C., and T. Brunet. 2024. A fast and robust gene knockout method for *Salpingoeca rosetta* clarifies the genetics of choanoflagellate multicellular development. bioRxiv.

14. Cosson, P., and T. Soldati. 2008. Eat, kill or die: when amoeba meets bacteria. Current Opinion in Microbiology 11:271–276.

15. De Lozanne, A., and J. A. Spudich. 1987. Disruption of the *Dictyostelium* myosin heavy chain gene by homologous recombination. Science 236:1086–1091.

16. Doudna, J. A., and E. Charpentier. 2014. Genome editing. The new frontier of genome engineering with CRISPR-Cas9. Science 346:1258096.

17. Eichinger, L., J. A. Pachebat, G. Glöckner, M. A. Rajandream, R. Sucgang, M. Berriman, J. Song, et al. 2005. The genome of the social amoeba *Dictyostelium discoideum*. Nature 435:43–57.

18. Faix, J., L. Kreppel, G. Shaulsky, M. Schleicher, and A. R. Kimmel. 2004. A rapid and efficient method to generate multiple gene disruptions in *Dictyostelium discoideum* using a single selectable marker and the Cre-loxP system. Nucleic Acids Research 32:e143.

19. Faktorová, D., R. E. R. Nisbet, J. A. Fernández Robledo, E. Casacuberta, L. Sudek, A. E. Allen, M. Ares, et al. 2020. Genetic tool development in marine protists: emerging model organisms for experimental cell biology. Nature Methods 17:481–494.

20. Fey, P., K. Compton, and E. C. Cox. 1995. Green fluorescent protein production in the cellular slime molds *Polysphondylium pallidum* and *Dictyostelium discoideum*. Gene 165:127–130.

21. Fey, P., R. J. Dodson, S. Basu, and R. L. Chisholm. 2013. One stop shop for everything *Dictyostelium*: dictyBase and the Dicty Stock Center in 2012. *Dictyostelium discoideum* Protocols, Methods in Molecular Biology.

22. Fey, P., A. S. Kowal, P. Gaudet, K. E. Pilcher, and R. L. Chisholm. 2007. Protocols for growth and development of *Dictyostelium discoideum*. Nature Protocols 2:1307–1316.

23. Francis, D. W. 1964. Some studies on phototaxis of *Dictyostelium*. Journal of Cellular and Comparative Physiology 64:131–138.

24. Fuqua, T., J. Jordan, M. E. van Breugel, A. Halavatyi, C. Tischer, P. Polidoro, N. Abe, et al. 2020. Dense and pleiotropic regulatory information in a developmental enhancer. Nature 587:235–239.

25. Gregor, T., K. Fujimoto, N. Masaki, and S. Sawai. 2010. The onset of collective behavior in social amoebae. Science 328:1021–1025.

26. Gruenheit, N., A. Baldwin, B. Stewart, S. Jaques, T. Keller, K. Parkinson, W. Salvidge, et al. 2021. Mutant resources for functional genomics in *Dictyostelium discoideum* using REMI-seq technology. BMC Biology 19:172.

27. Hall, G., S. Kelly, P. Schaap, and C. Schilde. 2022. Phylogeny-wide analysis of G-protein coupled receptors in social amoebas and implications for the evolution of multicellularity. Open Research Europe 2:134.

28. Huang, Y. Y., M. N. Price, A. Hung, O. Gal-Oz, S. Tripathi, C. W. Smith, D. Ho, et al. 2024. Barcoded overexpression screens in gut Bacteroidales identify genes with roles in carbon utilization and stress resistance. Nature Communications 15:6618.

29. Iriki, H., T. Kawata, and T. Muramoto. 2019. Generation of deletions and precise point mutations in *Dictyostelium discoideum* using the CRISPR nickase. PloS One 14:e0224128.

30. Ishikawa-Ankerhold, H. C., and A. Müller-Taubenberger. 2019. Actin assembly states in *Dictyostelium discoideum* at different stages of development and during cellular stress. The International Journal of Developmental Biology 63:417–427.

31. Jauslin, T., O. Lamrabet, X. Crespo-Yañez, A. Marchetti, I. Ayadi, E. Ifrid, C. Guilhen, et al. 2021. How phagocytic cells kill different bacteria: a quantitative analysis using *Dictyostelium discoideum*. mBio 12:e03169–20.

32. Jermyn, K. A., and J. G. Williams. 1991. An analysis of culmination in *Dictyostelium* using prestalk and stalk-specific cell autonomous markers. Development 111:779–787.

33. Jinek, M., K. Chylinski, I. Fonfara, M. Hauer, J. A. Doudna, and E. Charpentier. 2012. A programmable dual-RNA-guided DNA endonuclease in adaptive bacterial immunity. Science 337:816–821.

34. Jones, E. M., N. B. Lubock, A. Venkatakrishnan, J. Wang, A. M. Tseng, J. M. Paggi, N. R. Latorraca, et al. 2020. Structural and functional characterization of G protein–coupled receptors with deep mutational scanning. eLife 9:e54895.

35. Jones, G. M., J. Stalker, S. Humphray, A. West, T. Cox, J. Rogers, I. Dunham, et al. 2008. A systematic library for comprehensive overexpression screens in *Saccharomyces cerevisiae*. Nature Methods 5:239–241.

36. Joseph, J. M., P. Fey, N. Ramalingam, X. I. Liu, M. Rohlfs, A. A. Noegel, A. Müller-Taubenberger, et al. 2008. The actinome of *Dictyostelium discoideum* in comparison to actins and actin-related proteins from other organisms. PloS One 3:e2654.

37. Kay, R. R., J. E. Lutton, J. S. King, and T. Bretschneider. 2024. Making cups and rings: the “stalled-wave” model for macropinocytosis. Biochemical Society Transactions 52:1785–1794.

38. Kessin, R. H., G. G. Gundersen, V. Zaydfudim, and M. Grimson. 1996. How cellular slime molds evade nematodes. Proceedings of the National Academy of Sciences 93:4857–4861.

39. Kim, S., D. Kim, S. W. Cho, J. Kim, and J.-S. Kim. 2014. Highly efficient RNA-guided genome editing in human cells via delivery of purified Cas9 ribonucleoproteins. Genome Research 24:1012–1019.

40. Kin, K., Z. Chen, G. Forbes, H. Lawal, C. Schilde, R. Singh, C. Cole, et al. 2023. The protein kinases of Dictyostelia and their incorporation into a signalome. Cellular Signalling 108:110714.

41. Kolmogorov, M., J. Yuan, Y. Lin, and P. A. Pevzner. 2019. Assembly of long, error-prone reads using repeat graphs. Nature Biotechnology 37:540–546.

42. Kundert, P., A. Sarrion-Perdigones, Y. Gonzalez, M. Katoh-Kurasawa, S. Hirose, P. Lehmann, K. J. T. Venken, et al. 2020. A GoldenBraid cloning system for synthetic biology in social amoebae. Nucleic Acids Research 48:4139–4146.

43. Kuserk, F. T. 1980. The relationship between cellular slime molds and bacteria in forest soil. Ecology 61:1474–1485.

44. Kuspa, A., and W. F. Loomis. 1992. Tagging developmental genes in *Dictyostelium* by restriction enzyme-mediated integration of plasmid DNA. Proceedings of the National Academy of Sciences of the United States of America 89:8803–8807.

45. Kuwayama, H., S. Obara, T. Morio, M. Katoh, H. Urushihara, and Y. Tanaka. 2002. PCR-mediated generation of a gene disruption construct without the use of DNA ligase and plasmid vectors. Nucleic Acids Research 30:E2.

46. Kuzdzal-Fick, J. J., A. Moreno, C. M. E. Broersma, T. F. Cooper, and E. A. Ostrowski. 2023. From individual behaviors to collective outcomes: fruiting body formation in *Dictyostelium* as a group-level phenotype. Evolution 77:731–745.

47. Landolt, J. C., S. L. Stephenson, and J. C. Cavender. 2006. Distribution and ecology of Dictyostelid cellular slime molds in Great Smoky Mountains National Park. Mycologia 98:541–549.

48. Li, C. L. F., B. Santhanam, A. N. Webb, B. Zupan, and G. Shaulsky. 2016. Gene discovery by chemical mutagenesis and whole-genome sequencing in *Dictyostelium*. Genome Research 26:1268–1276.

49. Li, H. 2018. Minimap2: pairwise alignment for nucleotide sequences. Bioinformatics 34:3094– 3100.

50. Linkner, J., B. Nordholz, A. Junemann, M. Winterhoff, and J. Faix. 2012. Highly effective removal of floxed Blasticidin S resistance cassettes from *Dictyostelium discoideum* mutants by extrachromosomal expression of Cre. European Journal of Cell Biology 91:156–160.

51. Loomis, W. F. 1987. Genetic tools for *Dictyostelium discoideum*. Methods in Cell Biology 28:31–65.

52. Loomis, W. F. 2015. Genetic control of morphogenesis in *Dictyostelium*. Developmental Biology 402:146–161.

53. McRobbie, S. J., R. Tilly, K. Blight, A. Ceccarelli, and J. G. Williams. 1988. Identification and localization of proteins encoded by two DIF-inducible genes of *Dictyostelium*. Developmental Biology 125:59–63.

54. Medina, J. M., P. M. Shreenidhi, T. J. Larsen, D. C. Queller, and J. E. Strassmann. 2019. Cooperation and conflict in the social amoeba *Dictyostelium discoideum*. The International Journal of Developmental Biology 63:371–382.

55. Miranda, E. R., O. Zhuchenko, M. Toplak, B. Santhanam, B. Zupan, A. Kuspa, and G. Shaulsky. 2013. ABC transporters in *Dictyostelium discoideum* development. PloS One 8:e70040.

56. Muramoto, T., H. Iriki, J. Watanabe, and T. Kawata. 2019. Recent advances in CRISPR/Cas9-mediated genome editing in *Dictyostelium*. Cells 8:E46.

57. Narita, T. B., Y. Kawabe, K. Kin, R. A. Gibbs, A. Kuspa, D. M. Muzny, S. Richards, et al. 2020. Loss of the polyketide synthase StlB results in stalk cell overproduction in *Polysphondylium violaceum*. Genome Biology and Evolution 12:674–683.

58. Nomura, T., M. Yoshikawa, K. Suzuki, and K. Mochida. 2020. Highly efficient CRISPR-associated protein 9 ribonucleoprotein-based genome editing in *Euglena gracilis*. STAR Protocols 1:100023.

59. Ogasawara, T., J. Watanabe, R. Adachi, Y. Ono, Y. Kamimura, and T. Muramoto. 2022. CRISPR/Cas9-based genome-wide screening of *Dictyostelium*. Scientific Reports 12:11215.

60. Ostrowski, E. A. 2019. Enforcing cooperation in the social amoebae. Current biology: CB 29:R474–R484.

61. Paix, A., A. Folkmann, D. H. Goldman, H. Kulaga, M. J. Grzelak, D. Rasoloson, S. Paidemarry, et al. 2017a. Precision genome editing using synthesis-dependent repair of Cas9-induced DNA breaks. Proceedings of the National Academy of Sciences of the United States of America 114:E10745–E10754.

62. Paix, A., A. Folkmann, and G. Seydoux. 2017b. Precision genome editing using CRISPR-Cas9 and linear repair templates in *C. elegans*. Methods 121–122:86–93.

63. Papkou, A., T. Guzella, W. Yang, S. Koepper, B. Pees, R. Schalkowski, M. C. Barg, et al. 2019. The genomic basis of Red Queen dynamics during rapid reciprocal host–pathogen coevolution. Proceedings of the National Academy of Sciences of the United States of America 116:923–928.

64. Paschke, P., D. A. Knecht, A. Silale, D. Traynor, T. D. Williams, P. A. Thomason, R. H. Insall, et al. 2018. Rapid and efficient genetic engineering of both wild type and axenic strains of *Dictyostelium discoideum*. PloS One 13:e0196809.

65. Paschke, P., D. A. Knecht, T. D. Williams, P. A. Thomason, R. H. Insall, J. R. Chubb, R. R. Kay, et al. 2019. Genetic engineering of *Dictyostelium discoideum* cells based on selection and growth on bacteria. JoVE 143:58981.

66. Pilcher, K. E., P. Fey, P. Gaudet, A. S. Kowal, and R. L. Chisholm. 2007. A reliable general purpose method for extracting genomic DNA from *Dictyostelium* cells. Nature Protocols 2:1325–1328.

67. Raper, K. B. 1935. *Dictyostelium discoideum*, a new species of slime mold from decaying forest leaves 50:135–147.

68. Raper, K. B. 1984. The Dictyostelids. Princeton University Press, Princeton, N.J.

69. Robinson, D. N., and J. A. Spudich. 2000. Dynacortin, a genetic link between equatorial contractility and global shape control discovered by library complementation of a *Dictyostelium discoideum* cytokinesis mutant. The Journal of Cell Biology 150:823–838.

70. Schaap, P., T. Winckler, M. Nelson, E. Alvarez-Curto, B. Elgie, H. Hagiwara, J. Cavender, et al. 2006. Molecular phylogeny and evolution of morphology in the social amoebas. Science 314:661–663.

71. Schilde, C., A. Skiba, and P. Schaap. 2014. Evolutionary reconstruction of pattern formation in 98 *Dictyostelium* species reveals that cell-type specialization by lateral inhibition is a derived trait. EvoDevo 5:34.

72. Sekine, R., T. Kawata, and T. Muramoto. 2018. CRISPR/Cas9 mediated targeting of multiple genes in *Dictyostelium*. Scientific Reports 8:8471.

73. Sheikh, S., M. Thulin, J. C. Cavender, R. Escalante, S. I. Kawakami, C. Lado, J. C. Landolt, et al. 2018. A new classification of the Dictyostelids. Protist 169:1–28.

74. Steele, M. I., J. M. Peiser, P. M. Shreenidhi, J. E. Strassmann, and D. C. Queller. 2023. Predation-resistant *Pseudomonas* bacteria engage in symbiont-like behavior with the social amoeba *Dictyostelium discoideum*. The ISME journal 17:2352–2361.

75. Steinert, M., and K. Heuner. 2005. *Dictyostelium* as host model for pathogenesis. Cellular Microbiology 7:307–314.

76. Stevense, M., J. R. Chubb, and T. Muramoto. 2011. Nuclear organization and transcriptional dynamics in *Dictyostelium*. Development, Growth & Differentiation 53:576–586.

77. Stewart, B., N. Gruenheit, A. Baldwin, R. Chisholm, D. Rozen, A. Harwood, J. B. Wolf, et al. 2022. The genetic architecture underlying prey-dependent performance in a microbial predator. Nature Communications 13:319.

78. Sussman, R., and M. Sussman. 1967. Cultivation of *Dictyostelium discoideum* in axenic medium. Biochemical and Biophysical Research Communications 29:53–55.

79. Sutoh, K. 1993. A transformation vector for *Dictyostelium discoideum* with a new selectable marker *bsr*. Plasmid 30:150–154.

80. Tomchik, K. J., and P. N. Devreotes. 1981. Adenosine 3′,5′-monophosphate waves in *Dictyostelium discoideum*: a demonstration by isotope dilution—fluorography. Science 212:443–446.

81. Tsuchiya, H. M., J. F. Drake, J. L. Jost, and A. G. Fredrickson. 1972. Predator-prey interactions of *Dictyostelium discoideum* and *Escherichia coli* in continuous culture. Journal of Bacteriology 110:1147–1153.

82. Urtecho, G., K. D. Insigne, A. D. Tripp, M. S. Brinck, N. B. Lubock, C. Acree, H. Kim, et al. 2023. Genome-wide functional characterization of *Escherichia coli* promoters and sequence elements encoding their regulation. eLife 12.

83. Van Haastert, P. J. M., and P. N. Devreotes. 2004. Chemotaxis: signalling the way forward. Nature Reviews Molecular Cell Biology 5:626–634.

84. Vines, J. H., and J. S. King. 2019. The endocytic pathways of *Dictyostelium discoideum*. The International Journal of Developmental Biology 63:461–471.

85. Witke, W., W. Nellen, and A. Noegel. 1987. Homologous recombination in the *Dictyostelium* alpha-actinin gene leads to an altered mRNA and lack of the protein. The EMBO journal 6:4143–4148.

86. Yamashita, K., H. Iriki, Y. Kamimura, and T. Muramoto. 2021. CRISPR toolbox for genome editing in *Dictyostelium*. Frontiers in Cell and Developmental Biology 9:721630.

87. Zhu, X., C. Ricci-Tam, E. R. Hager, and A. E. Sgro. 2023. Self-cleaving peptides for expression of multiple genes in *Dictyostelium discoideum*. PLoS One 18:e0281211.

88. Zucko, J., N. Skunca, T. Curk, B. Zupan, P. F. Long, J. Cullum, R. H. Kessin, et al. 2007. Polyketide synthase genes and the natural products potential of *Dictyostelium discoideum*. Bioinformatics 23:2543–2549.

