## Supplementary information for "Selection-free CRISPR-Cas9 editing protocol for distant *Dictyostelid* species"

#### Table of contents

| Supplementary Figures | Page |
|---|---|
| **Figure S1.** Quantification of knock-out mutants through flow cytometry. | 3 |
| **Figure S2.** Aligned Sanger sequencing results of knock-out mutants. | 4 |
| **Figure S3.** Comparison between *de novo* assembly and RefSeq genome. | 5 |
| **Figure S4.** Knock-out efficiency for different target sites and SpyCas9 proteins. | 6 |
| **Figure S5**. Flow cytometry data of knock-in experiments. | 7 |
| **Figure S6.** Quantification of carrier effect. | 8 |
| **Figure S7.** CRISPR-Cas9 mediated knock-ins with EE and HDRE. | 9 |
| **Figure S8.** Quantification of sorting accuracy. | 10 |
| **Figure S9.** CRISPR-Cas9 editing multiple loci simultaneously. | 11 |
| **Figure S10.** Phylogenetic tree with representative *Dictyostelid* species. | 12 |
| **Figure S11.** Knock-out efficiency for different cell densities and donor oligo qualities. | 13 |
| **Captions for Supplementary Files** | |
| **File S1.** Plasmid maps. | 14 |
| **File S2.** Time-lapse of strains with *ecmA* and *act5* expression reporters. | 14 |
| **File S3.** Raw data slug images. | 14 |
| **File S4.** Raw data cell images. | 14 |
| **File S5.** Videos of *Dictyostelid* species (low zoom). | 14 |
| **File S6.** Videos of *Dictyostelid* species (high zoom). | 14 |
| **Supplementary Tables** | |
| **Table S1.** Strain list. | 15 |
| **Table S2.** List of generated knock-out strains. | 15 |
| **Table S3.** Analysis of knock-out strains using Nanopore sequencing | 16 |
| **Table S4.** List of generated knock-in strains. | 17 |
| **Table S5.** List of crRNAs. | 18 |
| **Table S6.** List of oligos. | 19 |
| **Supplementary Text files** | |
| **Text S1.** Vector sequences. | 21 |
| **Text S2.** Detailed protocol. | 23 |
| **References** | 45 |

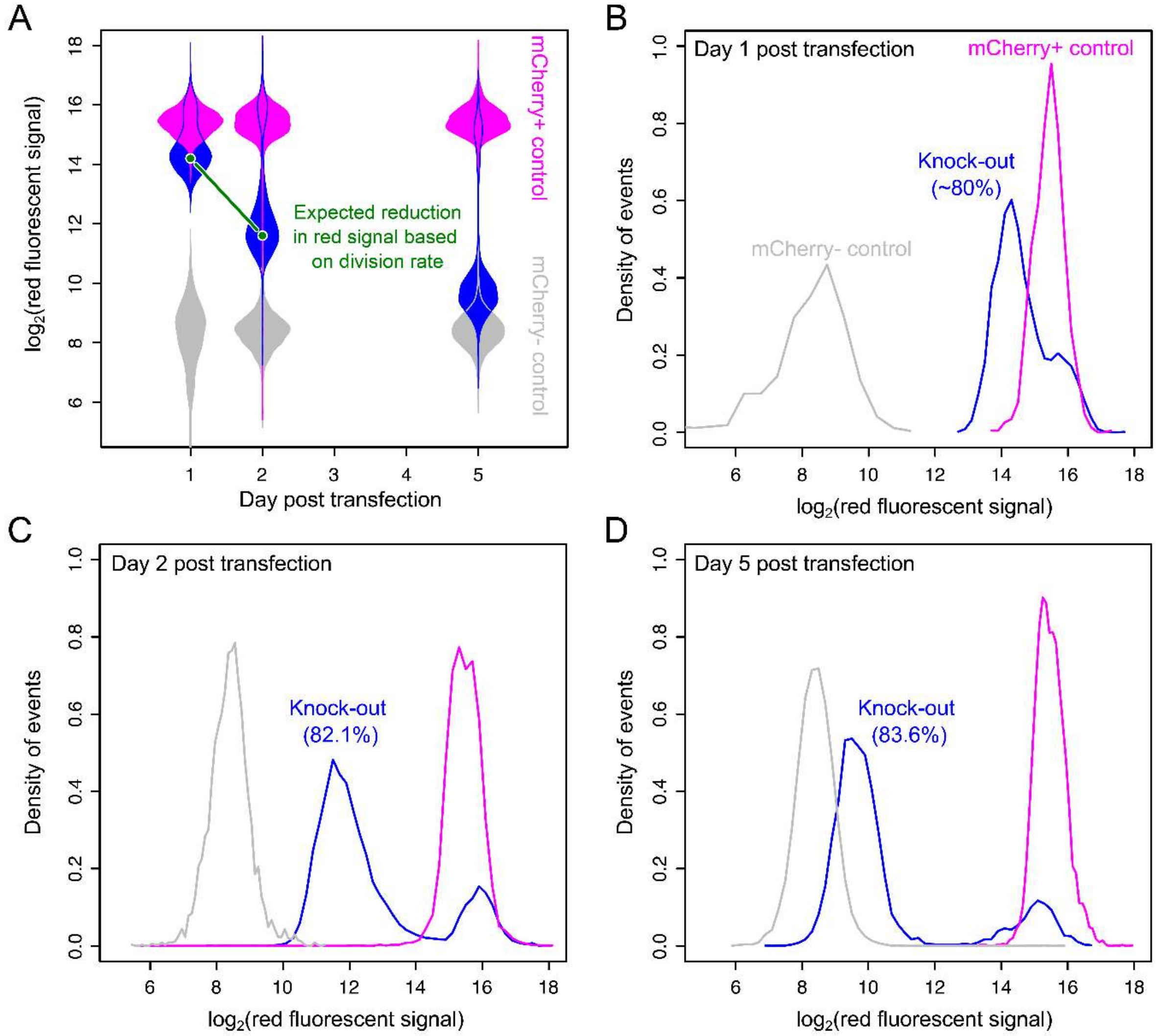

**Figure S1. Quantification of knock-out mutants through flow cytometry.** Knock-outs are produced by targeting amino acid 9 of *mCherry* CDS of the *D. discoideum* AX2 *act5*::*mCherry* strain with 2.4μM donor oligo (37bp insertion and 28bp homology arms). The same target and donor oligo are used for all knock-out experiments hereafter, unless noted otherwise. (A) Distribution of red fluorescent signal in CRISPR-Cas9 edited cells (blue), mCherry+ (magenta, *D. discoideum* AX2 *act5*::*mCherry*) and mCherry- (grey, *D. discoideum* AX2) control cells. Red fluorescent signal distribution is bimodal for CRISPR-Cas9 edited cells: some cells obtained knock-out mutation and therefore gradually lose fluorescent signal, whereas other cells were not edited and retain their fluorescent signal. Green line shows expected reduction in red fluorescent signal based on the expected dilution of mCherry protein per cell through cell division. The expected reduction perfectly follows the observed reduction. Distribution of red fluorescent signal for CRISPR-Cas9 edited cells, mCherry+ and mCherry- control cells (B) 1 day, (C) 2 days and (D) 5 days post-infection. Knock-out cells can be identified as soon as one day post-transfection, making it possible to isolate them after just one day.

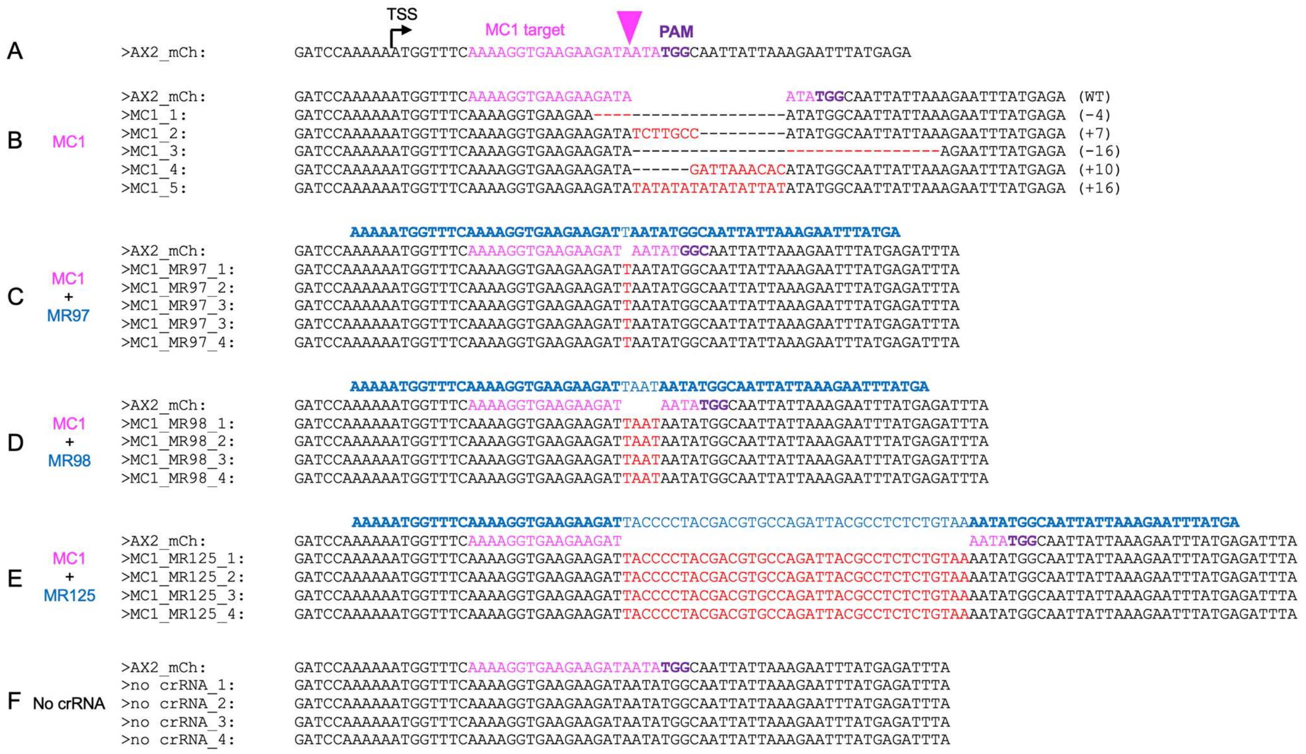

**Figure S2. Aligned Sanger sequencing results of knock-out mutants.** (A) Sequence of the *act5::mCherry* locus showing the target sequence (magenta), predicted cleavage site (magenta triangle) and downstream PAM (bold dark magenta) sequence. Sanger sequencing results for different clones of *D. discoideum* AX2 *act5::mCherry* transfected with SpyCas9, tracrRNA and (B) MC1 crRNA, (C) MC1 crRNA and donor oligo (blue) for 1bp insertion, (D) MC1 crRNA and donor oligo for 4bp insertion, (E) MC1 crRNA and donor oligo for 37bp insertion, or as control (F) no crRNA and no donor oligo. Indels are shown in red.

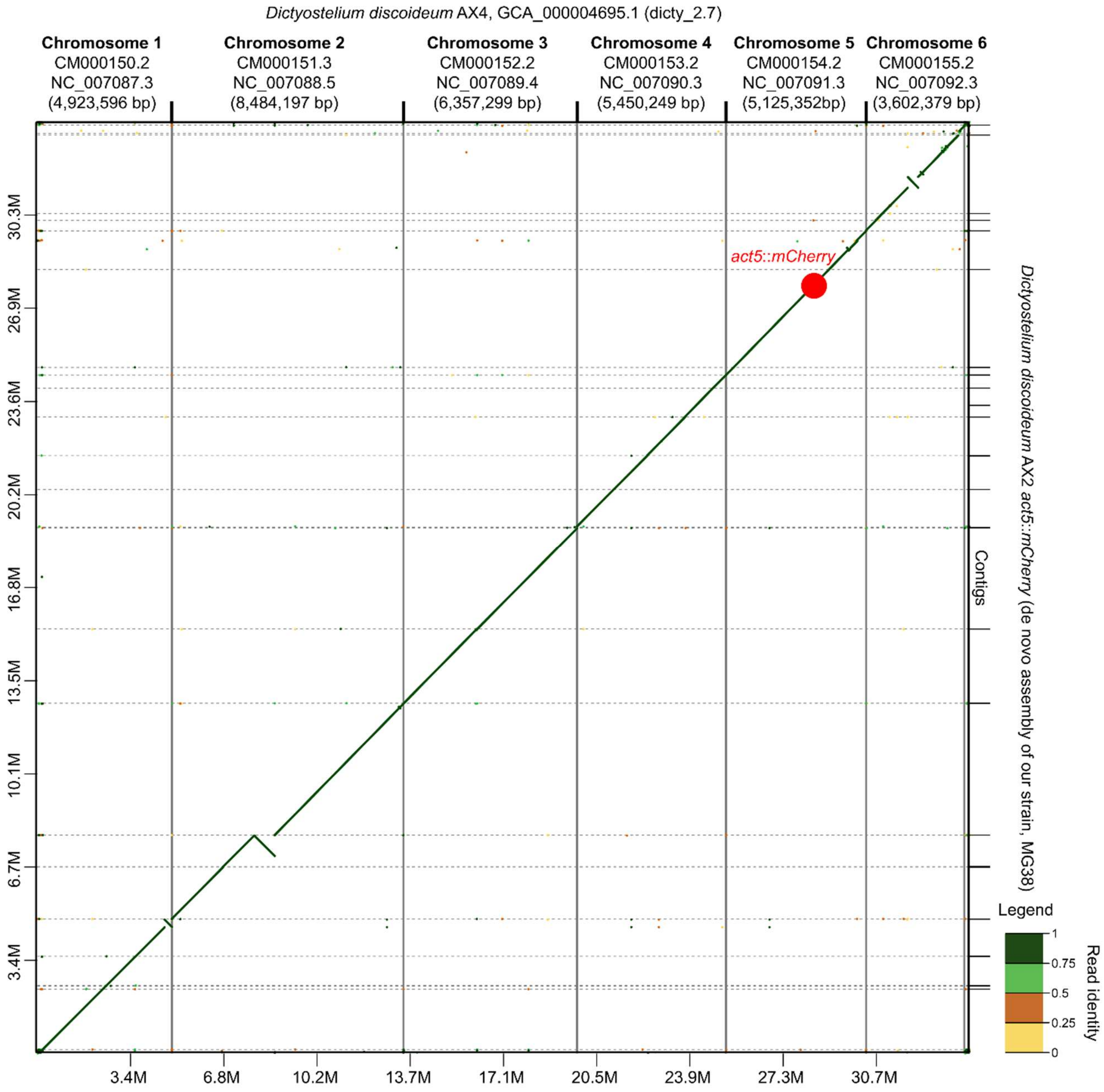

**Figure S3. Comparison between *de novo* assembly of our *D. discoideum* AX2 *act5*::*mCherry* strain and the *D. discoideum* AX4 RefSeq strain.** Mapping between chromosome-level assembly of *D. discoideum* AX4 (GCA_000004695.1) (x-axis) and our *de novo D. discoideum* AX2 *act5::mCherry* assembly (y-axis; MG38). For our *de novo* assembly, nanopore sequence data was pooled for all strains with an *act5*::*mCherry* integration site (Table S3). Small dots show mapped reads. Color legend shows quality of mapping: dark green is high-quality mapping and yellow is low-quality mapping. Single large red dot shows *act5::mCherry* integration site.

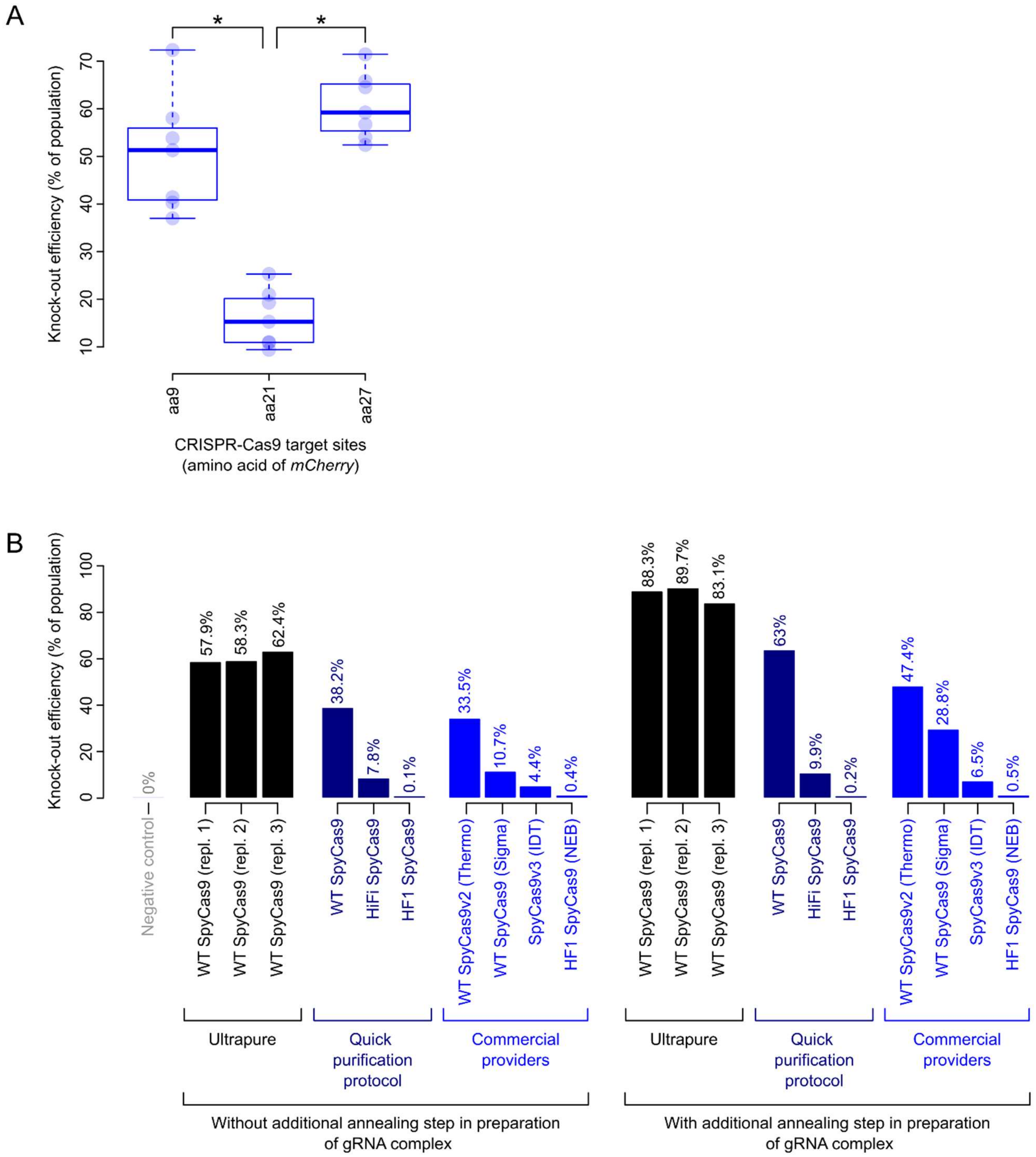

**Figure S4. Knock-out efficiency for different CRISPR-Cas9 target sites in *mCherry* and SpyCas9 proteins.** (A) Two of the three target sites show significantly higher knock-out efficiencies (pairwise Wilcoxon signed-rank test, $adjusted\ p < 0.01$). See Data S1 for data. (B) Knock-out efficiencies with different purification methods of SpyCas9 (a 3-day protocol to produce an ultrapure extract or a quick one-day protocol) and different versions of SpyCas9 (WT, HiFi and HF1). For ultrapure WT SpyCas9 three biological replicates are shown, using separate cell batches. Knock-out efficiencies of equivalent commercially available protein extracts, including TrueCut Cas9 Protein v2 (Thermo Fisher A36498), Cas9 Protein (Sigma-Aldrich CAS9PROT-50UG), Alt-S.p. Cas9 Nuclease V3 (Integrated DNA

Technologies (IDT) 1081058) and EnGen Spy Cas9 HF1 (New England Biolabs (NEB) M0667M) are also shown.

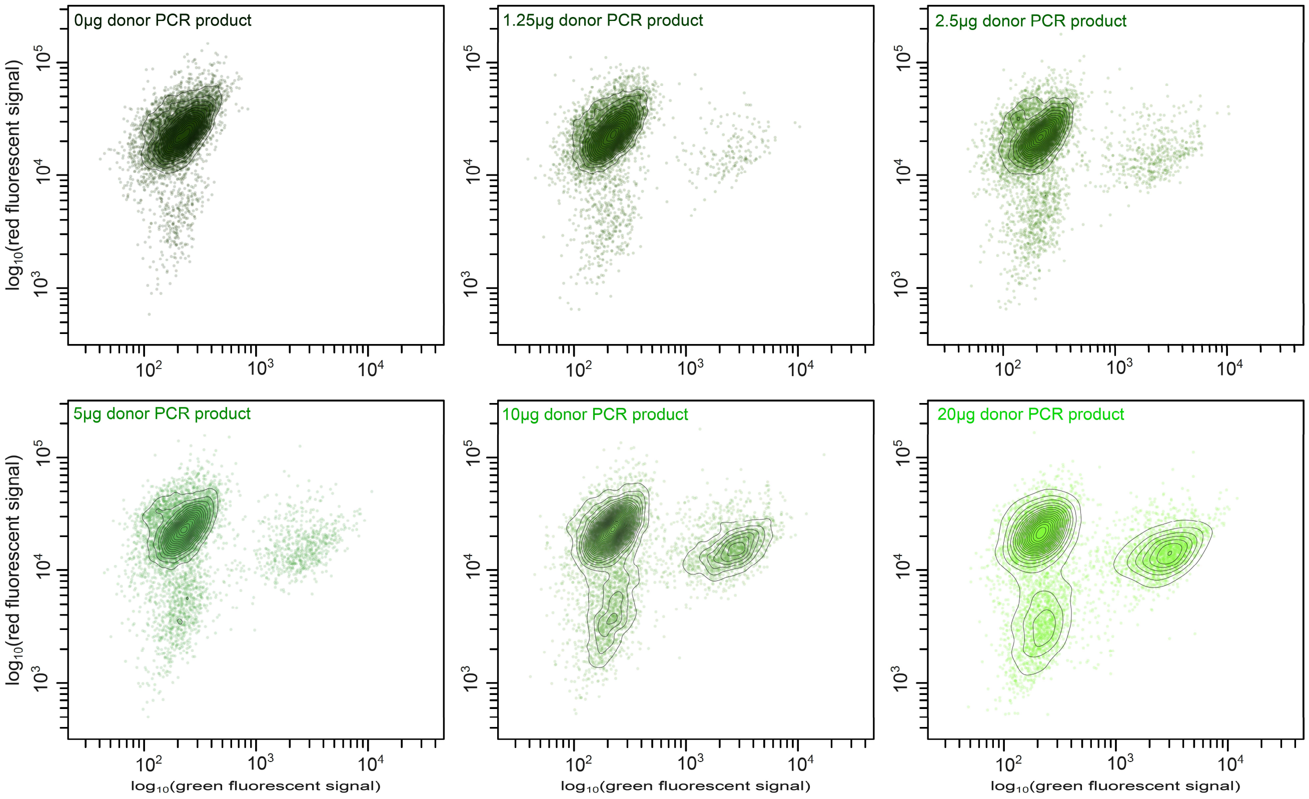

**Figure S5. Flow cytometry data of knock-in experiments with different concentrations of donor PCR products**. Knock-ins are produced by targeting amino acid 9 of *mCherry* CDS of *D. discoideum* AX2 *act5*::*mCherry* and a donor PCR product encoding for mNeonGreen-P2A in frame with mCherry. The same target and donor PCR product is used for all knock-ins experiments hereafter, unless noted otherwise. Flow cytometry results identical to those in Figure 2B, but separated by the quantity of donor PCR product added to the transfection: 0, 1.25, 2.5, 5, 10 or 20µg (0, 2.26, 4.52, 9.03, 18.06 or 36.12pmol respectively). Color codes correspond to Figure 2B. Contours indicate density of data points.

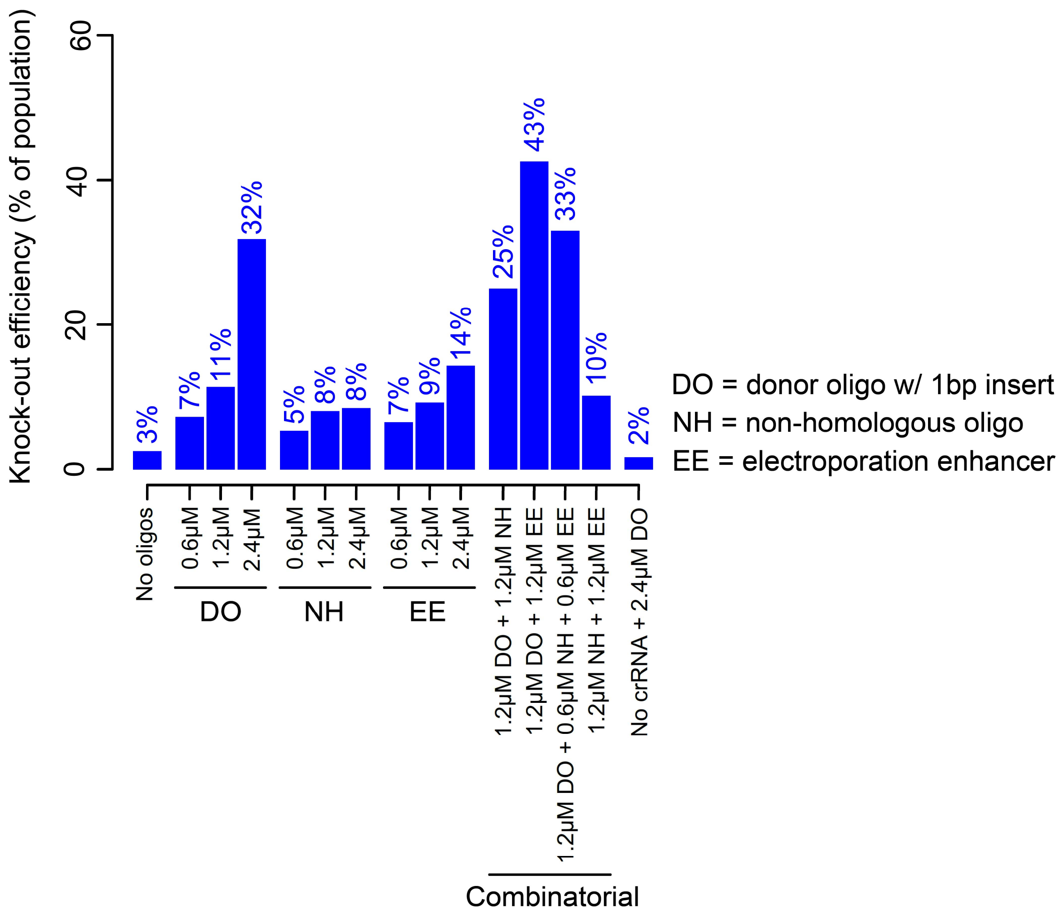

**Figure S6. Quantification of carrier effect using donor oligos (DO), non-homologous oligos (NH) and electroporation enhancer (EE).** Combinations of donor oligo (1bp insertion and 28/31bp left/right homology arms), NH 60nt oligo (Table S6), and EE (IDT) are added to discern the impact of donor oligos on homology-directed repair (HDR) and the carrier effect. Increasing amounts of donor oligo improves knock-out efficiencies. Addition of NH oligos and EE improves knock-out efficiency (8% and 14% maximum knock-out efficiency, respectively, compared to 3% knock-out efficiency without oligo). Combinations of donor oligo at 1.2µM with NH oligo, EE or NH+EE (final combined concentration of 2.4µM) further enhances knock-out efficiencies of the donor oligo (25%, 43% and 33%, respectively, compared to 11% knock-out efficiency with 1.2µM donor oligo alone), suggesting a carrier effect.

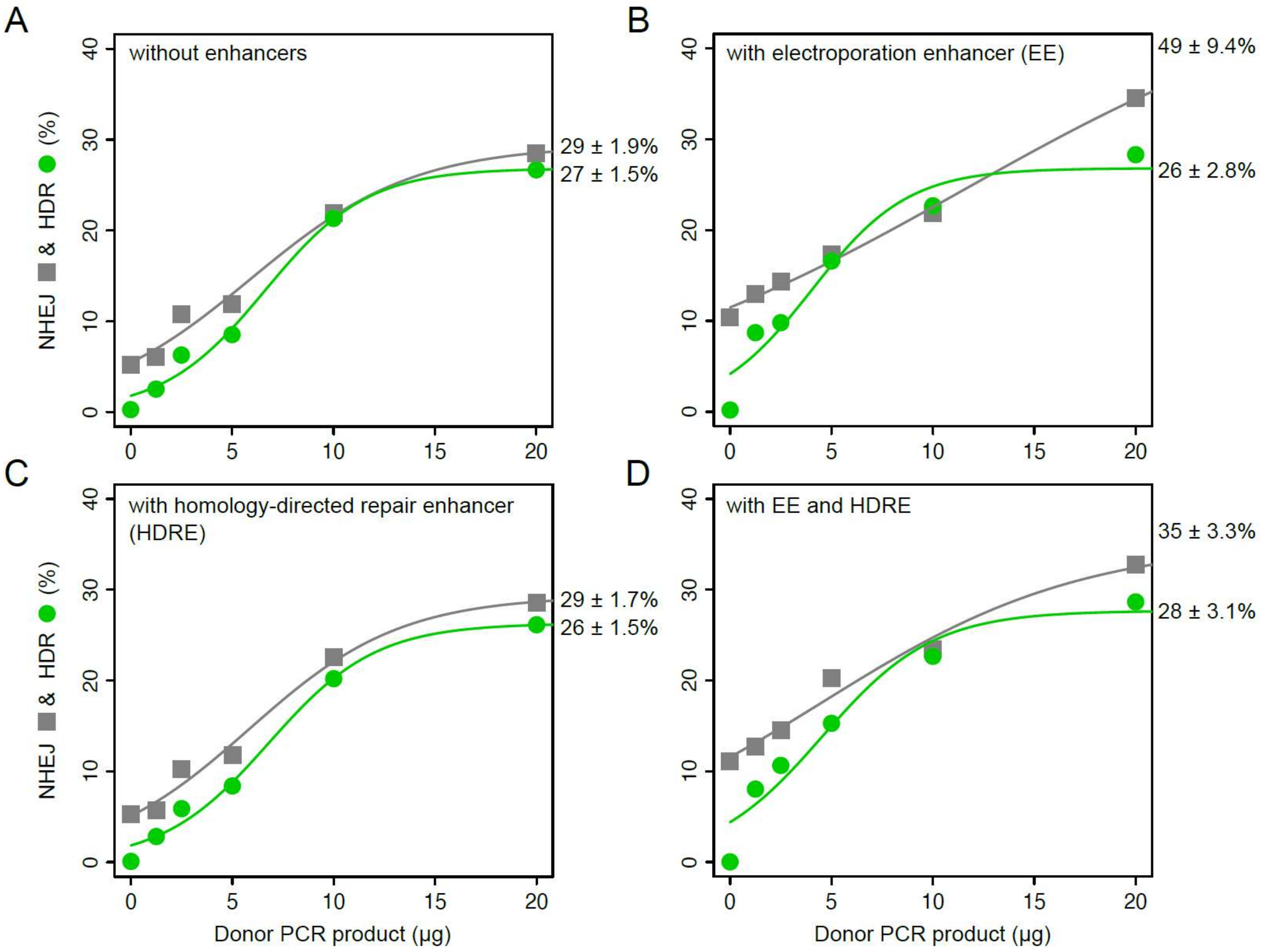

**Figure S7. CRISPR-Cas9 mediated knock-ins with electroporation enhancer (EE) and homology-directed repair enhancer (HDRE).** HDR (green) and NHEJ (grey) percentages (A) without enhancers, (B) with EE (final concentration at 1.2μM in the transfection mix), (C) with HDRE (final concentration at 1.2μM in growth media), and (D) with EE and HDRE. There was no effect of the HDRE on HDR and NHEJ probabilities, whereas EE increased NHEJ rates, consistent with a carrier effect. Percentages show maximum knock-out and knock-in efficiencies and standard errors estimated using a sigmoidal fit to the data. See also Figure 2 and Data S1 for data.

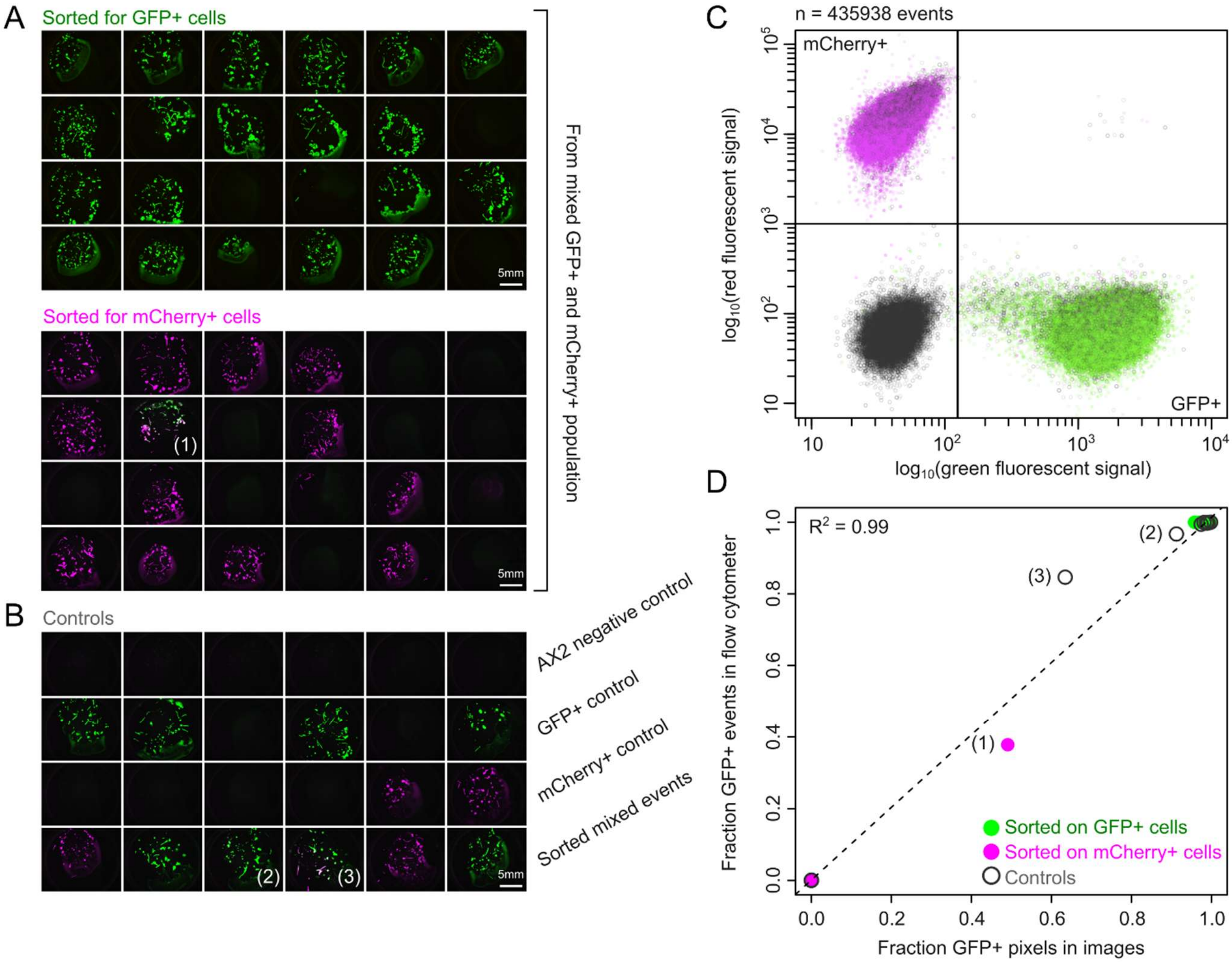

**Figure S8. Quantification of sorting accuracy using a combination of microscopy and flowcytometry.** (A) 24-well plates after sorting mixed population of *D. discoideum* AX2 mCherry+/GFP+ cells. Upper plate sorted for GFP+ cells and lower plate for mCherry+ cells. Plates are identical to those shown in Figure 4C-D. Cells were grown for 4 days after sorting, leaving sufficient time for cells to form plaques. (B) 24-well control plate where we sorted single cells from clonal populations of non-fluorescent *D. discoideum* AX2, *D. discoideum* AX2 *act15*::*gfp* or *D. discoideum* AX2 *act5::mCherry*, as negative controls, and mixed events (with GFP+ and mCherry+ signal) from a mixed population of *D. discoideum* AX2 *act15::gfp* and *D. discoideum* AX2 *act5::mCherry*, as a positive control for mixing. For each well, we also collected several sori and analyzed cells through flow cytometry to quantify percentage of GFP+ and mCherry+ cells. (C) Combined flow cytometry results from all wells: events from plate with GFP+ sorted cells shown in green, events from plate with mCherry+ sorted cells shown in magenta and events from control plate shown in grey. (D) Correlation between fraction of GFP+ pixels in microscopy images and GFP+ events in flowcytometry ($R^2 = 0.99, p < 0.001$). From the GFP+ and mCherry+ sorted plates only a single well showed mixed cells (1), while the other mixed wells, (2) and (3), were part of the positive controls in (B).

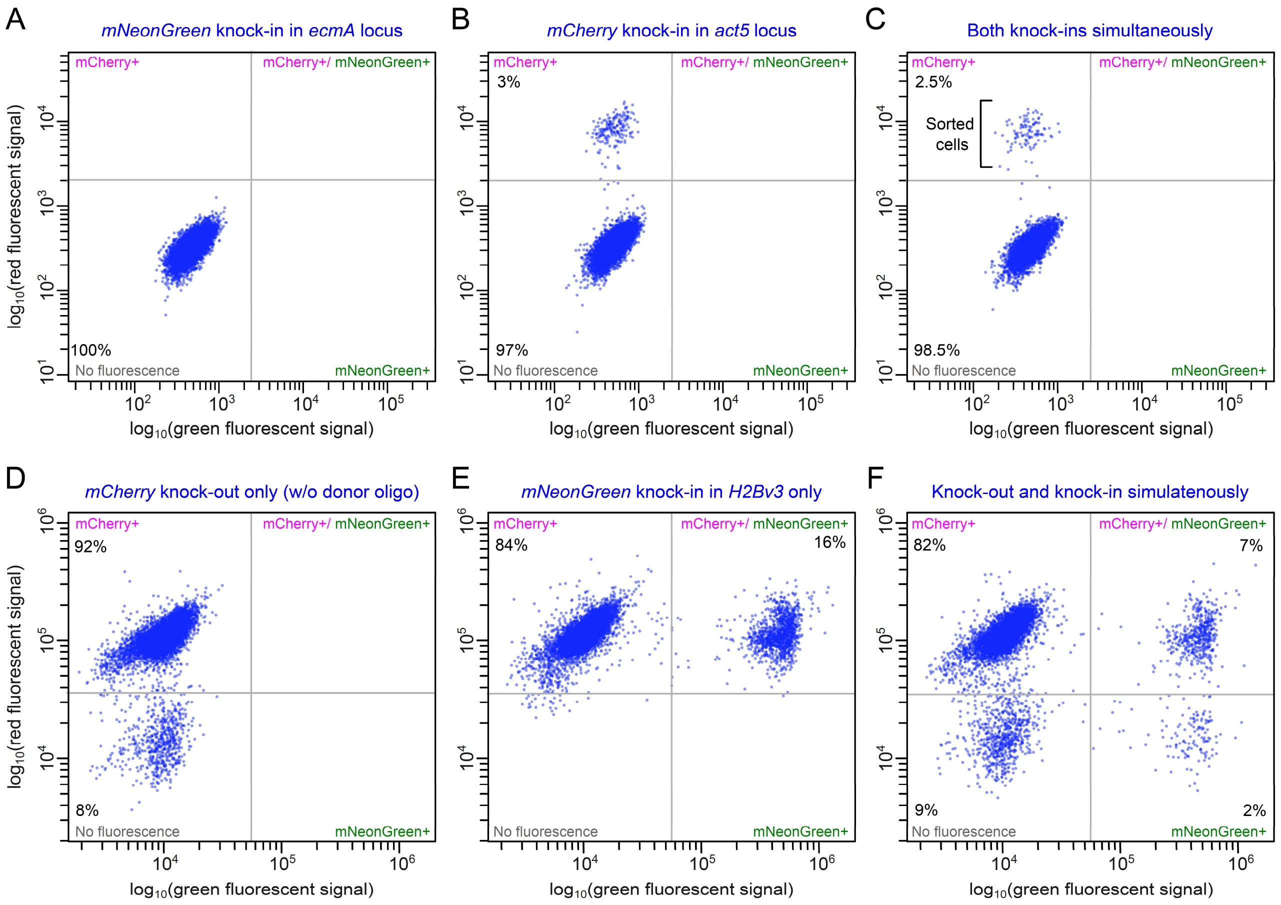

**Figure S9. CRISPR-Cas9 editing of multiple loci simultaneously**. (A)-(C) Simultaneous knock-ins in *D. discoideum* AX2 for *act5* (*act5*::*mCherry-P2A-act5*) and *ecmA* (*ecmA*::*mNeonGreen-P2A-ecmA*) expression reporters: Flow cytometry plots show results of (A) *ecmA* knock-in only, (B) *act5* knock-in only, and (C) double *act5* and *ecmA* knock-ins. We sorted for mCherry+ cells in the double knock-in condition, as *ecmA* is expressed in the slug stage only (i.e., no mNeonGreen+ cells are present during sorting), and screened with fluorescent microscopy for double knock-ins in sorted 24-well agar plates, which were confirmed by PCR. See Figure 5E for imaged slugs. (D)-(F) Simultaneous knock-out of *mCherry* (*act5*::*mCherry-*) and knock-in at *H2Bv3* (*H2Bv3*::*mNeonGreen-H2Bv3*) in *D. discoideum* AX2 *act5::mCherry* cells. Flow cytometry plots show results of (D) *mCherry* knock-out only, (E) *H2Bv3* knock-in only, and (F) a double knock-out and knock-in experiment. Cells with a knock-in had a significantly higher probability of also having a knock-out mutation: 22% (2 out of the 9% knock-in cells) instead of 10% (9 out of the 91% wild-type cells). Conversely, cells with a knock-out had a higher probability of also having a knock-in mutation: 18% (2 out of the 11% knock-out cells) instead of 8% (7 out of the 89% wild-type cells) (Fisher's Exact Test; $p < 10^{-10}$, odds ratio=2.3 with [1.8-2.8] 95%-confidence intervals, n=7223 cells).

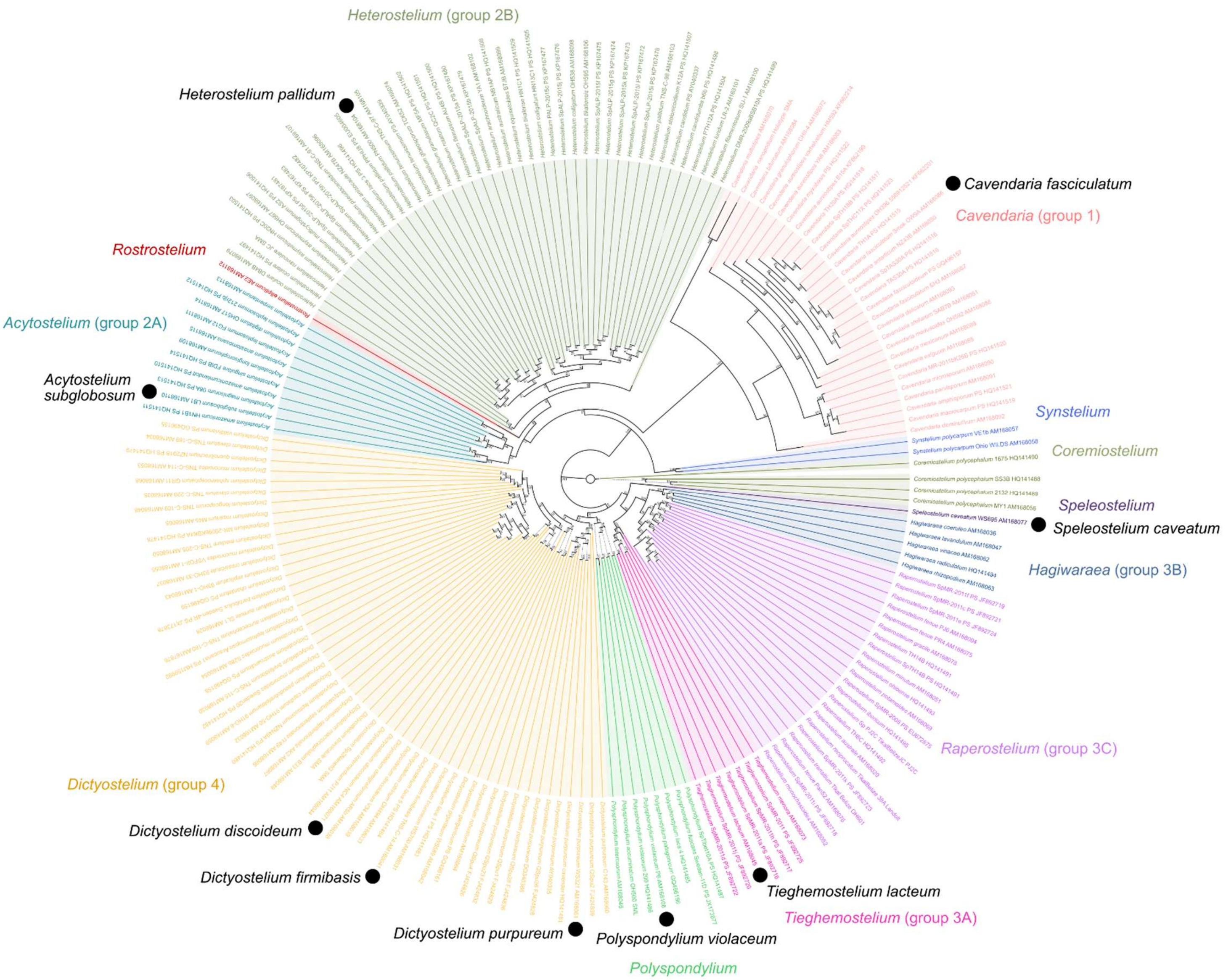

**Figure S10. Phylogenetic tree with representative *Dictyostelid* species.** Phylogenetic tree is based on multiple-sequence alignment of 18S rDNA of *Dictyostelids* as provided by (Sheikh et al. 2018). Colors in phylogenetic tree highlight *Dictyostelid* families and correspond to those in Figure 7A.

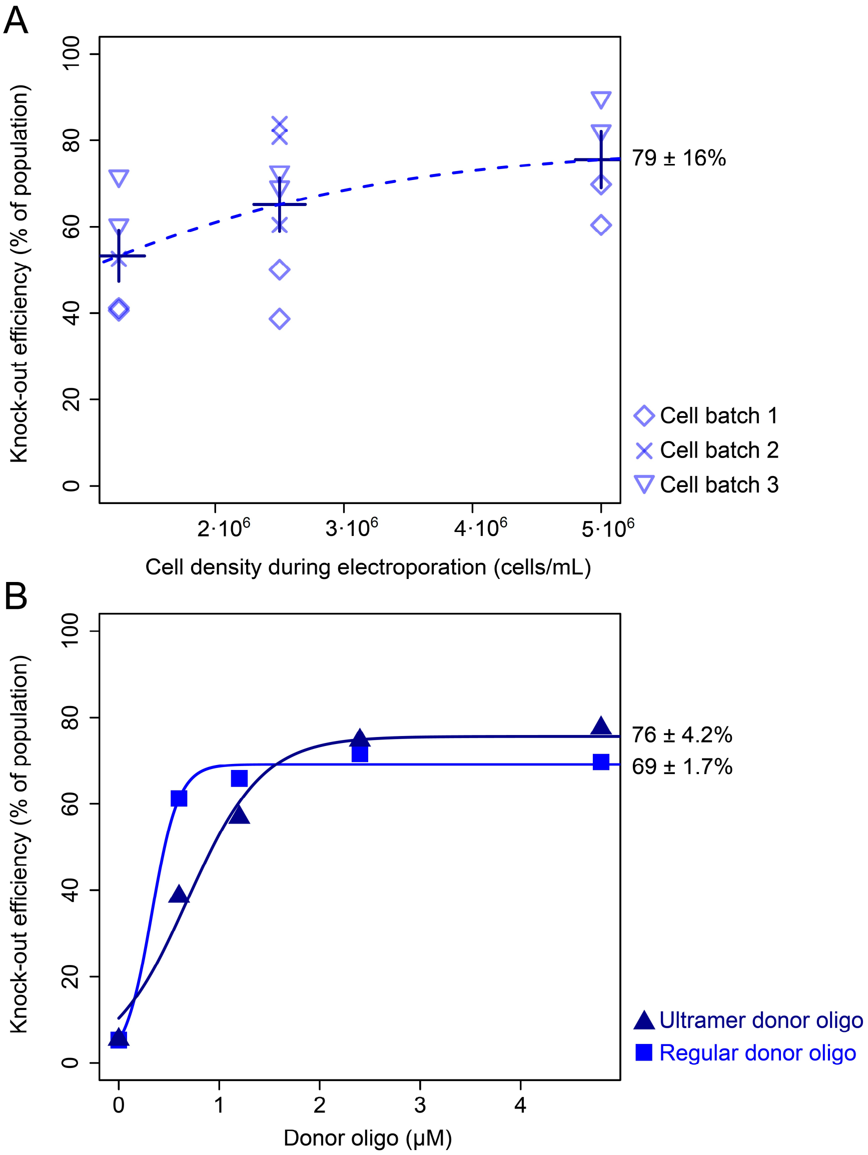

**Figure S11. Knock-out efficiency for different cell densities and donor oligo qualities.** (A) Knock-out efficiency using different cell densities per transfection for three batches of axenically-grown *D. discoideum* AX2 *act5*::*mCherry* cells. Dashed line indicates non-significant sigmoidal fit. (B) Knock-out efficiency of regular and ultramer oligos produced by IDT. Percentages show estimated maximum knock-out efficiencies and standard errors based on sigmoidal fits. See Data S1 for data.

#### Captions for Supplementary Files

All supplementary files (File S1-S6) are deposited to Zenodo: https://doi.org/10.5281/zenodo.15039721.

**File S1. Plasmid maps**. Snapgene files with plasmid maps for pMG005 with codon-optimized *mNeonGreen* and pMG008 with codon-optimized *mCherry*, as well as Snapgene files for pHO4d-Cas9, pET-FLAG-SpCas9-HF1 and pET-HiFi SpCas9-NLS-6xHis with SpyCas9 variants (see Figure S4B).

**File S2. Time-lapse of strains with *ecmA* and *act5* expression reporters**. Time-lapse video of feeding front and fruiting body development for (left) *D. discoideum* AX2, (middle) *ecmA* reporter and (right) *act5* reporter strains. *ecmA* reporter shows clear induction of *ecmA* expression at anterior part of slugs, while *act5* is expressed throughout the slugs.

**File S3. Raw image data of slugs in Figure 5**. Images show *ecmA* expression in slugs (z-stacks) and csv files show raw data for *ecmA* expression profiles along anterior-posterior axis of slugs.

**File S4. Raw image data of cells in Figure 6**. Images show maximal projection of *act5* expression in cells as well as cell masks and csv files provide raw pixel intensities from cell periphery to center.

**File S5. Videos of *Dictyotselid* species (low zoom)**. Overview time-lapse movies of WT and *act5*::*mNeonGreen-P2A-act5* knock-in strains of *D. discoideum*, *D. firmibasis*, *D. purpureum*, *P. violaceum*, *T. lacteum* and *H. pallidum*. For all species, growth of knock-in mutants (top) and WT (bottom) were similar: left panels show combined green fluorescence and spotlight image (using same color scale as in Figure 7); middle panels show green fluorescence only and right panels show spot lights only. Scale bars equal 5mm.

**File S6. Videos of *Dictyostelid* species (high zoom)**. Time-lapse movies at high magnification of *act5*::*mNeonGreen-P2A-act5* knock-in strains of *D. discoideum*, *D. firmibasis*, *D. purpureum*, *P. violaceum*, *T. lacteum* and *H. pallidum* (using same color scale as in Figure 7). Frames highlighted in Figure 7 are included as image files. Scale bars are equal to 1mm.

**Table S1**. Strain list.

| Species / strain | Resource ID | Resource |
|---|---|---|
| *Escherichia coli* B/r | DBS0305924 | dictyBase |
| *Dictyostelium discoideum* AX2 (Kay lab) | DBS0235521 | dictyBase |
| *Dictyostelium discoideum* AX2 *act15::gfp* | DBS0235536 | dictyBase |
| *Dictyostelium discoideum* AX2 *act5::mCherry, Hyg* | MG38 | This study[1] |
| *Dictyostelium discoideum* NC4 | DBS0304666 | dictyBase |
| *Dictyostelium discoideum* NC4 *act5::mCherry, Hyg* | HM1912 | (Paschke et al. 2018) |
| *Dictyostelium firmibasis* TNS-C-14 | DBS0235812 | dictyBase |
| *Dictyostelium purpureum* WS321 | DBS0235881 | dictyBase |
| *Polysphondylium violaceum* P6, S209 | DBS0236814 | dictyBase |
| *Tieghemostelium lacteum* S561 | DBS0235831 | dictyBase |
| *Acytostelium subglobosum* LB1 | DBS0235452 | dictyBase |
| *Heterostelium pallidum* PN500 | DBS0302501 | dictyBase |
| *Cavendaria fasciculatum* Swok OW9A | DBS0235811 | dictyBase |
| *Speleostelium caveatum* | DBS0235736 | dictyBase |

**Table S2.** List of generated knock-out strains.

| Identifier | Species / strain | Target gene / aa | Knock-out genotype (insertion size) | crRNA | Donor oligo | Confirmation PCR primers |
|---|---|---|---|---|---|---|
| MC1 | *D. discoideum* AX2 *act5::mCherry, Hyg* | *mCherry* / aa9 | *act5::mCherry* aa9 KO (random indel by NHEJ) | MC1 | NA | MR101/ MR103 |
| MC1_MR96 | *D. discoideum* AX2 *act5::mCherry, Hyg* | *mCherry* / aa9 | *act5::mCherry* aa9 KO (1bp) | MC1 | MR96 | MR101/ MR103 |
| MC1_MR97 | *D. discoideum* AX2 *act5::mCherry, Hyg* | *mCherry* / aa9 | *act5::mCherry* aa9 KO (1bp) | MC1 | MR97 | MR101/ MR103 |
| MC1_MR100 | *D. discoideum* AX2 *act5::mCherry, Hyg* | *mCherry* / aa9 | *act5::mCherry* aa9 KO (1bp) | MC1 | MR100 | MR101/ MR103 |
| MC1_MR124 | *D. discoideum* AX2 *act5::mCherry, Hyg* | *mCherry* / aa9 | *act5::mCherry* aa9 KO (37bp) | MC1 | MR124 | MR101/ MR103 |
| MC1_MR125 | *D. discoideum* AX2 *act5::mCherry, Hyg* | *mCherry* / aa9 | *act5::mCherry* aa9 KO (37bp) | MC1 | MR125 | MR101/ MR103 |
| MC1_MR127 | *D. discoideum* AX2 *act5::mCherry, Hyg* | *mCherry* / aa9 | *act5::mCherry* aa9 KO (37bp) | MC1 | MR127 | MR101/ MR103 |
| MC1_MR98 | *D. discoideum* AX2 *act5::mCherry, Hyg* | *mCherry* / aa9 | *act5::mCherry* aa9 KO (4bp) | MC1 | MR98 | MR101/ MR103 |
| MC2_MR131 | *D. discoideum* AX2 *act5::mCherry, Hyg* | *mCherry* / aa21 | *act5::mCherry* aa21 KO (37bp) | MC2 | MR131 | MR101 /MR103 |
| MC9_MR132 | *D. discoideum* AX2 *act5::mCherry, Hyg* | *mCherry* / aa27 | *act5::mCherry* aa27 KO (37bp) | MC9 | MR132 | MR101/ MR103 |
| MC1_MR125_ NC4 | *D. discoideum* NC4 *act5::mCherry, Hyg* | *mCherry* / aa9 | *act5::mCherry* aa9 KO (37bp) | MC1 | MR125 | MR101/ MR103 |

[1] Transfection of *D. discoideum* AX2 with pDM1514 following Paschke et al. (2018).

**Table S3.** Analysis of knock-out strains using Nanopore sequencing[2]

| Condition | Replicate | Description, sequence file with nanopore reads and alignment file[3] | Barcode | Overall coverage | % *mCherry* reads with correct edit (Alignment file to reference)[4] | % correct donor oligo integration (Alignment file to reference)[5] |
|---|---|---|---|---|---|---|
| Control 1 | 1 | *D. discoideum* AX2<br>**Sequence file**: Control_1.fastq.gz<br>**Alignment file to reference**: Control_1_to_reference.bam | 24 | 15.4x | NA | NA |
| Control 2 | 1 | *D. discoideum* AX2 *act5::mCherry*<br>**Sequence file**: Control_2.fastq.gz<br>**Alignment file to reference**: Control_2_to_reference.bam | 23 | 7.7x | 100% correct (no edits, 8 reads) (Control_2_to_mcherry.bam) | NA |
| Control 3 | 1 | Electroporated *D. discoideum* AX2 *act5::mCherry* cells, without RNP complex<br>**Sequence file**: Control_3_Rep1.fastq.gz<br>**Alignment file to reference**: Control_3_Rep1_to_reference.bam | 21 | 16.2x | 100% correct (no edits, 13 reads) (Control_3_Rep1_to_mcherry.bam) | NA |
| Control 3 | 2 | Electroporated *D. discoideum* AX2 *act5::mCherry* cells, without RNP complex<br>**Sequence file**: Control_3_Rep2.fastq.gz<br>**Alignment file to reference**: Control_3_Rep2_to_reference.bam | 22 | 18.0x | 100% correct (no edits, 12 reads) (Control_3_Rep2_to_mcherry.bam) | NA |
| Insert | 1 | *D. discoideum* AX2 *act5::mCherry* transfected with RNP complex only (crRNA: MC1), Sanger sequencing shows a 16bp insertion in *mCherry* produced by NHEJ<br>**Sequence file**: Insert.fastq.gz<br>**Alignment file to reference**: Insert_to_reference.bam | 16 | 15.1x | 100% correct (16bp insertion, 13 reads) (Insert_to_mcherry.bam) | NA |
| Deletion | 1 | *D. discoideum* AX2 *act5::mCherry* transfected with RNP complex only (crRNA: MC1), Sanger sequencing shows a 4bp deletion in *mCherry* produced by NHEJ<br>**Sequence file**: Deletion.fastq.gz<br>**Alignment file to reference**: Deletion_to_reference.bam | 17 | 25.8x | 100% correct (4bp deletion, 21 reads) (Deletion_to_mcherry.bam) | NA |
| Donor oligo | 1 | *D. discoideum* AX2 *act5::mCherry* transfected with RNP complex (crRNA: MC1) and donor oligo (MR125). Sanger sequencing shows 37bp insert from donor oligo.<br>**Sequence file**: Donor_oligo_Rep1.fastq.gz<br>**Alignment file to reference**: Donor_oligo_Rep1_to_reference.bam | 18 | 18.0x | 100% correct (37bp insertion, 9 reads) (Donor_oligo_Rep1_to_mcherry.bam) | 100% correct (5 reads) (Donor_oligo_Rep1_to_oligo.bam) |
| Donor oligo | 2 | *D. discoideum* AX2 *act5::mCherry* transfected with RNP complex (crRNA: MC1) and donor oligo (MR125). Sanger sequencing shows 37bp insert from donor oligo.<br>**Sequence file**: Donor_oligo_Rep2.fastq.gz<br>**Alignment file to reference**: Donor_oligo_Rep2_to_reference.bam | 19 | 15.0x | 100% correct (37bp insertion, 11 reads) (Donor_oligo_Rep2_to_mcherry.bam) | 100% correct (9 reads) (Donor_oligo_Rep2_to_oligo.bam) |
| Donor oligo | 3 | *D. discoideum* AX2 *act5::mCherry* transfected with RNP complex (crRNA: MC1) and donor oligo (MR125). Sanger sequencing shows 37bp insert from donor oligo.<br>**Sequence file**: Donor_oligo_Rep3.fastq.gz<br>**Alignment file to reference**: Donor_oligo_Rep3_to_reference.bam | 20 | 32.4x | 100% correct (37bp insertion, 24 reads) (Donor_oligo_Rep3_to_mcherry.bam) | 100% correct (10 reads) (Donor_oligo_Rep3_to_oligo.bam) |

[2] Based on Nanopore sequencing, no indication of increased off-target SNPs or indels due to CRISPR-Cas9 editing could be detected.
[3] All Nanopore sequencing files, alignment files and alignment reference (our *D. discoideum* AX2 assembly) are available on Zenodo (https://doi.org/10.5281/zenodo.15039721)
[4] *mCherry* sequence flanking the cutting site was queried against all nanopore reads using BLAST 2.15.0+ to confirm that all *mCherry* reads were edited as expected.
[5] 37nt donor oligo (without homology arms) was queried against all nanopore reads using BLAST 2.15.0+ to confirm that all donor oligos were integrated at the expected *mCherry* target site.

**Table S4.** List of generated knock-in strains.

| Identifier | Species / strain | Target gene / aa | Targeted locus tag[6] (GenBank assembly) | Knock-in genotype | crRNA | Donor PCR template | Donor PCR primers | Upstream junction PCR primers | Downstream junction PCR primer |
|---|---|---|---|---|---|---|---|---|---|
| MG060 | *D. discoideum* AX2 *act5::mCherry, Hyg* | *mCherry / aa9* | NA | *act5::mNeonGreen-P2A-mCherry* | MC1 | pMG005 | AP1169/ AP1170 | MR101/ MR113 | MR112/ MR103 |
| MG066 | *D. discoideum* NC4 | *act5 / aa1* | DDB_G0289663 (GCA_000004695.1) | *act5::mNeonGreen-act5* | MC5 | pMG005 | ME77/ MR78 | MR101/ MR113 | MR112/ MR102 |
| MG067 | *D. discoideum* AX2 | *act5 / aa1* | DDB_G0289663 (GCA_000004695.1) | *act5:mNeonGreen-act5* | MC5 | pMG005 | ME77/ MR78 | MR101/ MR113 | MR112/ MR102 |
| NA[7] | *D. discoideum* AX2 | *H2Bv3 / aa4* | DDB_G0286509 (GCA_000004695.1) | *H2Bv3::mNeonGreen-H2Bv3* | MC3 | pMG005 | MR75/ ME76 | MR85/ MR113 | MR112/ MR86 |
| NA | *D. discoideum* AX2 | *H2Bv3 / aa155* | DDB_G0286509 (GCA_000004695.1) | *H2Bv3::H2Bv3-mNeonGreen* | MC4 | pMG005 | MR73/ MR74 | MR89/ MR113 | MR112/ MR88 |
| MG077 | *D. discoideum* AX2 | *ecmA / aa27* | DDB_G0277853 (GCA_000004695.1) | *ecmA::mNeonGreen-P2A-ecmA* | MC10 | pMG005 | MR133/ MR134 | MR146/ MR113 | |
| MG075 | *D. discoideum* AX2 | *act5 / aa1* | DDB_G0289663 (GCA_000004695.1) | *act5::mNeonGreen-P2A-act5* | MC5 | pMG005 | MR77/ MR80 | MR101/ MR113 | MR112/ MR102 |
| MG087 | *D. discoideum* NC4 | *act5 / aa1* | DDB_G0289663 (GCA_000004695.1) | *act5::mNeonGreen-P2A-act5* | MC5 | pMG005 | MR77/ MR80 | MR101/ MR113 | MR112/ MR102 |
| MG137 | *D. discoideum* AX2 | *act5 / aa1, ecmA / aa27* | DDB_G0289663, DDB_G0277853 (GCA_000004695.1) | *act5::mCherry-*P2A-*act5*, *ecmA::mNeonGreen-*P2A-*ecmA* | MC5, MC10 | pMG008, pMG005 | MR169/ MR80, MR133/ MR134 | MR101/ MR103, MR146/ MR113 | MR32/ MR102 |
| MG084 | *P. violaceum* P6, S209 | *Act /aa12* | CYY_010281 (GCA_000277445.1) | *act::mNeonGreen-P2A-act* | MC31 | pMG005 | MR149/ MR150 | MR193/ MR113 | MR112/ MR157 |
| MG100 | *D. firmibasis* TNS-C-14 | *act / aa1* | RB653_008742 (GCA_036169595.1) | *act::mNeonGreen-P2A-act* | MC12 | pMG005 | MR147/ MR148 | MR153/ MR113 | MR112/ MR156 |
| MG130 | *H. pallidum* PN500 | *act* | PPL_06386 (GCA_000004825.1) | *act::mNeonGreen-P2A-act* | MC14 | pMG005 | MR151/ MR152 | MR155/ MR113 | MR112/ MR158 |
| NA | *A. subglobosum* LB1 | *act* | SAMD00019534_067160 (GCA_000787575.2) | *act::mNeonGreen-P2A-act* | MC16 | pMG005 | MR174/ MR175 | NA | NA |
| MG116 | *T. lacteum* 561 | *act / aa13* | DLAC_02274 (GCA_001606155.1) | *act::mNeonGreen-P2A-act* | MC18 | pMG005 | MR172/ MR173 | | MR112/ MR156 |
| MG129 | *D. purpureum* WS321 | *act / aa12* | DICPUDRAFT_92254 (GCA_000190715.1) | *act::mNeonGreen-P2A-act* | MC13 | pMG005 | MR149/ MR150 | MR154/ MR113 | MR112/ MR157 |

[6] Correct integration of the donor DNA product at the targeted locus was confirmed by PCR with primers specific for the targeted locus and sanger sequencing.
[7] No identifier associated, because *D. discoideum* AX2 *H2Bv3*::*H2Bv3-mNeonGreen*, *D. discoideum* AX2 *H2Bv3*::*mNeonGreen-H2Bv3* and *A. subglobosum act*::*mNeonGreen-P2A-act* strains could not be propagated due to deleterious growth effects.

**Table S5**. List of crRNAs.

| Identifiers | Spacer sequence | Target: Strain / gene / aa |
|---|---|---|
| MC1 | AAAAGGTGAAGAAGATAATA | *D. discoideum* / *mCherry* / aa9 |
| MC2 | TATGAGATTTAAAGTTCATA | *D. discoideum* / *mCherry* / a21 |
| MC9 | CATATGGAAGGTTCAGTTAA | *D. discoideum* / *mCherry* / aa27 |
| MC5 | ATTATATATAAAAAAAATGGA | *D. discoideum* / *act5* / aa1 |
| MC3 | TTCAAAATGGTATTCGTTAA | *D. discoideum* / *H2Bv3* / aa4 |
| MC10 | ACATGCTGTATTGGTTTGTA | *D. discoideum* / *ecmA* / aa27 |
| MC4 | ACTGAAAGCAAAAACTAAAT | *D. discoideum* / *H2Bv3* / aa155 |
| MC12 | AAAATACAATAAAAAAATGGA | *D. firmibasis* / *act* / aa1 |
| MC13 | CAAGCTTTAGTTATCGATAA | *D. purpureum* / *act* / aa12 |
| MC14 | AGGTGAAGACGTTCAAGCTT | *H. pallidum* / *act* / aa8 |
| MC16 | AGGAGAAGACGTTCAAGCTT | *A. subglosum* / *act5* / aa7 |
| MC18 | TGATGGGAATACAGCTCTTG | *T. lacteum* / *act* / aa13 |
| MC31 | CAAGCTTTAGTTATCGATAA | *P. violaceum* / *act* / aa12 |

**Table S6**. List of oligos (homology arms in blue).

| Identifier | Sequence | Description |
|---|---|---|
| MR96 | AAGGTGAAGAAGATTAATATGGCAATTAT | Donor oligo. Target: *mCherry* aa9. Insertion: 1bp. Homology arms: 14bp. |
| MR97 | AAAAATGGTTTCAAAAGGTGAAGAAGATTAA TATGGCAATTATTAAAGAATTTATGA | Donor oligo. Target: *mCherry* aa9. Insertion: 1bp. Homology arms: 28bp. |
| MR100 | AAAATAAATTATATATAAAAAAGATCCAAAA AATGGTTTCAAAAGGTGAAGAAGATTAATAT GGCAATTATTAAAGAATTTATGAGATTTAAA GTTCATATGGAAGGTTCAGT | Donor oligo. Target: *mCherry* aa9. Insertion: 1bp. Homology arms: 56bp. |
| MR124 | AAGGTGAAGAAGATTACCCCTACGACGTGCC AGATTACGCCTCTCTGTAATAATATGGCAAT TAT | Donor oligo. Target: *mCherry* aa9. Insertion: 37bp. Homology arms: 14bp. |
| MR125 | AAAAATGGTTTCAAAAGGTGAAGAAGATTAC CCCTACGACGTGCCAGATTACGCCTCTCTGT AATAATATGGCAATTATTAAAGAATTTATGA | Donor oligo. Target: *mCherry* aa9. Insertion: 37bp. Homology arms: 28bp. |
| MR127 | AAAATAAATTATATATAAAAAAGATCCAAAA AATGGTTTCAAAAGGTGAAGAAGATTACCCC TACGACGTGCCAGATTACGCCTCTCTGTAAT AATATGGCAATTATTAAAGAATTTATGAGAT TTAAAGTTCATATGGAAGGTTCAGT | Donor oligo. Target: *mCherry* aa9. Insertion: 37bp. Homology arms: 56bp. |
| MR98 | AAAAATGGTTTCAAAAGGTGAAGAAGATTAA TAATATGGCAATTATTAAAGAATTTATGA | Donor oligo. Target: *mCherry* aa9. Insertion: 4bp. Homology arms: 28bp. |
| MR131 | TATTAAAGAATTTATGAGATTTAAAGTTTAC CCCTACGACGTGCCAGATTACGCCTCTCTGT AATAATATGGAAGGTTCAGTTAATGGTCATG A | Donor oligo. Target: *mCherry* aa21. Insertion: 37bp. Homology arms: 28bp. |
| MR132 | GAGATTTAAAGTTCATATGGAAGGTTCATAC CCCTACGACGTGCCAGATTACGCCTCTCTGT AATGTTAATGGTCATGAATTTGAAATTGAAG | Donor oligo. Target: *mCherry* aa27. Insertion: 37bp. Homology arms: 28bp. |
| AP1169 | GATCCAAAAAATGGTTTCAAAAGGTGAAGAA GATGTATCTAAGGGAGAAGAAG | Forward primer for *act5*::*mNeonGreen-P2A-mCherry* knock-in. Template: pMG005. |
| AP1170 | CTTTAAATCTCATAAATTCTTTAATAATTGC CATATTAGAACTAGATCCACTAGGTC | Reverse primer for *act5*::*mNeonGreen-P2A-mCherry* knock-in. Template: pMG005. |
| MR73 | CTGTCAACAAGTACAATCCAACTGAAAGCAA AAACGACTACAAAGATCACGATG | Forward primer for *H2Bv3*::*mNeonGreen-H2Bv3* knock-in. Template: pMG005. |
| MR74 | TTTTTAAGGAATATAGTTCATTTGGAACCAA TTTACTTATATAATTCATCCATTCC | Reverse primer for *H2Bv3*::*mNeonGreen-H2Bv3* knock-in. Template: pMG005. |
| MR75 | AATAGTCAATTAATATAATTCAAAATGGTAT TCGTATCTAAGGGAGAAGAAG | Forward primer for *H2Bv3*::*H2Bv3-mNeonGreen* knock-in. Template: pMG005. |
| ME76 | TGAGTTGAACCTTTGGTTGCTTTCTTTTGAC CTTTAACACCAGAACCACCAGCATAA | Reverse primer for *H2Bv3*::*H2Bv3-mNeonGreen* knock-in. Template: pMG005. |
| ME77 | ATAAAAACTTAAAATAAATTATATATAAAAA AATGGTATCTAAGGGAGAAGAAG | Forward primer for *act5*::*mNeonGreen-act5* and *act5*::*mNeonGreen-P2A-act5* knock-in. Template: pMG005. |
| MR169 | ATAAAAACTTAAAATAAATTATATATAAAAA AATGGTTTCAAAAGGTGAAGAAG | Forward primer for *act5*::*mCherry-P2A-act5* knock-in. Template: pMG008 |
| MR80 | TTATCGATAACTAAAGCTTGAACATCTTCAC CGTCAGAACTAGATCCACTAGGTC | Reverse primer for *act5*::*mNeonGreen-P2A-act5* and *act5*::*mCherry-P2A-act5* knock-in. Template: pMG005 and pMG008, respectively. |
| MR78 | TTATCGATAACTAAAGCTTGAACATCTTCAC CGTCACCAGAACCACCAGCATAA | Reverse primer for *act5*::*mNeonGreen-act5* knock-in. Template: pMG005. |
| MR133 | TAATATTTAATAGCGGAAACTGAAACCATAG TATCTAAGGGAGAAGAAG | Forward primer for *ecmA*::*mNeonGreen-P2A-ecmA* knock-in. Template: pMG005. |
| MR134 | ATCATCACAATCACATGCTGTATTGGTTTGA GAACTAGATCCACTAGGTC | Reverse primer for *ecmA*::*mNeonGreen-P2A-ecmA* knock-in. Template: pMG005. |
| MR147 | ATTTCAAACTATAATATATAAAATACAATAA AAAATGGTATCTAAGGGAGAAGAAG | Forward primer for *D. firmibasis act*::*mNeonGreen-P2A-act* knock-in. Template: pMG005. |
| MR148 | TATCGATAACTAAAGCTTGAACATCTTCACC TTCAGAACTAGATCCACTAGGTC | Reverse primer for *D. firmibasis act*::*mNeonGreen-P2A-act* knock-in. Template: pMG005. |
| MR149 | AGATGGAAGGTGAAGATGTTCAAGCTTTAGT TATCGTATCTAAGGGAGAAGAAG | Forward primer for *D. purpureum act*::*mNeonGreen-P2A-act* knock-in. Template: pMG005. |
| MR150 | GCAAAACCGGCTTTGCACATACCTGAACCGT TATCAGAACTAGATCCACTAGGTC | Reverse primer for *D. purpureum act*::*mNeonGreen-P2A-act* knock-in. Template: pMG005. |
| MR151 | ATAATAAATAAACAATGGAAGGTGAAGACGT TCAAGTATCTAAGGGAGAAGAAG | Forward primer for *H. pallidum act*::*mNeonGreen-P2A-act* knock-in. Template: pMG005. |
| MR152 | CTAGATGATCAACATACGTTATCAATTACCA AAGCAGAACTAGATCCACTAGGTC | Reverse primer for *H. pallidum act*::*mNeonGreen-P2A-act* knock-in. Template: pMG005. |

| | | |
|---|---|---|
| MR172 | TGTGTAAAGCCGGTTTTGCTGGTGATGATGCCCCAGTATCTAAGGGAGAAGAAG | Forward primer for *T. lacteum act*::*mNeonGreen-P2A-act* knock-in. Template: pMG005. |
| MR173 | CTTGGACGACCGACAATTGATGGGAATACAGCTCTAGAACTAGATCCACTAGGTC | Reverse primer for *T. lacteum act*::*mNeonGreen-P2A-act* knock-in. Template: pMG005. |
| MR174 | TAATAATAATAATAATGGAAGGAGAAGACGTTCAAGTATCTAAGGGAGAAGAAG | Forward primer for *A. subglobosum act*::*mNeonGreen-P2A-act* knock-in. Template: pMG005. |
| MR175 | TTGCACATACCGGAACCGTTGTCAATGACCAAAGCAGAACTAGATCCACTAGGTC | Reverse primer for *A. subglobosum act*::*mNeonGreen-P2A-act* knock-in. Template: pMG005. |
| MR63 | GTTCCTTCCACCGGATACTTG | Reverse primer confirmation PCR. Target: downstream of *act5* |
| MR129 | TACCCCTACGACGTGCCA | Forward primer confirmation PCR. Target: *HA* |
| MR128 | TTACAGAGAGGCGTAATCTGG | Reverse primer confirmation PCR. Target: *HA* |
| MR103 | TCCCATGCAAATGGTAATGGAC | Reverse primer confirmation PCR. Target: *mCherry* |
| MR32 | ATGGGTTGGGAAGCATCATCA | Forward primer confirmation PCR. Target: *mCherry* |
| MR112 | CCAAGCCAATGGCAGCAAAT | Forward primer confirmation PCR. Target: *mNeonGreen* |
| MR113 | AGCGGCTTGAAATGGACTCA | Reverse primer confirmation PCR. Target: *mNeonGreen* |
| MR101 | CACCAGGTATATTACCAGATGGC | Forward primer confirmation PCR. Target: upstream of *act5* |
| MR89 | TCATTCTCACTGGTGAGTTAGC | Forward primer confirmation PCR. Target: *H2Bv3* |
| MR86 | GGTGGTTGAAGCGGTTTTCTC | Reverse primer confirmation PCR. Target: *H2Bv3* |
| MR88 | CATCCCAATCGATGGCATTCA | Reverse primer confirmation PCR. Target: downstream of *H2Bv3* |
| MR146 | TGTAAACTCGTGAGGGGGTTGG | Forward primer confirmation PCR. Target: upstream of *ecmA* |
| MR153 | GCCAACCGATATTGATAACGACC | Forward primer confirmation PCR. Target: upstream of *D. firmibasis act*. |
| MR154 | CAGGTTTTGAAAGTTGGGAAGCC | Forward primer confirmation PCR. Target: upstream of *D. purpureum act*. |
| MR155 | TGAGCAATCGATTGGAGTTGG | Forward primer confirmation PCR. Target: upstream of *H. pallidum act*. |
| MR156 | GAGTCTTTTTGACCCATACCGACC | Reverse primer confirmation PCR. Target: *D. firmibasis act*. |
| MR157 | CTCTTTGATTGGGCTTCATCACC | Forward primer confirmation PCR. Target: *D. purpureum act*. |
| MR158 | GTCTTGGACGACCGACGATC | Reverse primer confirmation PCR. Target *H. pallidum act*. |
| 18S_D142F | TGGATAACCGCAGTAAACGGG | 1st round of PCR for 18S sequencing[8] |
| 18SR_B | TGATCCTTCTGCAGGTTCAC | 1st round of PCR for 18S sequencing[2] |
| 18S_D307F | GTTTGGCCTACCATGGTTGTAA | 2nd round of PCR for 18S sequencing[2] |
| 18S_D862R | CACCTCTCGCCCCAATATGA | 2nd round of PCR for 18S sequencing[2] |
| MR44 | GATCCAAAAAAATGGGAGTAGCTGATTTAATAAAAAAATTTGAATCAATATCTAAAGAAGA | Non-homologous oligo (Figure S6) |

[8] Baldauft et al. 2018

**Text S1**. Vector sequences.

>pMG005 (kozak like-Start Codon-SV40-NLS-Linker3-nuleoplasmin NLS- 3xFLAG-tag-Codon optimized mNeonGreen-HA-tag-Linker1-P2A-Linker2B-Stop Codon-act8T)

CTAAATTGTAAGCGTTAATATTTTGTTAAAATTCGCGTTAAATTTTTGTTAAATCAGCTCATTTTTTAACCAATAGGCCGAAATCGGCAA
AATCCCTTATAAATCAAAAGAATAGACCGAGATAGGGTTGAGTGGCCGCTACAGGGCGCTCCCATTCGCCATTCAGGCTGCGCAACTGTT
GGGAAGGGCGTTTCGGTGCGGGCCTCTTCGCTATTACGCCAGCTGGCGAAAGGGGGATGTGCTGCAAGGCGATTAAGTTGGGTAACGCCA
GGGTTTTCCCAGTCACGACGTTGTAAAACGACGGCCAGTGAGCGCGACGTAATACGACTCACTATAGGGCGAATTGGCGGAAGGCCGTCA
AGGCCGCATAAAAAATGGTGAGTTCATTAAGACCACCAAAGAAAAAGAGAAAGGTTTGTCCAGGTGATAGATGGAGTAGTACAGGAGGTG
GAAGAAGTAGAACTTCAAAGAGACCTGCCGCTACAAAAAAGGCCGGACAAGCAAAGAAGAAAAAGGACTACAAAGATCACGATGGAGATT
ATAAGGATCACGATATAGATTACAAAGATGATGATGATAAAGTATCTAAGGGAGAAGAAGATAATATGGCCTCTCTTCCTGCCACCCATG
AATTACACATTTTCGGTTCTATTAATGGAGTTGATTTCGATATGGTTGGACAAGGAACTGGTAATCCAAATGATGGTTATGAGGAATTAA
ATCTTAAAAGTACAAAGGGAGATCTTCAATTTAGTCCATGGATCTTAGTACCTCACATCGGATACGGTTTTCACCAATACTTGCCATACC
CAGATGGTATGAGTCCATTTCAAGCCGCTATGGTAGATGGAAGTGGTTACCAAGTACACCGTACAATGCAATTTGAAGATGGAGCTAGTC
TTACAGTTAATTACCGTTATACATACGAGGGATCTCATATCAAAGGTGAGGCCCAAGTAAAGGGAACCGGTTTCCCTGCAGATGGACCAG
TAATGACTAATAGTTTAACTGCCGCCGATTGGTGTAGATCAAAAAAGACATATCCAAATGATAAGACTATAATATCAACATTCAAATGGT
CTTACACTACTGGAAATGGTAAGAGATATCGTTCTACCGCCAGAACAACTTACACATTTGCCAAGCCAATGGCAGCAAATTACTTAAAGA
ATCAACCTATGTACGTTTTTCGTAAAACAGAGTTGAAGCACTCAAAGACCGAGTTGAATTTTAAAGAGTGGCAAAAGGCATTTACAGATG
TTATGGGAATGGATGAATTATATAAGTACCCATACGATGTACCTGATTATGCTGGTGGTTCTGGTGCTACAAATTTCTCTTTGTTAAAGC
AAGCTGGAGATGTTGAAGAAAATCCTGGACCTTCTAGTGGACCTAGTGGATCTAGTTCTTGAATTATTTAATAAATAATAAAAAAACAAA
TTGTTGTAATAATCTAATATTTTCTTTTTTTTTTAATTTTTTTTTTTTTAAATCTTAATAATTATTAAGTTATTTTAATTTTTTTTTTTTT
TTTTTTTTTTTTTTTTTTTTTTTTCTATCAAAAAAATCAAATATATTTAAAAAATTTATTATTTACAGATACATTTTGAATGGTGAAGA
TAAATATATGCATTAGATGTAAAACAGCCAAAGAGTATGAAAATCAAAAAGATACTGGGCCTCATGGGCCTTCCGCTCACTGCCCGCTTT
CCAGTCGGGAAACCTGTCGTGCCAGCTGCATTAACATGGTCATAGCTGTTTCCTTGCGTATTGGGCGCTCTCCGCTTCCTCGCTCACTGA
CTCGCTGCGCTCGGTCGTTCGGGTAAAGCCTGGGGTGCCTAATGAGCAAAAGGCCAGCAAAAGGCCAGGAACCGTAAAAAGGCCGCGTTG
CTGGCGTTTTTCCATAGGCTCCGCCCCCCTGACGAGCATCACAAAAATCGACGCTCAAGTCAGAGGTGGCGAAACCCGACAGGACTATAA
AGATACCAGGCGTTTCCCCCTGGAAGCTCCCTCGTGCGCTCTCCTGTTCCGACCCTGCCGCTTACCGGATACCTGTCCGCCTTTCTCCCT
TCGGGAAGCGTGGCGCTTTCTCATAGCTCACGCTGTAGGTATCTCAGTTCGGTGTAGGTCGTTCGCTCCAAGCTGGGCTGTGTGCACGAA
CCCCCCGTTCAGCCCGACCGCTGCGCCTTATCCGGTAACTATCGTCTTGAGTCCAACCCGGTAAGACACGACTTATCGCCACTGGCAGCA
GCCACTGGTAACAGGATTAGCAGAGCGAGGTATGTAGGCGGTGCTACAGAGTTCTTGAAGTGGTGGCCTAACTACGGCTACACTAGAAGA
ACAGTATTTGGTATCTGCGCTCTGCTGAAGCCAGTTACCTTCGGAAAAAGAGTTGGTAGCTCTTGATCCGGCAAACAAACCACCGCTGGT
AGCGGTGGTTTTTTTGTTTGCAAGCAGCAGATTACGCGCAGAAAAAAAGGATCTCAAGAAGATCCTTTGATCTTTTCTACGGGGTCTGAC
GCTCAGTGGAACGAAAACTCACGTTAAGGGATTTTGGTCATGAGATTATCAAAAAGGATCTTCACCTAGATCCTTTTAAATTAAAAATGA
AGTTTTAAATCAATCTAAAGTATATATGAGTAAACTTGGTCTGACAGTTACCAATGCTTAATCAGTGAGGCACCTATCTCAGCGATCTGT
CTATTTCGTTCATCCATAGTTGCCTGACTCCCCGTCGTGTAGATAACTACGATACGGGAGGGCTTACCATCTGGCCCCAGTGCTGCAATG
ATACCGCGAGAACCACGCTCACCGGCTCCAGATTTATCAGCAATAAACCAGCCAGCCGGAAGGGCCGAGCGCAGAAGTGGTCCTGCAACT
TTATCCGCCTCCATCCAGTCTATTAATTGTTGCCGGGAAGCTAGAGTAAGTAGTTCGCCAGTTAATAGTTTGCGCAACGTTGTTGCCATT
GCTACAGGCATCGTGGTGTCACGCTCGTCGTTTGGTATGGCTTCATTCAGCTCCGGTTCCCAACGATCAAGGCGAGTTACATGATCCCCC
ATGTTGTGCAAAAAAGCGGTTAGCTCCTTCGGTCCTCCGATCGTTGTCAGAAGTAAGTTGGCCGCAGTGTTATCACTCATGGTTATGGCA
GCACTGCATAATTCTCTTACTGTCATGCCATCCGTAAGATGCTTTTCTGTGACTGGTGAGTACTCAACCAAGTCATTCTGAGAATAGTGT
ATGCGGCGACCGAGTTGCTCTTGCCCGGCGTCAATACGGGATAATACCGCGCCACATAGCAGAACTTTAAAAGTGCTCATCATTGGAAAA
CGTTCTTCGGGGCGAAAACTCTCAAGGATCTTACCGCTGTTGAGATCCAGTTCGATGTAACCCACTCGTGCACCCAACTGATCTTCAGCA
TCTTTTACTTTCACCAGCGTTTCTGGGTGAGCAAAAACAGGAAGGCAAAATGCCGCAAAAAAGGGAATAAGGGCGACACGGAAATGTTGA
ATACTCATACTCTTCCTTTTTCAATATTATTGAAGCATTTATCAGGGTTATTGTCTCATGAGCGGATACATATTTGAATGTATTTAGAAA
AATAAACAAATAGGGGTTCCGCGCACATTTCCCCGAAAAGTGCCAC

>pMG008 (kozak like-Start Codon-SV40-NLS-Linker3-nuleoplasmin NLS- 3xFLAG-tag-Codon optimized mCherry-HA-tag-Linker1-P2A-Linker2B-Stop Codon-act8T)

CTAAATTGTAAGCGTTAATATTTTGTTAAAATTCGCGTTAAATTTTTGTTAAATCAGCTCATTTTTTAACCAATAGGCCGAAATCGGCAA
AATCCCTTATAAATCAAAAGAATAGACCGAGATAGGGTTGAGTGGCCGCTACAGGGCGCTCCCATTCGCCATTCAGGCTGCGCAACTGTT
GGGAAGGGCGTTTCGGTGCGGGCCTCTTCGCTATTACGCCAGCTGGCGAAAGGGGGATGTGCTGCAAGGCGATTAAGTTGGGTAACGCCA
GGGTTTTCCCAGTCACGACGTTGTAAAACGACGGCCAGTGAGCGCGACGTAATACGACTCACTATAGGGCGAATTGGCGGAAGGCCGTCA
AGGCCGCATAAAAAATGGTGAGTTCATTAAGACCACCAAAGAAAAAGAGAAAGGTTTGTCCAGGTGATAGATGGAGTAGTACAGGAGGTG
GAAGAAGTAGAACTTCAAAGAGACCTGCCGCTACAAAAAAGGCCGGACAAGCAAAGAAGAAAAAGGACTACAAAGATCACGATGGAGATT
ATAAGGATCACGATATAGATTACAAAGATGATGATGATAAAGTTTCAAAAGGTGAAGAAGATAATATGGCAATTATTAAAGAATTTATGA
GATTTAAAGTTCATATGGAAGGTTCAGTTAATGGTCATGAATTTGAAATTGAAGGTGAAGGTGAAGGTAGACCATATGAAGGTACACAAA
CAGCAAAATTAAAAGTTACAAAAGGTGGTCCATTACCATTTGCATGGGATATTTTATCACCACAATTTATGTATGGTTCAAAAGCATATG
TTAAACATCCAGCAGATATTCCAGATTATTTAAAATTATCATTTCCAGAAGGTTTTAAATGGGAAAGAGTTATGAATTTTGAAGATGGTG
GTGTTGTTACAGTTACACAAGATTCATCATTACAAGATGGTGAATTTATTTATAAAGTTAAATTAAGAGGTACAAATTTTCCATCAGATG
GTCCAGTTATGCAAAAGAAAACAATGGGTTGGGAAGCATCATCAGAAAGAATGTATCCAGAAGATGGTGCATTAAAAGGTGAAATTAAAC
AAAGATTAAAATTAAAAGATGGTGGTCATTATGATGCAGAAGTTAAAACAACATATAAAGCAAAAAAACCAGTTCAATTACCAGGTGCAT
ATAATGTTAATATTAAATTAGATATTACATCACATAATGAAGATTATACAATTGTTGAACAATATGAAAGAGCAGAAGGTAGACATTCAA
CAGGTGGTATGGATGAATTATATAAATACCCATACGATGTACCTGATTATGCTGGTGGTTCTGGTGCTACAAATTTCTCTTTGTTAAAGC
AAGCTGGAGATGTTGAAGAAAATCCTGGACCTTCTAGTGGACCTAGTGGATCTAGTTCTTGAATTATTTAATAAATAATAAAAAAAACAAA
TTGTTGTAATAATCTAATATTTTCTTTTTTTTTTAATTTTTTTTTTTTAAATCTTAATAATTATTAAGTTATTTTAATTTTTTTTTTTTT
TTTTTTTTTTTTTTTTTTTTTTTTTCTATCAAAAAAATCAAATATATTTAAAAAATTTATTATTTACAGATACATTTTGAATGGTGAAGA
TAAATATATGCATTAGATGTAAAACAGCCAAAGAGTATGAAAATCAAAAAGATACTGGGCCTCATGGGCCTTCCGCTCACTGCCCGCTTT
CCAGTCGGGAAACCTGTCGTGCCAGCTGCATTAACATGGTCATAGCTGTTTCCTTGCGTATTGGGCGCTCTCCGCTTCCTCGCTCACTGA
CTCGCTGCGCTCGGTCGTTCGGGTAAAGCCTGGGGTGCCTAATGAGCAAAAGGCCAGCAAAAGGCCAGGAACCGTAAAAAGGCCGCGTTG
CTGGCGTTTTTCCATAGGCTCCGCCCCCCTGACGAGCATCACAAAAATCGACGCTCAAGTCAGAGGTGGCGAAACCCGACAGGACTATAA
AGATACCAGGCGTTTCCCCCTGGAAGCTCCCTCGTGCGCTCTCCTGTTCCGACCCTGCCGCTTACCGGATACCTGTCCGCCTTTCTCCCT
TCGGGAAGCGTGGCGCTTTCTCATAGCTCACGCTGTAGGTATCTCAGTTCGGTGTAGGTCGTTCGCTCCAAGCTGGGCTGTGTGCACGAA
CCCCCCGTTCAGCCCGACCGCTGCGCCTTATCCGGTAACTATCGTCTTGAGTCCAACCCGGTAAGACACGACTTATCGCCACTGGCAGCA
GCCACTGGTAACAGGATTAGCAGAGCGAGGTATGTAGGCGGTGCTACAGAGTTCTTGAAGTGGTGGCCTAACTACGGCTACACTAGAAGA
ACAGTATTTGGTATCTGCGCTCTGCTGAAGCCAGTTACCTTCGGAAAAAGAGTTGGTAGCTCTTGATCCGGCAAACAAACCACCGCTGGT
AGCGGTGGTTTTTTTGTTTGCAAGCAGCAGATTACGCGCAGAAAAAAAGGATCTCAAGAAGATCCTTTGATCTTTTCTACGGGGTCTGAC
GCTCAGTGGAACGAAAACTCACGTTAAGGGATTTTGGTCATGAGATTATCAAAAAGGATCTTCACCTAGATCCTTTTAAATTAAAAATGA
AGTTTTAAATCAATCTAAAGTATATATGAGTAAACTTGGTCTGACAGTTACCAATGCTTAATCAGTGAGGCACCTATCTCAGCGATCTGT
CTATTTCGTTCATCCATAGTTGCCTGACTCCCCGTCGTGTAGATAACTACGATACGGGAGGGCTTACCATCTGGCCCCAGTGCTGCAATG
ATACCGCGAGAACCACGCTCACCGGCTCCAGATTTATCAGCAATAAACCAGCCAGCCGGAAGGGCCGAGCGCAGAAGTGGTCCTGCAACT
TTATCCGCCTCCATCCAGTCTATTAATTGTTGCCGGGAAGCTAGAGTAAGTAGTTCGCCAGTTAATAGTTTGCGCAACGTTGTTGCCATT
GCTACAGGCATCGTGGTGTCACGCTCGTCGTTTGGTATGGCTTCATTCAGCTCCGGTTCCCAACGATCAAGGCGAGTTACATGATCCCCC
ATGTTGTGCAAAAAAGCGGTTAGCTCCTTCGGTCCTCCGATCGTTGTCAGAAGTAAGTTGGCCGCAGTGTTATCACTCATGGTTATGGCA
GCACTGCATAATTCTCTTACTGTCATGCCATCCGTAAGATGCTTTTCTGTGACTGGTGAGTACTCAACCAAGTCATTCTGAGAATAGTGT
ATGCGGCGACCGAGTTGCTCTTGCCCGGCGTCAATACGGGATAATACCGCGCCACATAGCAGAACTTTAAAAGTGCTCATCATTGGAAAA
CGTTCTTCGGGGCGAAAACTCTCAAGGATCTTACCGCTGTTGAGATCCAGTTCGATGTAACCCACTCGTGCACCCAACTGATCTTCAGCA
TCTTTTACTTTCACCAGCGTTTCTGGGTGAGCAAAAACAGGAAGGCAAAATGCCGCAAAAAAGGGAATAAGGGCGACACGGAAATGTTGA
ATACTCATACTCTTCCTTTTTCAATATTATTGAAGCATTTATCAGGGTTATTGTCTCATGAGCGGATACATATTTGAATGTATTTAGAAA
AATAAACAAATAGGGGTTCCGCGCACATTTCCCCGAAAAGTGCCAC

**Text S2.** Detailed protocol.

### Selection-free CRISPR-Cas9 editing protocol for distant *Dictyostelid* species

**Table of contents**

| | Page |
|---|---|
| **Graphic protocol** | 24 |
| **List of materials** | 25 |
| **Buffers and reactions** | 27 |
| **Part 1. CRISPR-Cas9 editing protocol with RNP complex in *D. discoideum*** | 31 |
| **Step 1**. Identifying good CRISPR-Cas9 targets | 31 |
| **Step 2**. Designing donor oligos and donor PCR products | 32 |
| **Step 3**. Culturing axenic and non-axenic strains of *D. discoideum* | 36 |
| **Step 4**. Transfecting cells with RNP complex by electroporation | 37 |
| **Step 5**. Flow cytometry analysis | 40 |
| **Step 6**. Isolation of gene-edited cells using cell sorting | 41 |
| **Step 7**. Propagation of positive clones | 43 |
| **Part 2. Notes for CRISPR-Cas9 editing in other *Dictyostelid* species** | 44 |

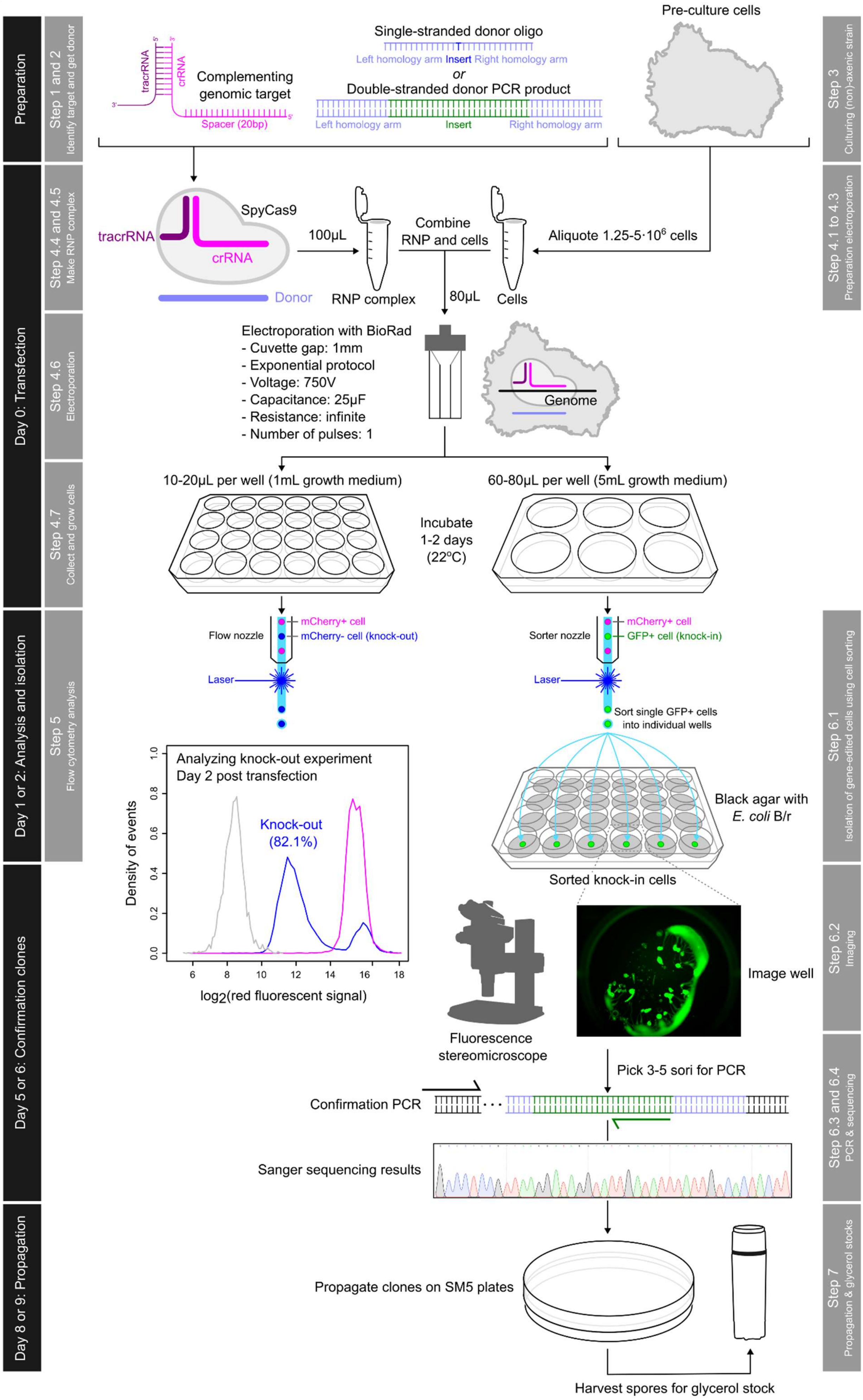

Preparation
Step 1 and 2
Identify target and get donor
tracrRNA
crRNA
Complementing genomic target
Spacer (20bp)
Single-stranded donor oligo
Left homology arm Insert Right homology arm
or
Double-stranded donor PCR product
Left homology arm
Insert
Right homology arm
Pre-culture cells
Step 3
Culturing (non)-axenic strain
Day 0: Transfection
Step 4.4 and 4.5
Make RNP complex
SpyCas9
tracrRNA
crRNA
Donor
100µL
RNP complex
Combine RNP and cells
80µL
Cells
Aliquote 1.25-5·10^6 cells
Step 4.1 to 4.3
Preparation electroporation
Step 4.6
Electroporation
Electroporation with BioRad
- Cuvette gap: 1mm
- Exponential protocol
- Voltage: 750V
- Capacitance: 25µF
- Resistance: infinite
- Number of pulses: 1
Genome
Step 4.7
Collect and grow cells
10-20µL per well (1mL growth medium)
60-80µL per well (5mL growth medium)
Incubate 1-2 days (22°C)
Day 1 or 2: Analysis and isolation
Step 5
Flow cytometry analysis
Flow nozzle
mCherry+ cell
mCherry- cell (knock-out)
Laser
Sorter nozzle
mCherry+ cell
GFP+ cell (knock-in)
Laser
Sort single GFP+ cells into individual wells
Step 6.1
Isolation of gene-edited cells using cell sorting
Analyzing knock-out experiment
Day 2 post transfection
Knock-out (82.1%)
Density of events
log2(red fluorescent signal)
Black agar with E. coli B/r
Sorted knock-in cells
Day 5 or 6: Confirmation clones
Fluorescence stereomicroscope
Image well
Step 6.2
Imaging
Pick 3-5 sori for PCR
Confirmation PCR
Sanger sequencing results
Step 6.3 and 6.4
PCR & sequencing
Day 8 or 9: Propagation
Propagate clones on SM5 plates
Harvest spores for glycerol stock
Step 7
Propagation & glycerol stocks

List of materials

| Material | Provider | Catalogue number/Reference |
|---|---|---|
| **Software** | | |
| Snapgene | | https://www.snapgene.com |
| Cas designer | | http://www.rgenome.net/cas-designer/ |
| **Chemicals and media** | | |
| Agarose | Sigma-Aldrich | A9539-500G |
| Acetic acid (glacial) | Sigma-Aldrich | 1000631000 |
| Activated charcoal | Sigma-Aldrich | C9157-500G |
| Bacto Dehydrated Agar | BD Biosciences | 214010 |
| Calcium Chloride ($CaCl_2$) | Sigma-Aldrich | C3881-500 |
| DRAQ7 | Thermo Fisher | D15106 |
| Ethanol | Sigma-Aldrich | 100.983 |
| Folic acid | Sigma-Aldrich | F8758-25G |
| Glycerol | Sigma-Aldrich | 1040572511 |
| HEPES, ultra-pure | Biomol | 5288.1 |
| HL5 Medium including Glucose | Formedium | HLG0102 |
| IGEPAL CA-630 | Sigma-Aldrich | I8896-50M |
| Potassium phosphate dibasic ($K_2HPO_4$) | Sigma-Aldrich | P8281-500G |
| Potassium chloride (KCl) | Sigma-Aldrich | 60130-1KG |
| Potassium phosphate monobasic ($KH_2PO4$) | Sigma-Aldrich | P0662-500G |
| Magnesium chloride hexahydrate ($MgCl_2·6H2O$) | Sigma-Aldrich | M9272-1kg |
| Magnesium sulfate ($MgSO_4$) | Sigma-Aldrich | M2643-500G |
| Sodium chloride (NaCl) | Sigma-Aldrich | 71380-1KG-M |
| Sodium phosphate monobasic ($NaH_2PO_4$) | Sigma-Aldrich | S3139-500G |
| Sodium bicarbonate ($NaHCO_3$) | Sigma-Aldrich | S5761-1KG |
| Penicillin G potassium salt (Benzylpenicillin potassium salt) | Sigma-Aldrich | P7794-10MU |
| SM Broth/5 | Formedium | SMB50102 |
| Streptomycin sulfate | Sigma-Aldrich | S9137-100G |
| Titriplex III (EDTA disodium salt) | Sigma-Aldrich | 1084180100 |
| Trizma base | Sigma-Aldrich | T1503-1KG |
| Vitamin B12 (Cyanocobalamin) | Sigma-Aldrich | V6629-250MG |
| Nuclease-free water ($H_2O$), for molecular biology (DEPC-free) | Sigma-Aldrich | W4502-10X50ML |
| Bacto Tryptone | Thermo Fisher | 211705 |
| Bacto Yeast Extract | Thermo Fisher | 212750 |
| **CRISPR reagents** | | |
| Alt-R CRISPR-Cas9 tracrRNA, 20 nmol | IDT | 1072533 |
| Alt-R CRISPR-Cas9 crRNA | IDT | Custom |
| Alt-R S.p. Cas9 Nuclease V3, 100 µg | IDT | 1081058 |
| Alt-R Cas9 Electroporation Enhancer, 10 nmol | IDT | 1075915 |
| Alt-R HDR Enhancer V2, 30 µL | IDT | 10007910 |
| pHO4d-Cas9 (for SpyCas9 purification) | Addgene | 67881 |
| TrueCut Cas9 Protein v2 | Thermo Fisher | A36498 |
| **Enzymes and kits** | | |
| MinElute PCR Purification Kit | Qiagen | 28006 |
| QIAquick PCR Purification Kit | Qiagen | 28106 |

| | | |
|---|---|---|
| Proteinase K | NEB | P8107S |
| Q5 Hot Start High-Fidelity 2X Master Mix | NEB | M0494L |
| Taq 2X Master Mix - 500 reactions | NEB | M0270L |
| **Plasticware/disposable material** | | |
| Disposable loops 10µL | Thermo Fisher Scientific | 10048750 |
| Disposable loops 1µL | Thermo Fisher Scientific | 10344741 |
| Spreaders, T-shaped | VWR | 612-2653 |
| Nunc multidish 24-wells | Thermo Fisher Scientific | 10604903 |
| Nunc multidish 6-wells | Thermo Fisher Scientific | 10119831 |
| Petri dishes, steriplan, Duran 100x20 | Duran | 391-2840 |
| Gene Pulser/MicroPulser electroporation cuvettes, 0.1cm gap | Bio-Rad Laboratories | 1652089 |
| Countess cell counting chamber slides | Thermo Fisher Scientific | C10312 |
| Falcon 5 mL round bottom polystyrene test tube, without cap | Falcon | 352008 |
| Falcon round-bottom polystyrene test tubes with cell strainer snap cap, 5mL | Falcon | 352235 |
| **Equipment** | | |
| BD FACSAria Fusion Flow Cytometer | BD Biosciences | |
| Gene Pulser Xcell Microbial System | Bio-Rad Laboratories | 1652662 |
| BD FACSymphony A3 Analyzer | BD Biosciences | |
| Axio Zoom.V16 Stereo Zoom Microscope | Zeiss | |
| Countess 3 starter package 1 | Thermo Fisher Scientific | A49865 |
| NanoDrop 8000 Spectrophotometer | Thermo Fisher Scientific | ND-8000-GL |
| BioPhotometer plus | Eppendorf AG | |
| C1000 Touch Thermal Cycler | Bio-Rad Laboratories | 1851197 |

#### Buffers and reactions

| Reagents/ solution | Final concentration | Stock reagent/ solution | Amount/ volume |
|---|---|---|---|
| 10x KK2 | 161.6mM $KH_2PO_4$ | $KH_2PO_4$ Powder | 22g |
| | 40mM $K_2HPO_4$ | $K_2HPO_4$ Powder | 7g |
| **Instructions.** Dissolve in 1L of deionized $H_2O$ and autoclave. Store at RT. | | | |
| 1x KK2-MC | 1xKK2 | 10xKK2, autoclaved | 100mL |
| | 50µM $MgCl_2$ | 1 M $MgCl_2$, autoclaved | 50µL |
| | 50µM $CaCl_2$ | 1 M $CaCl_2$, autoclaved | 50µL |
| **Instructions.** Mix autoclaved components, adjust volume to 1L with deionized $H_2O$, filter sterilized. Store at RT. | | | |
| SM5-agar | 1x SM Broth/5 | SM Broth/5 powder | 7.9g |
| | 1.6% BD Bacto Dehydrated Agar | BD Bacto Dehydrated Agar powder | 16g |
| **Instructions.** Mix reagents while stirring, adjust volume to 1L with deionized $H_2O$, autoclave at 110°C. Use 25mL of SM5-agar for each 10cm Petri dish. Store at 4°C. Dry plates in a laminar hood for 15 min before use. | | | |
| SM5-charcoal-agar | 1x SM Broth/5 | SM Broth/5 powder | 7.9g |
| | 1.6% BD Bacto Dehydrated Agar | BD Bacto Dehydrated Agar powder | 16g |
| | 0.5% Activated charcoal | Activated charcoal | 5g |
| **Instructions.** Mix reagents while stirring, adjust volume to 1L with deionized $H_2O$, autoclave at 110°C. Use 1mL of SM5-agar-charcoal per well for 24-well plates and 5mL per well for 6-well plates. Store at 4°C. Dry plates in a laminar hood for 15 min before use. | | | |
| L-Broth | 1% Bacto Tryptone | Bacto Tryptone powder | 10g |
| | 0.5% Bacto Yeast Extract | Bacto Yeast Extract | 5g |
| | 0.5% NaCl | NaCl Powder | 5g |
| **Instructions.** Mix all reagents while stirring, adjust volume to 1L with deionized $H_2O$, and autoclave at 110°C. Store at RT. | | | |
| L-Agar | 1% Bacto Tryptone | Bacto Tryptone Dehydrated | 10g |
| | 0.5% Bacto Yeast Extract | Bacto Yeast Extract Dehydrated | 5g |
| | 1% NaCl | NaCl Powder | 10g |
| | 1.5% BD Bacto Dehydrated Agar | BD Bacto Dehydrated Agar, powder | 15g |
| **Instructions.** Mix all reagents while stirring, adjust volume to 1L with deionized $H_2O$, and autoclave at 110°C. Use 25mL L-Agar for each 10cm petri dish. Store at 4°C. Dry plates in a laminar hood for 15 min before use. | | | |
| H50 | 50mM KCl | 1M KCl, autoclaved | 50mL |
| | 20mM HEPES pH 7.0 | 400mM HEPES pH7, filter sterilized | 50mL |
| | 10mM NaCl | 1M NaCl, autoclaved | 10mL |
| | 5mM $NaHCO_3$ | 500mM $NaHCO_3$, filter sterilized | 10mL |
| | 1mM $NaH_2PO_4$ | 100mM $NaH_2PO_4$, autoclaved | 100mL |
| | 1 mM $MgSO_4$ | 1M $MgSO_4$, autoclaved | 1mL |
| **Instructions.** Mix components, adjust volume to 1L with deionized $H_2O$ and filter sterilize. Store at 4°C. | | | |
| Quick Genomic DNA extraction buffer | 0.5x Taq buffer | 10x Taq buffer | 1µL |
| | 0.5% IGEPAL CA-630 | IGEPAL CA-630 liquid | 0.1µL |
| | 50ng/µL Proteinase K | 20mg/mL Proteinase K, solution | 0.05µL |
| | $H_2O$ | Nuclease-Free $H_2O$ | 18.85µL |

**Instructions.** Mix all components on ice, and use 20µL per extraction. Prepare freshly before use.

| | | | |
|---|---|---|---|
| Quick Genomic DNA extract | Quick Genomic DNA extraction buffer | Quick Genomic DNA extraction buffer | 20µL |
| | 3-5 sorus of monoclonal species | Sorus of Fruiting bodies growing in SM5-agar plates | 3-5 sorus |

**Instructions.** Prepare Quick genomic DNA extraction buffer freshly. Make 20µL aliquots in PCR tubes. With a 10µL tip, pick 3-5 sorus and dip them in the extraction buffer. Incubate at 56°C for 45 min and 95°C for 10 min. Store at 4°C for up to 3 days.

| | | | |
|---|---|---|---|
| Clone confirmation PCR reaction | 1x Taq polymerase | Taq 2X Master Mix | 10µL |
| | 0,2µM Forward primer | 10µM Forward primers, in 10mM Tris-Cl, pH 8.5 | 0.4µL |
| | 0.2µM Reverse primer | 10µM Reverse primers, in 10mM Tris-Cl, pH 8.5 | 0.4µL |
| | $H_2O$ | Nuclease-free $H_2O$ | To 20µL |
| | Quick Genomic DNA extract | Quick Genomic DNA extract | 2µL |

**Instructions.** Set up PCR reaction on ice by mixing all components. A master mix can be prepared for several samples, and 18µL aliquots prepared in PCR tubes. Two microliter of Quick Genomic DNA extract are added to each reaction. PCR cycle: 1x initial denaturalization cycle at 95°C for 2 min, 35x amplification cycles at 95°C for 30 sec, $T_a$ depending on primers for 20 sec, 68°C for 1 min, and 1x final extension cycle at 68°C for 10 min. Run 2µL PCR product, with 2µL 6x loading dye and 6µL nuclease-free $H_2O$, in a 1% agarose gel. Purify the remaining 18µL with a Qiagen PCR purification column, elute with 25µL EB and send for Sanger sequencing.

| | | | |
|---|---|---|---|
| 1000x Pen/Strep for HL5 | 100mg/mL Streptomycin sulfate | Streptomycin sulfate, powder | 1g |
| | 70mg/mL Penicillin G potassium salt | Penicillin G potassium salt (Benzylpenicillin potassium salt), powder | 0.7g |

**Instructions.** Mix all components and adjust to 10mL with deionized $H_2O$. Filter sterilize with 0.22µm syringe filters. Make 1mL aliquots and store at -20°C. Add to HL5 just before use to a final concentration of 1x.

| | | | |
|---|---|---|---|
| 10000x VitB12 / Folic acid | 2mg/mL Folic acid | Folic acid, powder | 20mg |
| | 6mg/mL Vitamin B12 | Vitamin B12 (Cyanocobalamin), powder | 60mg |

**Instructions.** Mix all components and adjust to 10mL with deionized $H_2O$. Adjust pH to 7.0 with NaOH. Filter sterilize with 0.22µm syringe filters. Make 0.1mL aliquots and store at -20°C. Add to HL5 just before use to a final concentration of 1x.

| | | | |
|---|---|---|---|
| HL5 broth | 35.5g/L HL5 | HL5 Medium including Glucose, powder | 35.5g |

**Instructions.** Mix with deionized $H_2O$ while stirring, adjust volume to 1L, and autoclave at 110°C. Store at 4°C.

| | | | |
|---|---|---|---|
| HL5+FAB | HL5 broth | HL5 broth, autoclaved | Required volume for the experiment |
| | 1x Pen/Strep for HL5 | 1000x Pen/Strep for HL5, solution | 1:1000 (v:v) |
| | 1x VitB12/Folic acid | 10000x VitB12/Folic acid, solution | 1:10000 (v:v) |

**Instructions.** Warm the needed volume of 1x HL5 Broth to 22°C and add Pen/Strep and VitB12/Folic acid to a final concentration of 1x just before use.

| | | | |
|---|---|---|---|
| RNP complex | 12µM Cas9 | ~15µg/µL Cas9 in 20mM HEPES pH 7.5, 500mM KCl, 10% glycerol. Produced by EMBL PeP-core facility[9] | 1.31µL[10] |

[9] The best results are obtained with ultrapure WT SpyCas9 (Figure S4B). However, good efficiencies are also obtained using a quick 1-day purification protocol. TrueCut Cas9 Protein v2 (Thermo Fisher, A36498) is the best-performing commercially available SpyCas9 we tested (Figure S4B).

[10] Volume might change depending on batch (depending on SpyCas9 activity).

| | | | |
|---|---|---|---|
| | 15µM gRNA complex | 50 µM gRNA complex (100µM Alt-R CRISPR-Cas9 tracrRNA and 100µM Alt-R CRISPR-Cas9 crRNA in nuclease-free $H_2O$, pre-annealed) | 3µL[11] |
| | $H_2O$ | Nuclease-free water | Up to 10µL |
| **Instructions.** For each transfection, mix 1.5µL of 100µM Alt-R CRISPR-Cas9 tracrRNA and 1.5µL of 100µM Alt-R CRISPR-Cas9 crRNA in a PCR tube, and anneal by heating at 95ºC for 5 min, and slowly cooling down (-1ºC/cycle, 20sec/cycle). Scale up depending on the number of transfections with the same crRNA. The 50µM gRNA complex can be stored at -20ºC for up to 1 year. Mix components in a 1.5mL tube at RT following the indicated order, by slowly adding each component while swirling the tip, and pipetting once up and down after the addition of each component. | | | |
| Transfection mix | RNP complex (see above) | 12µM Cas9 / 22µM Alt-R CRISPR-Cas9 tracrRNA / 22µM Alt-R CRISPR-Cas9 crRNA | 10µL |
| | Donor oligo, as indicated | 100µM oligo, in nuclease-free $H_2O$ | As indicated |
| | Donor PCR product, as indicated | 1µg/µL donor PCR product | As indicated |
| | EE | 100µM Alt-R Cas9 Electroporation Enhancer (EE), in nuclease-free $H_2O$ | Optional[12] |
| | H50 | H50 buffer at RT | Up to 100µL in small volume increments |
| | $1.25\text{-}5\cdot10^6$ Dictyostelid cells | Cells washed in ice-cold H50 | Cell pellet |
| **Instructions.** To 10µL of RNP complex, add donor oligos, donor PCR products and/or EE when indicated. Bring up volume to 100µL by adding H50 buffer in small increments (for example 25+25+40µL). Spin down aliquots of cells, remove supernatant, and resuspend pellet with 100µL RNP complex in H50 buffer. Transfer 80µL to a pre-chilled electroporation cuvette. After electroporation, transfer cells to growth media as indicated. | | | |
| Donor PCR product | 1x Q5 Hot Start High-Fidelity | Q5 Hot Start High-Fidelity 2X Master Mix | 200µL |
| | 0.5µM Forward Primer | 10µM Forward Primer in 10mM Tris-Cl, pH 8.5 | 20µL |
| | 0.5µM Reverse Primer | 10µM Reverse Primer in 10mM Tris-Cl, pH 8.5 | 20µL |
| | 0.2ng/µL plasmid DNA Template | 5ng/µL plasmid DNA Template in 10mM Tris-Cl, pH 8.5 | 16µL |
| | $H_2O$ | Nuclease-Free $H_2O$ | Up to 400µL |
| **Instructions.** For a transfection with 10µg of donor PCR product, prepare 400µL of donor PCR reaction by mixing all the components on ice. Split in 8x 50µL in PCR tubes, and use the following PCR cycle: 1x Initial Denaturalization cycle at 98ºC for 30s, 30x Amplification cycles at 98ºC for 10 sec, 61.5ºC for 10 sec, 72ºC for 30 sec, and 1x Final Extension cycle at 72ºC for 4 min. Pool together 400µL of PCR reaction and purify using one Qiagen MiniElute column. Elute with 12µL nuclease-free $H_2O$. Quantify using a NanoDrop 8000 Spectrophotometer. $T_a$ might have to be optimized for different templates/primers. | | | |
| DRAQ7 | 3µM DRAQ7 | DRAQ7 Dye (0.3mM in aqueous buffer) | 1:100 (v:v) |
| **Instructions.** Add 1:100 (v:v) to cells in KK2-MC buffer and briefly vortex. Incubate 5 min before analyzing cells with flow cytometer. | | | |
| 50xTAE | 0.5M Trizma base | Trizma base, powder | 242g |
| | 0.05M EDTA, pH 8.0 | 0.5M EDTA, pH 8.0, autoclaved | 100mL |
| | 57% Acetic acid | Acetic acid (glacial), 100% anhydrous | 57mL |
| **Instructions.** Mix all components and adjust to 1L with deionized $H_2O$. | | | |
| 1xTAE | 1xTAE | 50xTAE | 20mL |

[11] Optional: pre-annealing of tracrRNA and crRNA to generate a gRNA complex can be omitted, resulting in slightly lower efficiencies (Figure S4B).
[12] Comes with the risk of inserting EE derived DNA sequences into your target site.

| | | | |
|---|---|---|---|
| **Instructions.** Add 20mL 50xTAE to 980mL deionized $H_2O$. Store at RT. | | | |
| 1% agarose gel | 1% agarose | Agarose, powder | 1g |
| | 1xTAE | 1xTAE | 100mL |
| **Instructions.** Add agarose to 100mL 1xTAE while stirring. Microwave for 1 min, stir, microwave for 1 min. Let it cool down for 5 min. Poor on a tray, add 3μL SYBR Safe, gently mix evenly, place a comb, and let the gel polymerase for 20 min. After loading, run the gel for 30-40 min at 100-120V. | | | |

#### Part 1. CRISPR-Cas9 editing protocol with RNP complex in *D. discoideum*.

##### Step 1. Identifying good CRISPR-Cas9 targets.

*The RNP complex consists of the endonuclease Cas9, crRNA and tracrRNA. The 20nt spacer sequence of crRNA complements the genomic target site. The target site also needs an adjacent PAM sequence to be recognized by Cas9. We use Streptococcus pyogenes Cas9 (SpyCas9) that recognizes the PAM sequence 'NGG'. Before we can start gene editing, we should first identify good CRISPR-Cas9 target sites in the genome, which allows us to design and order a custom crRNA. The other components of the RNP complex are invariable (SpyCas9 and tracrRNA).*

*Tools to identify CRISPR-Cas9 targets.*

- *There are several tools available for identifying potential CRISPR targets. By providing both the species of interest and gene of interest (GOI), these tools will identify unique CRISPR targets within the GOI that have no or minimal off-target hits elsewhere in the genome. Popular tools are for example Cas-designer (http://www.rgenome.net/cas-designer/) (Bae et al. 2014a, 2014b; Park et al. 2015) and CRISPOR (http://crispor.tefor.net/) (Concordet and Haeussler 2018). The reference genome of D. discoideum is available for both these tools.*

*Tools to simulate CRISPR-Cas9 edit.*

- *Before starting gene editing, we recommend simulating the CRISPR-Cas9 edit with a cloning software, such as SnapGene or Benchling, to ensure the correct crRNA and donor template design for CRISPR-Cas9 editing. In this protocol, we use SnapGene for that purpose (https://www.snapgene.com/).*

*For introducing knock-out or knock-in mutations, it is important to use the genomic sequence associated with the GOI for designing the crRNA, while making sure that the insertion site is present in an exonic region (i.e., present in the coding DNA sequence). In addition, the cutting site should be as close as possible to the desired insertion site. We recommend a window of +/-10bp from the cutting site (Paix et al. 2019). In the current study, we always cut within +/-3bp from the desired insertion site.*

1.1. Identifying a CRISPR-Cas9 target for a GOI.

- Find both the genomic sequence and the coding DNA sequence (CDS) associated with a GOI on dictyBase (http://dictybase.org/).
- Open Cas designer and fill out the below information.
  - PAM type: SpyCas9 from *Streptococcus pyogenes*: 5'-NGG-3'
  - Target genome: Others, *Dictyostelium discoideum* (dictyBase)
  - Target sequence: Genome sequence containing the CDS[13] of a GOI.
  - crRNA length: 20.
- Identify spacer sequences that are associated with best target sites, ideally:
  - No off-targets hits for spacer sequences with 0, 1 or 2 mismatches.
  - Out-of-frame score of ≥66.
  - GC content >20% (without the PAM sequence).
  - No AT rich or repetitive sequences[14].

[13] Cas-designer only allows up to 1000bp sequences. Longer genes can be split into smaller fragments. Targeting the 5' end of the CDS works best for creating complete knock-outs. Targeting the 3' end might result in only partial deletion of the protein.

[14] Cas designer tool returns 30 possible hits per search. Since *Dictyostelid* genomes are AT rich and repetitive, when there are several spacer sequences possible, choosing a spacer associated with a less repetitive sequence can reduce off-target effects and simplify downstream PCRs to confirm gene edits.

- Choose two or three spacer sequences targeting a GOI[15]. Below we show three spacer sequences we identified for *mCherry* in *D. discoideum* AX2 *act5::mCherry* (visualized in SnapGene).

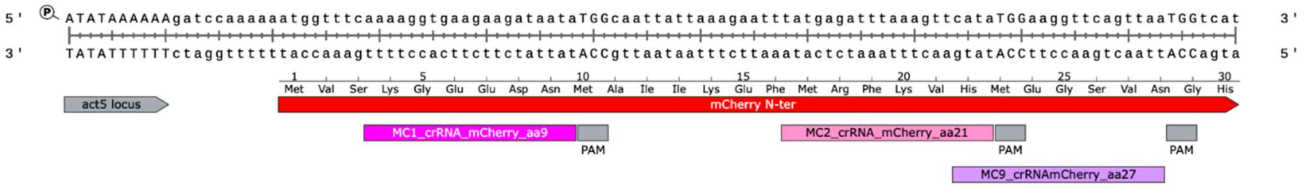

1.2. Order and prepare crRNA.

- When ordering from IDT, enter the 20nt spacer sequence upstream from the PAM. The ordering tool will automatically convert the DNA sequence into an RNA sequence (see also Figure 1A). In the case of MC1 targeting *mCherry*, the spacer sequence for the crRNA is:

```
Target sequence   5'-...GGTTTCAAAAGGTGAAGAAGATAATATGGCAATTA...-3'
Ordered crRNA               5'-aaaaggtgaagaagataata-3'
```

- 2nmol crRNA is sufficient for 8 or 9 experiments.
- Resuspend crRNA in nuclease-free $H_2O$ to a final concentration of 100µM.

1.3. Order and prepare tracrRNA.

- As the same tracrRNA can be used for all CRISPR-Cas9 experiments, we recommend ordering a larger quantity: 20nmol of tracrRNA is sufficient for 80-90 experiments.
- Resuspend tracrRNA in nuclease-free $H_2O$ to a final concentration of 100µM.

#### Step 2. Designing donor oligos and donor PCR products.

*To introduce precise genome edits using HDR, a donor oligo or donor PCR product that serves as a repair template is needed. Similar to designing crRNAs, when designing a repair template, it is important to use the genomic sequence of the target site, and not the CDS, since target sites can be close to exon/intron junctions. Genome editing is also possible without repair template, but will result in less-efficient NHEJ-mediated indels. Knock-out of a GOI can be efficiently produced with donor oligos with 1 or 4bp insertions that result in a frameshift. Short sequences, such as FLAG-tags, HA-tags or myc-tags can also be introduced using single-stranded donor oligos. For knock-outs, we find that donor oligos with a 37bp insertion (coding for HA-tag, a stop codon and a frameshift mutation) and 28bp homology arms give the best efficiencies (Figure 1). To introduce larger inserts, to for example generate knock-ins, PCR products can be used as repair templates for HDR. The donor PCR product consists of the insertion sequence (for example a the CDS of a fluorescent protein) and ~30bp homology arms (see Figure 2 and Paix et al. 2017). The best efficiencies are obtained for lower insertion sizes, ideally smaller than 1000bp. Similarly to HDR with donor oligos, for producing knock-ins by HDR with a donor PCR product, the insertion site should be as close as possible to the cutting site. It is also important that the CRISPR-Cas9 target site is removed after creating the knock-in to ensure single edits only. The target site can be removed by either removing the PAM sequence or removing complementarity to the crRNA spacer sequence.*

[15] Make sure the cutting sites of the selected spacers are on the CDS of a GOI (and not in an intronic region).

2.1. Design of donor oligos with 1bp insertion for producing knock-outs.

- Design the donor oligo with a 1bp insertion to introduce a frameshift mutation in the GOI with 28bp homology arms flanking the insertion site[16]. If possible, search for a 1bp insertion near the cutting site that will introduce a stop codon[17]. Below is the donor oligo we used for introducing a 1bp insertion in *mCherry* 1bp away from the cutting site, causing both a stop codon and frameshift mutation.

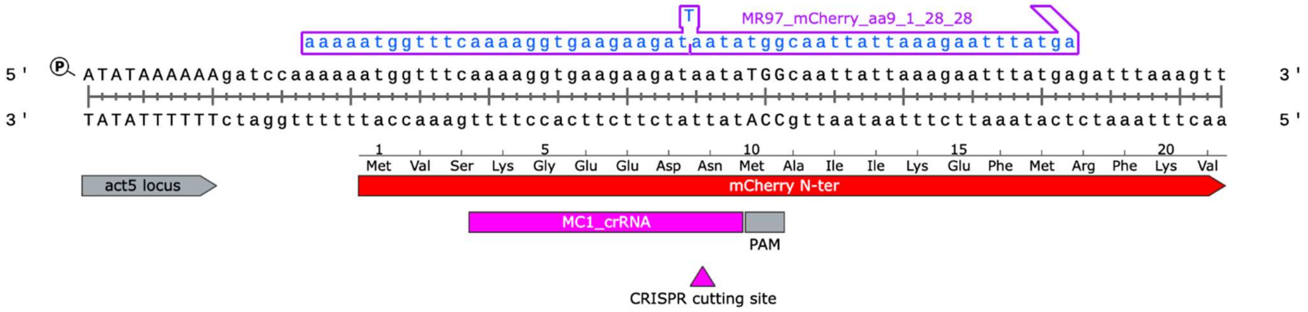

- Simulate the CRISPR-Cas9 edit using your crRNA and designed donor oligo in a cloning software (e.g., SnapGene). Below is the final product of the CRISPR-Cas9 edit with a 1bp insertion for our *mCherry* target. By inserting 1bp we introduce several stop codons in the amino acid sequence of *mCherry*.

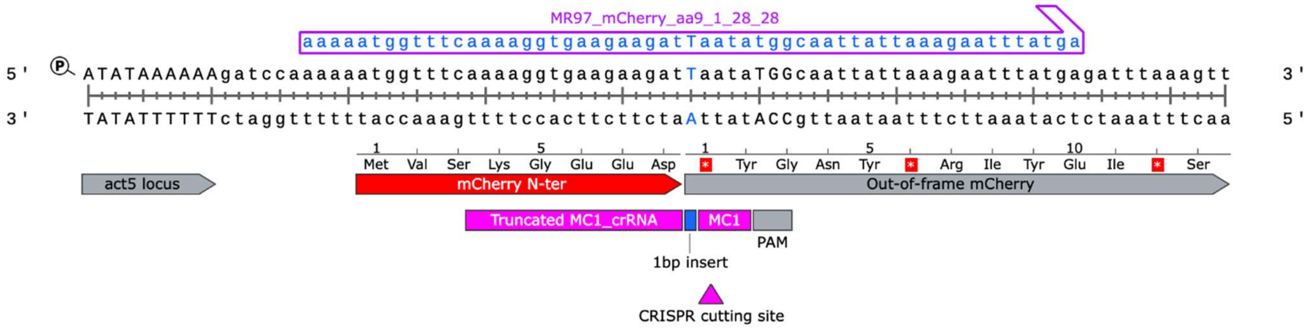

2.2. Design of donor oligos for introducing tags.

- Design a donor oligo with the sequence of the tag of interest and 28bp homology arms flanking the insertion site.
- To knock-out the target gene, add a stop codon and an additional 1bp to produce a frameshift mutation. In the example below we introduce an HA-tag, a stop codon and a frameshift mutation to knock-out *mCherry*.

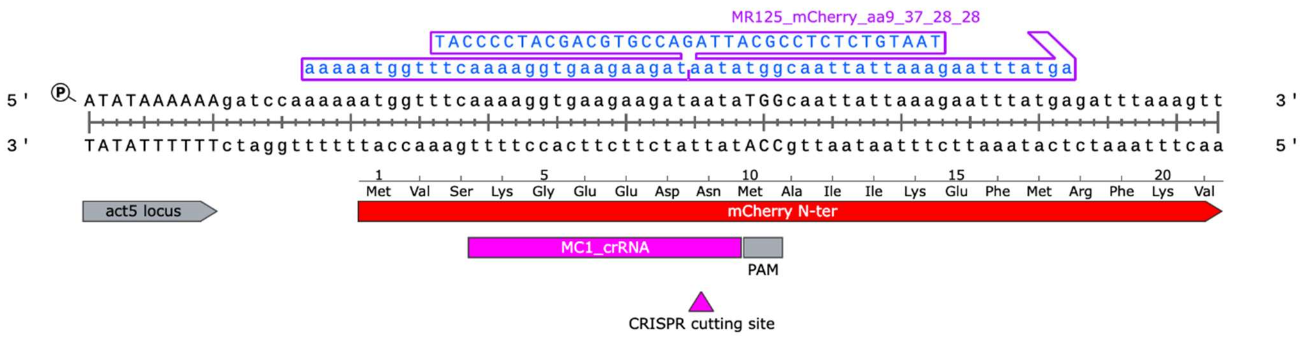

16 SpyCas9 cuts the genome between the 3rd and 4th bp upstream of the PAM. We recommend designing an insertion that falls within +/-10bp from the cutting site (Paix et al. 2019). In the example above, the insertion site is 1bp from the predicted cutting site.

17 Frameshift mutations will lead to several random stop codons downstream of the insertion thereby ensuring proper knock-out.

- Simulate the CRISPR-Cas9 edit in SnapGene to ensure that the tag sequence is in frame, and that stop codons have been introduced when knocking-out the target gene.

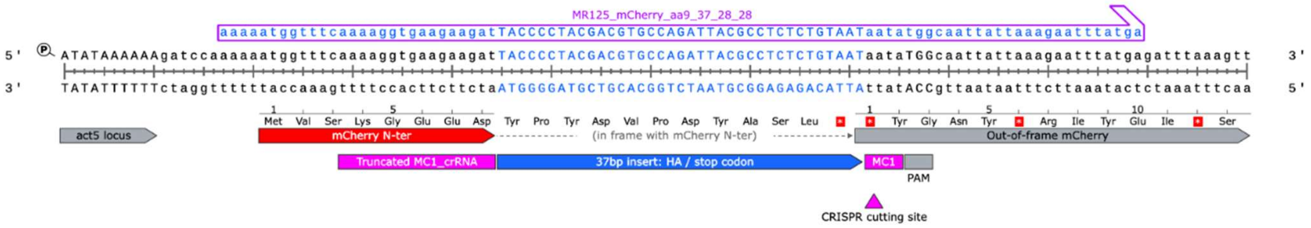

2.3. Design of donor PCR products for generating knock-in mutants.

- Design forward and reverse primers with 18-22 nucleotides of homology to the template DNA (e.g., pMG005) and 30-40bp overhangs with homology to the genomic sequences flanking the insertion site (i.e., homology arms). We recommend having a G or C at the 3'end of each primer. The $T_m$ of the primer pair should be similar (<3ºC difference). We usually target for a $T_m$ of 59ºC and adjust the length of the homology arms accordingly. In the example below, we create an in-frame knock-in mutation in *mCherry* using *mNeonGreen-P2A* as template (pMG005). The final product expresses both *mNeonGreen* and *mCherry* from the same promoter (Zhu et al. 2023). In our example, the forward/reverse primers were designed with 34/37bp homology arms (blue sequence) and 19/20bp annealing (black sequence) to the template (pMG005), respectively.

AP1169: 5'GATCCAAAAAATGGTTTCAAAAGGTGAAGAAGATGTATCTAAGGGAGAAGAAG 3'
AP1170: 5'CTTTAAATCTCATAAATTCTTTAATAATTGCCATATTAGAACTAGATCCACTAGGTC 3'

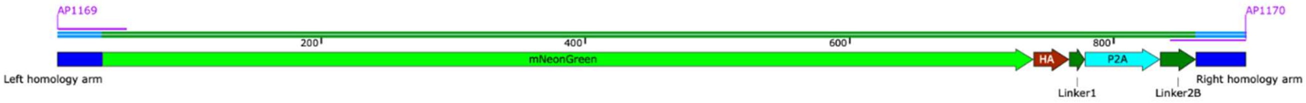

**AP1169/AP1170 donor PCR product**
899 bp

- Using a cloning software, simulate the PCR product and CRISPR-Cas9 edit to ensure the crRNA and donor PCR product are designed properly and no unintentional frameshift mutations or stop codons are introduced. For our example, the knock-in is expected to look as follows:

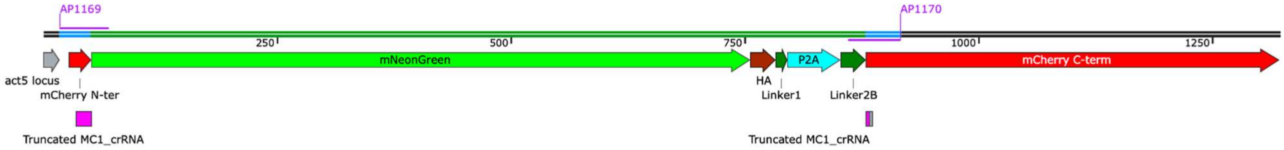

**act5::mNeonGreen-P2A-mCherry**

2.4. PCR reaction.

- Prepare ~10µg of PCR product[18] for each transfection, in 8x50µL PCR reactions and run the PCR using the following conditions:

| Template | Primer fwd | Primer rev | PCR product size (bp) |
|---|---|---|---|
| pMG005 | AP1169 | AP1170 | 899 |

[18] 10µg of donor DNA product gives good knock-in efficiencies (Figure 2), although lower amounts are effective as well (Figure 3).

| Reagent | Volumes for 8x50μL |
|---|---|
| Q5 Hot Start High-Fidelity 2X Master Mix | 200μL |
| 10μM forward primer | 20μL |
| 10μM reverse primer | 20μL |
| 5ng/μL template DNA | 16μL |
| Nuclease-free $H_2O$ | Up to 400μL (Split in 8 reactions) |

| Step | Temperature | Time |
|---|---|---|
| Initial Denaturation | 98°C | 30 sec |
| Amplification: 30 cycles | 98°C | 10 sec |
| | 61.5°C[19] | 10 sec |
| | 72°C | 30 sec |
| Final Extension | 72°C | 4 min |
| Hold | 4–10°C | |

- Run 2μL of PCR product from one of the tubes (with 2μL 6x loading dye and 6μL $H_2O$) in a 1% agarose gel to confirm the PCR was successful.
- In the meantime, pool together all PCR products (8x50μL) and purify them using a single Qiagen MiniElute PCR purification column. In brief:
    - Pool together 400μL PCR reaction in a 15mL tube.
    - Add 2000μL PB buffer to the tube.
    - Add 20μL 3M NaOAc to the tube.
    - Add 750μL PCR product/PB buffer to the MiniElute column.
    - Spin at 13000xg for 30 sec.
    - Discard flowthrough.
    - Repeat using the same column, until all the PCR product has been applied to the same column.
    - Add 750μL PE buffer.
    - Spin at 13000xg for 30 sec.
    - Discard flowthrough.
    - Spin at 13000xg for 2 min to eliminate all the ethanol.
    - Transfer the column to a clean 1.5mL tube.
    - Let the column stand for 2 min.
    - Add 12μL of nuclease-free $H_2O$ to the center of the column.
    - Let the column stand for 2 min.
    - Spin at 13000xg for 1 min.
    - Discard column and recover DNA.
    - Use 1μL of purified DNA to measure the concentration by NanoDrop.
    - We expect to have a concentration of at least 1μg/μL.
    - The PCR product can be stored at -20°C until further use.

[19] $T_a$ might have to be adjusted depending on the primers and the amplified sequence. We design all our primers to have a $T_m$ of ~59°C, and successfully use a $T_a$ of 61.5°C during PCR amplification.

#### Step 3. Culturing axenic and non-axenic strains of *D. discoideum.*

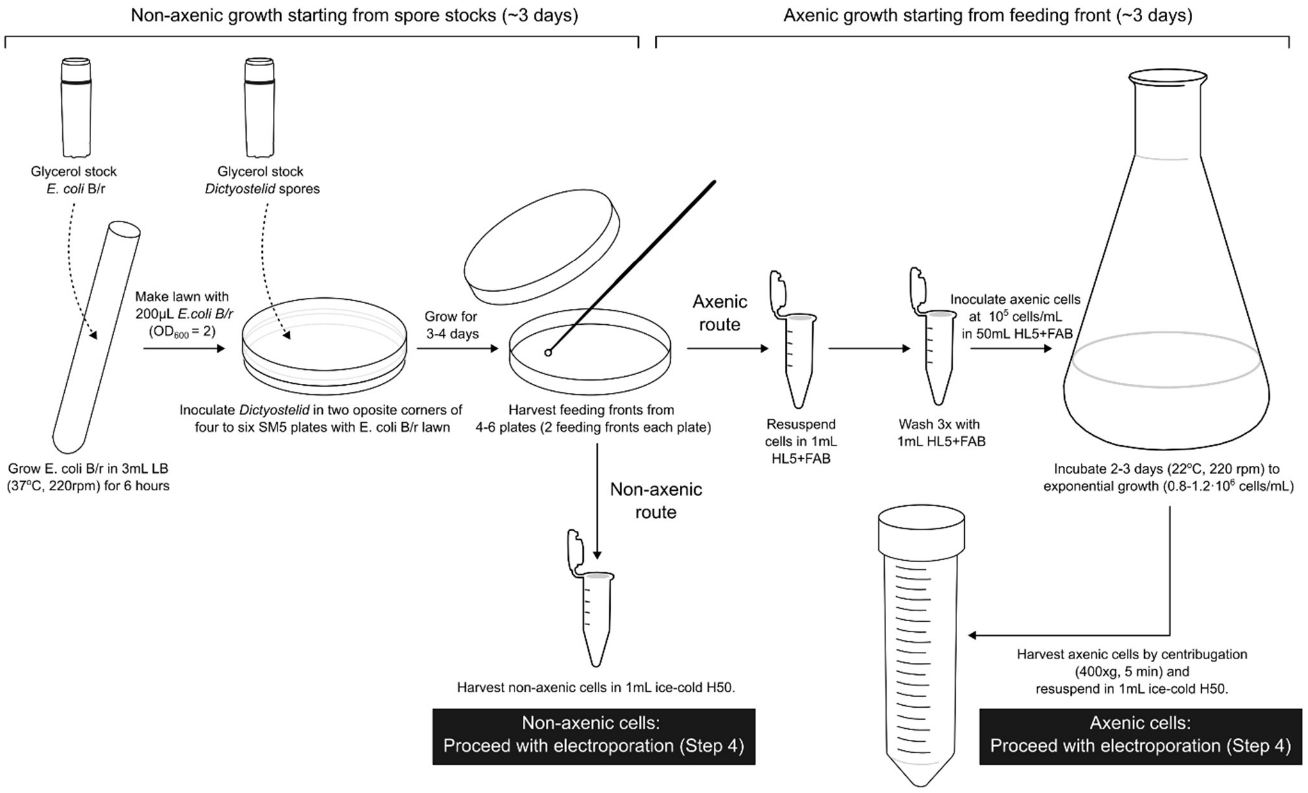

3.1. Non-axenic growth of *D. discoideum* on *E. coli* B/r.

- Prepare *E. coli* B/r.
  - Inoculate 50mL of LB with *E. coli* B/r from glycerol stock.
  - Shake O/N at 37°C 220 rpm[20].
  - Transfer culture to a 50mL falcon tube.
  - Pellet cells (4000xg, 5 min) and remove supernatant.
  - Wash pellet twice with 1mL KK2 buffer.
  - Resuspend pellet in 1mL KK2.
  - Measure culture density ($OD_{600}$).
  - Concentrated *E. coli* B/r in KK2 can be stored at 4°C for up to one week.
- Prepare four to six SM5-agar plates for each strain.
  - Dry plates in the hood with the lid off for 15 min.
  - Adjust density of *E. coli* B/r in KK2 from previous step to $OD_{600}$=2 with KK2 buffer.
  - Spread 200µL of *E. coli* B/r at $OD_{600}$=2 on the plates.
  - Distribute evenly with a spreader to form a bacteria lawn as food source.
  - Let it dry in the hood for 3-5 min.
- Inoculate *D. discoideum* strain directly from a glycerol stock[21].
  - Take glycerol stock from -80°C on dry ice.
  - With a loop or pipette, take some spores from the glycerol stock.
  - Inoculate two opposite corners of the plate.
- Incubate at 22°C for:
  - 2-3 days for harvesting both feeding fronts from the same plate.
  - 5-7 days for harvesting spores from a lawn of fruiting bodies.

---

[20] Alternatively, a 3mL culture can be grown at 37°C (220 rpm) for 6 hours on the same day.

[21] Our glycerol stocks are made from fruiting body spores.

3.2. Axenic growth of *D. discoideum* in HL5.

- Harvest *D. discoideum* cells from SM5-agar plates.
    - With a 10μL loop, harvest cells from the feeding fronts from 4 to 6 SM5-agar plates (2 feeding fronts from each plate).
    - Resuspend cells in 1mL of HL5+FAB in a 1.5mL tube.
    - Spin down cells (400xg, 2 min) and remove supernatant.
    - Wash cells twice with 1mL HL5+FAB.
    - Resuspend cells in 1mL HL5+FAB.
    - Dilute cells 1/10 in HL5+FAB.
    - Measure cell density using a hemocytometer or an automated cells counter (e.g., Countess III cell counter).
- Inoculate an HL5+FAB culture.
    - Prepare 50mL of HL5+FAB in a 250mL flask (sufficient for 5-10 transfections).
    - Add cells to have a final density of $10^5$ cells/mL.
    - Grow cells at 22°C (180rpm) for 2-3 days until a density of $0.8\text{-}1.2 \cdot 10^6$ cells/mL (exponential growth).

#### Step 4. Transfecting cells with RNP complex by electroporation.

4.1. Prepare non-axenic cells for electroporation.

- Prepare 4 to 6 SM5-agar plates with the cells of interest as indicated in Step 3.1.
- Incubate plates for 2-3 days or until there are two visible feeding fronts per plate.
- Harvest cells from the feeding fronts of 4 to 6 plates with a 10μL inoculation loop.
- Transfer them to a 1.5mL tube with 1mL of ice-cold H50 buffer.
- Spin cells down (400xg, 2 min) and remove supernatant
- Wash cells with 1mL of ice-cold H50 twice.
- Resuspend cells in 1mL of ice-cold H50.
- Make a 1/100 dilution and count cells.
- Make aliquots of $1.25\text{-}5 \cdot 10^6$ cells for each transfection[22]. The volume is not important, because cells will be pelleted again later. Keep on ice.
- Proceed with Step 4.3.

4.2. Prepare axenic cells for electroporation.

- Count cells from Step 3.2 daily until they reach exponential phase.
    - Transfer 1mL of culture with serological pipette to a 1.5mL tube.
    - Count cells as in previous step. If they are within the right interval of $0.8\text{-}1.2 \cdot 10^6$ cells/mL, proceed with the next step. If cells are overgrown (over $2 \cdot 10^6$ cells/mL), back dilute them to $5 \cdot 10^5$ cells/mL, and let them grow for another day. They should be ready on the following day.
- Prepare aliquots for transfection.
    - Transfer 45mL of cell suspension ($0.8\text{-}1.2 \cdot 10^6$ cells/mL) to a 50mL tube.
    - Spin cells down (400xg, 5 min) and remove supernatant. Be careful not to remove the pellet as it does not firmly attach to the bottom of the tube.
    - Add 1mL of ice-cold H50 and transfer cells to a 1.5mL tube.
    - Spin cells down (400xg, 2 min) and remove supernatant
    - Wash cells with 1mL of ice-cold H50 twice.
    - Resuspend cells in 1mL of ice-cold H50.
    - Make a 1/100 dilution and count cells.

[22] We typically use $2.5 \cdot 10^6$ cells for each transfection. See Figure S11A.

- Make aliquots of 1.25-5·$10^6$ cells for each transfection. The volume is not important, because cells will be pelleted again later. Keep on ice.
- Proceed with Step 4.3.

4.3. Prepare plates for growth after transfection.
- For analysis:
  - For non-axenic cells, prepare 24-well plates with 1mL *E. coli* B/r suspension ($OD_{600}$=1-2 in KK2-MC) per well for each transfection.
  - For axenic cells, prepare 24-well plates with 1mL HL5+FAB per well for each transfection.
- For sorting:
  - For non-axenic cells, prepare 6-well plates with 5mL *E. coli* B/r suspension ($OD_{600}$=1-2 in KK2-MC) per well for each transfection.
  - For axenic cells, prepare 6-well plates with 5mL HL5+FAB per well for each transfection.
- Keep on the side.

4.4. Prepare the RNP complex.
- Anneal tracrRNA and crRNA[23] by mixing the following components in a 0.2mL PCR tube to prepare the gRNA complex at a final concentration of 50µM. Scale up depending on the number of reactions.

| Reagent / solution | Amount/transfection |
|---|---|
| 100µM tracrRNA | 1.5µL |
| 100µM crRNA | 1.5µL |

- Heat at 95°C for 5 min.
- Cool down slowly to RT (-1°C/cycle, 20sec/cycle).
- The gRNA complex can be stored at -20°C up to 1 year.
- Assemble the RNP complex by mixing the following reagents at RT in the order specified by the table in a 1.5mL tube. Add the reagents very slowly, swirling the tip into the mix, and gently pipetting up and down once or twice after the addition of each reagent.

| Reagent / solution | Amount/transfection | Final concentration |
|---|---|---|
| ~15µg/µL (~90µM) SpyCas9[24] | 1.31µL | ~12µM |
| 50µM gRNA complex | 3µL | 15µM |
| Nuclease-free $H_2O$ | Up to 10µL | |

4.5. Transfection mix in electroporation buffer.
- Add the following reagents to the RNP mix as indicated.

[23] Pre-annealing of the crRNA and tracrRNA results in better efficiencies (Figure S4B), and therefore we recommend including this step. However, this step can be eliminated, and the crRNA and tracrRNA can be directly added to the Cas9 protein.
[24] The best results are obtained with ultrapure WT SpyCas9 (Figure S4B). However, good efficiencies are also obtained with a regular 1-day purification protocol (see Methods for both quick and extended purification protocols). TrueCut Cas9 Protein v2 (Thermo Fisher, A36498) is the best-performing commercially available SpyCas9 we tested (see Figure S4B).

| Reagent / solution | Amount/transfection with donor oligo | Amount/transfection with donor PCR product | Final concentration |
|---|---|---|---|
| RNP complex | 10µL | 10µL | |
| 100µM donor oligo | 0-4.8µL | NA | 0-4.8µM |
| 1µg/µL donor PCR product | NA | 0-20µL | 0-20µg/100µL[25] |
| 100µM EE (IDT)[26] | 0-1.2µL (optional) | 0-1.2µL (optional) | 0-1.2µM (optional) |
| H50 buffer | Up to 100µL | Up to 100µL | |

- Add H50 buffer to the RNP complex in small volume increments (for example, 15+25+40µL) to a final volume of 100µL.
- Combine cells and transfection mix[27].
    - Spin down aliquot of cells ($1.25\text{-}5\cdot10^6$ cells; 400xg, 2 min) and remove supernatant.
    - Resuspend cell pellet by adding 100µL of RNP complex and gently pipetting up and down once or twice.
    - Keep cell suspension with RNP complex on ice.

4.6. Electroporation.
- Pre-chill electroporation cuvettes (1mm gap, Bio-Rad).
- Transfer 80µL of each sample to ice-cold electroporation cuvette.
- Dry the sides of the cuvette with a kimwipe.
- Electroporate using a Gene Pulser Xcell Microbial System.
- Electroporation settings:
    - Exponential protocol
    - Voltage (V): 750
    - Capacitance (µF): 25
    - Resistance (Ohms): Infinite
    - Cuvette (mm) :1
    - Number of pulses: 1
- Place cuvette on ice immediately after electroporation.

4.7. Grow cells after electroporation.
- Let the electroporated cells stand on ice for 5 min.
- For analysis, transfer 10-20µL to a well of 24-well plate prepared in Step 4.3, containing:
    - For non-axenic cells, 1mL *E. coli* B/r suspension ($OD_{600}$=1-2 in KK2-MC) per well.
    - For axenic cells, 1mL HL5+FAB per well.
- For sorting, transfer 60-80µL to a well of a 6-well plate prepared in Step 4.3, containing:
    - For non-axenic cells, 5mL *E. coli* B/r suspension ($OD_{600}$=1-2 in KK2-MC) per well.
    - For axenic cells, 5mL HL5+FAB per well.
- Incubate cells for 1-3 days.
- Replace media every day.
- Proceed to analysis by flow cytometry or clone isolation by FACS sorting.

---

[25] In our example, where we knock-in *mNeonGreen-P2A* into *mCherry*, the donor PCR fragment is 899bp long. Therefore, 1µg of PCR product = 1.8pmols, and 1µg/100µL = 1.8µM.

[26] If desired, EE (from IDT) can be added to the transfection mix, however, we only observe a small improvement in knock-out efficiency (see Figure S6 and S7), and we generally do not recommend it, as it comes with the risk of inserting EE-derived DNA into the CRISPR-Cas9 target site.

[27] Note that for combinatorial knock-ins or knock-outs, independent transfection mixes can be prepared, and then mixed at a 1:1 ratio before adding them to the cells and proceeding with Step 4.6.

##### Step 5. Flow cytometry analysis.

*When generating a fluorescent knock-in mutant at the act5 locus, fluorescent cells can be detected a few hours after transfection. Flow cytometry can therefore be done within a day. When knocking out a fluorescent protein, like the mCherry gene in our example, loss of fluorescence intensity in cells is also visible within one day, but it becomes more apparent as the fluorescent protein is diluted out and degraded. We therefore typically analyze knock-out and knock-in efficiencies 2 days post-transfection.*

5.1. Prepare cells for flow cytometry analysis.

- Harvest transfected cells 1-3 days after transfection from 24-well plate.
    - Remove media with a cell culture vacuum aspirator (or a pipette).
    - Add 400µL KK2-MC buffer.
    - Harvest cells by gently pipetting them up and down.
    - Transfer 200µL of cells to a 5mL Falcon round-bottom tube.
    - Add 2µL DRAQ7.
    - Briefly vortex.
    - Incubate 5-10 min.

5.2. Analysis by flow cytometry.

- Analyze cells in BD FACSymphony A3 analyzer with BD FACSDiva software Version 9.1.2 (or another available flow cytometer).
    - For the FACSymphony, we use the following settings.
        - FSC: 100V
        - SSC: 150V
        - 488-530_30: 285V
        - 561-610_20: 434V
        - 640-670_30: 500V,
        - Flow rate lower than 500 cells/sec.
    - When possible, include both negative and positive controls for gating purposes.
- Visualize results with FlowJo or other available tools.
    - Gate cells using FSC and SSC channels. Particularly important for gating against bacterial cells when using non-axenic cultures.
    - Gate for living cells using 640-670_30. Living cells are impermeable to DRAQ7, and are therefore negative in the 640-670_30 channel.
    - Gate for singlets using FSC-A vs FSC-H.
    - Gate fluorescent and non-fluorescent cells based on 561-610_20 (mCherry-/mCherry+ cells) or 488-530_30 (mNeonGreen-/mNeonGreen+ cells) channels. Use controls to confirm appropriate gating.

**Step 6. Isolation of gene-edited cells using cell sorting.**

*When knocking-in a fluorescent protein under the control of a promoter that is expressed during growth (i.e., vegetative life stage), like the act5 gene, expression can be detected within a few hours after transfection. We therefore normally sort cells one or two days post-transfection. When knock-in mutants have a growth defect, sorting cells earlier might increase efficiency.*

6.1. Cell sorting.

- Prepare 24-well plates.
  - Add 1mL SM5-charcoal-agar[28] to each well using a multi-dispenser pipette[29] and let it solidify with the lid off for 20-30 min under a vertical flow hood. Plates can be stored at 4°C for up to 1 month.
  - On the day of sorting, add 25µL of *E. coli* B/r cell suspension ($OD_{600=}0.33$ in KK2-MC buffer) to each well. Spread the cells using a heat-bended glass Pasteur pipette.
  - Let it dry for 10-25 min.
- Harvest transfected cells from 6-well plate.
  - Remove media with a cell culture vacuum aspirator (or a pipette).
  - Add 5mL KK2-MC buffer with 2x pen/strep[30].
  - Remove buffer.
  - Add 500µL KK2-MC buffer with 2x pen/strep.
  - Harvest cells by gently pipetting up and down.
  - Collect cells in a 5mL round-bottom tube with a 35µm cell strainer snap cap.
- Sort individual cells onto the 24-well plate with SM5-charcoal-agar using the BD FACSAria Fusion Flow Cytometer sorter with BD FACSDiva software Version 8.0.1 (or another cell sorter).
  - For the FACSAria, we used the following settings:
    - FSC: 0V
    - SSC: 242V
    - 488-530_30: 362V
    - 561-610_20: 539V
    - 640-670_30: 466V,
    - Nozzle: 100µL,
    - Single cell sorting, target: 1 cell.
    - Flow rate lower than 400 cells/s.
  - When possible, also sort positive and negative controls.
- Incubate plates at 22°C for 3 to 5 days.

6.2. Clone confirmation by imaging

- When generating fluorescent knock-ins or non-fluorescent knock-outs, image the 24-well plate using a fluorescent stereomicroscope[31].
- Positive clones can also be confirmed through PCR.

---

[28] We add charcoal to the SM5-agar media to have black growth media and reduce autofluorescence of the media during imaging of the plates to identify positive clones by fluorescence microscopy (see below).
[29] Adding 1mL of agar-media with a multi-dispenser pipette, such as Eppendorf's Multipette E3x**,** ensures adding the exact same volume to each well, and therefore makes imaging easier, as the same focal plane can be used for imaging all wells.
[30] Antibiotics can be avoided if bacteria contamination of the sorter is not an issue.
[31] We recommend the Zeiss Axio Zoom.V16 fluorescent stereomicroscope.

6.3. Clone confirmation by PCR

- Isolate genomic DNA from fruiting bodies:
  - Prepare a PCR strip, and add 20μL genomic DNA extraction buffer to each tube. Scale up reaction depending on the number of samples.

| Reagent / solution | Volume |
|---|---|
| 10x Taq buffer | 1μL |
| IGEPAL | 0.1μL |
| 20mg/mL Proteinase K | 0.05μL |
| Nuclease-free $H_2O$ | 18.85μL |

  - Harvest 1-3 sori per well using a 10μL pipette tip.
  - Dip the tip in the genomic DNA extraction buffer.
  - Incubate at 56°C for 45 min, and then at 95°C for 10 min.
  - gDNA extract can be stored at 4°C up to 3 days, but best results are obtained when used immediately.

- PCR reaction:
  - Design primers flanking the targeted region. Aim for PCR products between 500 and 1000bp. For long inserts, two primer sets can be designed, spanning each junction. In the following example we use MR101/MR113 for the upstream junction, and MR112/MR103 for the downstream junction.

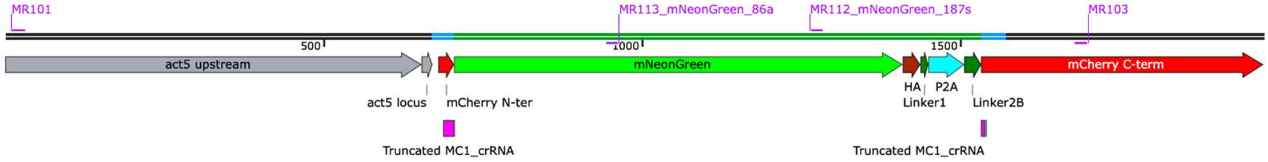

**act5::mNeonGreen-P2A-mCherry**

  - Use 2μL of genomic DNA from previous step to prepare the PCR reaction.

| Reagent / solution | Volume |
|---|---|
| Taq 2X Master Mix | 10μL |
| 10μM Forward primers | 0,4μL |
| 10μM Reverse primers | 0,4μL |
| Nuclease-Free $H_2O$ | 7,2μL |
| | 18μL |
| Quick Genomic DNA extract | 2μL |

  - Perform confirmation PCR using the following settings:

| Step | Temperature | Time |
|---|---|---|
| Initial Denaturation | 95°C | 2 min |
| Amplification: 35 cycles | 95°C | 30 sec |
| | $T_a$[32] | 20 sec |
| | 68°C | 1 min |
| Final Extension | 68°C | 10 min |
| Hold | 4–10°C | |

  - Check PCR results on 1% agarose gel as described before (Step 2.4).

[32] $T_a$ might have to be adjusted depending on the primers and the amplified sequence. We successfully use Ta=56°C for all the confirmation PCRs with the primer sets in Table S6.

6.4. Sanger sequencing

- Clean the remaining PCR product (18µL) with a Qiagen PCR purification column following manufacturer's instructions. Elute in 25µL EB and check purity with a NanoDrop Spectrophotometer.
- Adjust the concentration of the purified DNA following the recommendations of the sequencing company and send the samples for sequencing with one or both primers.

**Step 7. Propagation of positive clones.**

Clones confirmed by fluorescent microscopy and PCR can be stocked.

7.1. Propagate cells.

- Prepare normal SM5-agar plates
- Spread 200µL of *E. coli* B/r in KK2 buffer at $OD_{600}$=2 and let it dry for 5 min.
- Pick one sorus from a positive well using a pipette tip.
- Spot this on the bacterial lawn.
- Incubate plates at 22ºC for 3-7 days until the fruiting bodies cover the whole plate.

7.2. Optional: confirm purity of clone by flow cytometry analysis.

- Harvest some cells from the feeding front with a loop in 1mL KK2.
- Analyze by flow cytometry as detailed in Step 5.2[33].

7.3. Make glycerol stocks.

- Harvest sori using a 10µL loop
- Dip loop in 1mL of 25% glycerol/KK2-MC buffer.
- Store at -80ºC.

[33] Expression of the fluorescent reporter will depend on the expression pattern of the targeted gene.

#### Part 2. Notes for CRISPR-Cas9 editing in other *Dictyostelid* species

Our CRISPR-Cas9 editing protocol can also be applied to other *Dictyostelid* species, but comes with several small adjustments. Below we specify these adjustments for each of the seven steps in our protocol.

**Step 1.** Identifying good CRISPR-Cas9 targets.

- Many *Dictyostelid* genomes are not available for popular tools like Cas-designer. When targeting a *Dictyostelid* genome different to *D. discoideum*, a custom tool for identifying CRISPR-Cas9 targets can be used. Alternatively, a conventional tool (e.g., Cas-designer) with the *D. discoideum* reference can be used. When using a conventional tool, search for potential off-target hits, for instance by Blasting your spacer and PAM sequence against the *Dictyostelid* genome of interest. Detecting all possible off-target hits for spacers with one or two nucleotide mismatches can be easily automated in Python.

**Step 2.** Designing donor oligos and donor PCR products.

- Same as for *D. discoideum.*

**Step 3.** Culturing *Dictyotelid*s.

- Most *Dictyostelid* species can be grown following the same protocol as described for non-axenic culturing of *D. discoideum*. Sometimes slightly different media conditions are recommended (e.g., LP-agar or SM5-agar with charcoal). Dictybase provides useful information there.

**Step 4.** Transfecting cells with RNP complex by electroporation.

- Cells were prepared and transfected following the same protocol as for non-axenic *D. discoideum*. Although we have not tested this, we can imagine that optimal electroporation conditions might sometimes differ between species. Growth rates on SM5-agar plates also differ between species, and therefore the appearance of a clear feeding front might take a different number of days: for example, for *T. lacteum*, it typically takes 5 to 7 days to get a clear feeding front, while for *D. purpureum*, it only takes 1 or 2 days and the plate is overgrown in 5 days. It is therefore important to account for such growth differences when transfecting multiple species in parallel.

**Step 5.** Flow cytometry analysis.

- The same parameters were used for all species. However, the gating slightly varied depending on the size/granularity of cells and the brightness of fluorescent reporters in the different species.

**Step 6.** Isolation of gene-edited cells using cell sorting.

- The same parameters were used for all species. However, the gating slightly varied depending on the size/granularity of cells and the brightness of fluorescent reporters in the different species.

**Step 7.** Propagation of positive clones.

- Same as for *D. discoideum.*

**References**

Bae, S., J. Kweon, H. S. Kim, and J. S. Kim. 2014*a*. Microhomology-based choice of Cas9 nuclease target sites. Nature Methods 11:705–706.

Bae, S., J. Park, and J. S. Kim. 2014*b*. Cas-OFFinder: a fast and versatile algorithm that searches for potential off-target sites of Cas9 RNA-guided endonucleases. Bioinformatics 30:1473–1475.

Concordet, J. P., and M. Haeussler. 2018. CRISPOR: intuitive guide selection for CRISPR/Cas9 genome editing experiments and screens. Nucleic Acids Research 46:W242–W245.

Paix, A., A. Folkmann, D. H. Goldman, H. Kulaga, M. J. Grzelak, D. Rasoloson, S. Paidemarry, et al. 2017. Precision genome editing using synthesis-dependent repair of Cas9-induced DNA breaks. Proceedings of the National Academy of Sciences 114:E10745–E10754.

Paix, A., D. Rasoloson, A. Folkmann, and G. Seydoux. 2019. Rapid tagging of human proteins with fluorescent reporters by genome engineering using double-stranded DNA donors. Current Protocols in Molecular Biology 129:e102.

Park, J., S. Bae, and J. S. Kim. 2015. Cas-Designer: a web-based tool for choice of CRISPR-Cas9 target sites. Bioinformatics 31:4014–4016.

Paschke, P., D. A. Knecht, A. Silale, D. Traynor, T. D. Williams, P. A. Thomason, R. H. Insall, et al. 2018. Rapid and efficient genetic engineering of both wild type and axenic strains of *Dictyostelium discoideum*. PLoS One 13:e0196809.

Sheikh, S., M. Thulin, J. C. Cavender, R. Escalante, S. I. Kawakami, C. Lado, J. C. Landolt, et al. 2018. A new classification of the Dictyostelids. Protist 169:1–28.

Zhu, X., C. Ricci-Tam, E. R. Hager, and A. E. Sgro. 2023. Self-cleaving peptides for expression of multiple genes in *Dictyostelium discoideum*. PLoS One 18:e0281211.
